# UNCG RNA tetraloop as a formidable force-field challenge for MD simulations

**DOI:** 10.1101/2020.07.27.223826

**Authors:** Klaudia Mráziková, Vojtěch Mlýnský, Petra Kührová, Pavlína Pokorná, Holger Kruse, Miroslav Krepl, Michal Otyepka, Pavel Banáš, Jiří šponer

## Abstract

Explicit solvent atomistic molecular dynamics (MD) simulations represent an established technique to study structural dynamics of RNA molecules and an important complement for diverse experimental methods. However, performance of molecular mechanical (MM) force fields (*ff*s) remains far from satisfactory even after decades of development, as apparent from a problematic structural description of some important RNA motifs. Actually, some of the smallest RNA molecules belong to the most challenging systems for MD simulations and, among them, the UNCG tetraloop is saliently difficult. We report a detailed analysis of UNCG MD simulations, depicting the sequence of events leading to the loss of the UNCG native state during MD simulations. We identify molecular interactions, backbone conformations and substates that are involved in the process. The total amount of MD simulation data analyzed in this work is close to 1.3 millisecond. Then, we unravel specific *ff* deficiencies using diverse quantum mechanical/molecular mechanical (QM/MM) and QM calculations. Comparison between the MM and QM methods shows discrepancies in the description of the 5’-flanking phosphate moiety and both signature sugar-base interactions. Our work indicates that poor behavior of the UNCG tetraloop in simulations is a complex issue that cannot be attributed to one dominant and straightforwardly correctable factor. Instead, there is a concerted effect of multiple *ff* inaccuracies that are coupled and amplifying each other. We attempted to improve the simulation behavior by some carefully-tailored interventions but the results are still far from satisfactory, underlying the difficulties in development of accurate nucleic acids *ff*s.

## INTRODUCTION

Explicit solvent atomistic molecular dynamics (MD) simulations represent an established tool to study structural dynamics of RNA molecules.^1–3^ MD simulations can complement diverse experimental methods by, e.g., allowing to go beyond the typical static (ensemble-averaged) description of RNA molecules obtained by structural biology methods. However, the reliability of MD predictions depends on quality of the used molecular mechanical (MM) force fields (*ff*s). Unfortunately, performance of these *ff*s remains far from satisfactory despite ongoing development, and is also very system-dependent.^4,5^ Many RNA molecules and protein/RNA complexes are quite well described in MD simulations, at least when starting the simulations from experimental structures. However, other systems progressively deteriorate upon extending the simulations. Surprisingly, even small RNA molecules may present persisting challenges for MD simulations and among them, the UNCG tetraloop is saliently difficult.^6–11^

RNA Tetraloops (TLs) are stem-loop RNA structures, which are common secondary structure elements in RNA. TLs participate in tertiary folding, RNA-RNA and protein-RNA interactions, ligand binding, and other important biological functions.^12,13^ The UNCG TL (N stands for any nucleotide) is one of the two most abundant TLs.^14–16^ It is thermodynamically very stable and possesses a clearly-defined dominant folded topology stabilized by a set of signature molecular interactions (**Figure 1**). The native state of UNCG TL remains difficult to be described by the current RNA *ffs*. Recent folding simulations of the UNCG TL indicated a large free-energy imbalance between native (folded) and other states (misfolded/unfolded conformations).^6–10,17–20^ In addition, the characteristic UNCG native structure is lost in sufficiently long standard simulations. Two different effects were deemed to dominantly lead to the incorrect folded/unfolded free-energy balance, namely, (i) excessive stabilization of the unfolded ssRNA structure by intramolecular base-phosphate and sugar-phosphate interactions and (ii) destabilization of the native folded state by underestimation of the native H-bonds including the stem base pairing.^8,9,21,22^ Recently, we introduced a general external potential tuning H-bond interactions (gHBfix),^8,19^ which is used as an additional *ff* term to improve RNA simulations. Even with this modification, description of the UNCG TL remained imbalanced and we were not capable to fold the 8-mer TL. Despite occasional literature claims reporting success in UNCG simulations in the past (not unambiguously confirmed by other groups;^8,9^ reviewed in ref. 4), we are not aware of any existing RNA *ff* able to properly describe UNCG TL with all its signature interactions.^19,23^ Some success has been recently achieved with the DESRES *ff* with specific ion parameters using the UNCG 14-mer with longer A-RNA stem.^24^ However, the same *ff* leads to a complete disruption (including secondary structure rearrangements) of some other important RNA systems even in standard simulations on sub-microsecond timescale.^19,23^ Such side-effects in simulations of functional RNA molecules outweigh partial improvements for small models.^19,23^ Thus, the UNCG *ff* issue remains unresolved.

**Figure 1.**
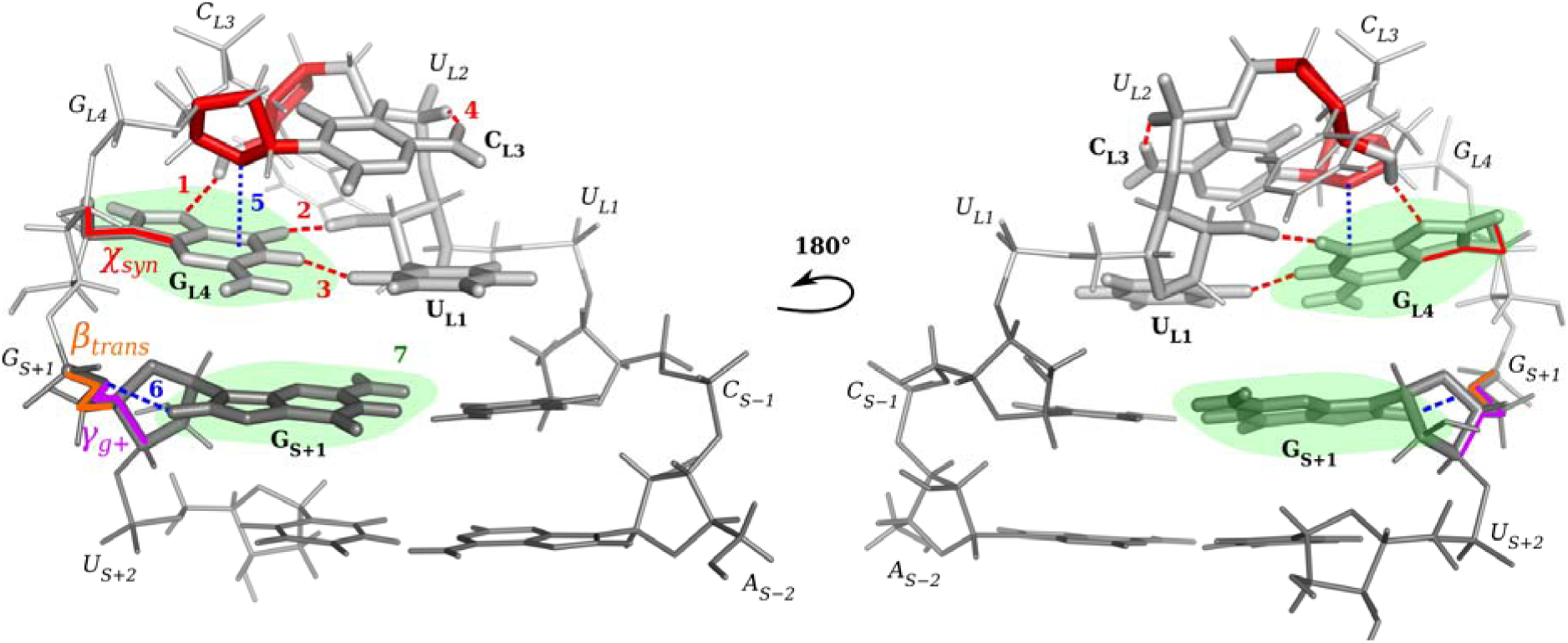
Native structure of the UNCG TL. Nucleotides of the loop are labelled as U_L1_, U_L2_, C_L3_ and G_L4_, and stem nucleotides are labeled as A_S−2_, C_S−1_, G_S+1_ and U_S+2_. Phosphates and bases are identified by normal and bold font, respectively. We note that r(acUUCGgu) and r(gcUUCGgc) 8-mers were studied in this work. Signature features of UNCG TL are colored in red and include the U_L2_(2’-OH)…G_L4_(N7) H-bond (marked as “1” in the Figure), U_L1_(2’-OH)…G_L4_(O6) H-bond (2), U_L1_(O2)…G_L4_(N1) H-bond (3), C_L3_(N4H)…U_L1_(*pro*-R_P_) H-bond (7BPh interaction^29^) (4), *syn*-conformation of G_L4_ nucleotide (*χ*_*syn*_) and C2’-*endo* pucker of U_L2_ and C_L3_ ribose. Other key structural features that were investigated in this work include: the C_L3_/G_L4_ ribose-base stacking (5), G_S+1_(C8H8)…G_S+1_(O5’) 0BPh interaction (6), mutual position of G_L4_ and G_S+1_ nucleobases (7), and *β*_*trans*_ (orange) and *γ*_*g*+_ (violet) conformations of G_S+1_ nucleotide.

In the present work, we analyze origins of the notoriously problematic description of the UNCG TL. First, we carry out a detailed analysis of our earlier and new MD simulations and obtain a detailed picture of the sequence of events that leads to loss of the native state. We identify molecular interactions, backbone substates and other conformational features that are involved in the process. Then, we try to unravel specific *ff* deficiencies using massive quantum mechanical/molecular mechanical (QM/MM)^25^ and QM calculations. Obviously, as discussed elsewhere,^26–28^ QM and QM/MM calculations cannot replace MD simulations due to the lack of Boltzmann sampling. There is generally a limited transferability between calculations of potential-energy surfaces (gradient optimizations or energy scans) as done by QM methods and free energies that determine populations in MD simulations. Still, careful comparison of accurate-enough QM and QM/MM calculations with equivalent *ff* data provides substantial insights into the imbalances that are inherent to *ff*s. This is the goal of our QM/MM and QM computations. Finally, we try to improve the simulation behavior by some carefully-tailored interventions. Our work clearly indicates that poor behavior of the UNCG simulations is a complex issue and cannot be explained by one single dominant factor that would be straightforwardly correctable. Instead, there is a concerted effect of multiple *ff* inaccuracies which are coupled and magnifying each other’s effect. Such accumulation of diverse inaccuracies in description of a short UNCG RNA sequence that in addition folds into a very stiff structure provides no conformational freedom to buffer the individual imbalances. It leads, even for moderate imbalances, to substantial structural strains.

## METHODS

### Starting structures

The initial coordinates were taken either from high-resolution NMR structure of the r(ggcacUUCGgugcc) 14-mer (PDB ID 2KOC^30^) or excised from the 2.8 Å-resolution X-ray structure (PDB ID 1F7Y^31^) as the r(gcUUCGgc) 8-mer. The starting topologies and coordinates for MD simulations were prepared using the tLEaP module of AMBER 16 program package.^32^ Structure of the isolated small TL motif in solution is in dynamic temperature-dependent equilibrium between folded and unfolded conformations. Most of the calculations were done using 8-mer oligonucleotides, resulting, upon correct folding, into a two-base-pair stem and the TL itself. As TLs with longer stems require higher temperature for melting (unfolding),^33,34^ minimal 8-nucleotide long (8-mers) TL motifs with just two base pairs are commonly used in computational studies.^6–10,18,19^ There is a clear experimental evidence that even these short oligomers should be dominantly in the folded state,^35^ validating the 8-mer calculations.

### Classical MD simulations

We used the standard *ff*99bsc0*χ*_OL3_^36–39^ AMBER RNA *ff* with the van der Waals (vdW) modification of phosphate oxygens developed by Steinbrecher et al.,^40^ where the affected dihedrals were adjusted as described elsewhere.^22^ This RNA *ff* is abbreviated as *χ*_OL3CP_ henceforth and libraries can be found in the Supporting Information of ref. 8. In the simulations we in addition applied our gHBfix^19^ H-bonding potential that was shown to improve overall performance of the RNA *ff*. In most cases we used its basic version from ref. 19, abbreviated subsequently as gHBfix19 in ref. ^41^. When any modified alternative is used, it is always specified in the text. Majority of simulations were solvated with the OPC^42^ water model and in ∼0.15 M KCl using the Joung−Cheatham^43^ ion parameters. Specific tests involved TIP3P^44^ and SPC/E^45^ water models and were also performed in Na^+^ net-neutral and 1.0 M KCl salt-excess ion environments. Other details about the MD protocols can be found in Section S1 in the Supporting Information. We have used also simulations from our preceding papers.^19,23^

### REST2 Settings

The replica exchange solute tempering (REST2)^46^ simulations of the r(gcUUCGgc) TL were performed at 298 K with either 12 or 16 replicas. Details about settings can be found elsewhere.^8^ The scaling factor (λ) values ranged from 1 to 0.59984 and from 1.0454 to 0.59984 for 12 and 16 replicas, respectively. Those values were chosen to maintain an exchange rate above 20%. The effective solute temperature ranged either from 298 K (12 replicas) or 285 K (16 replicas) to ∼500 K. The hydrogen mass repartitioning^47^ with a 4-fs integration time step was used.

### QM/MM, QM/COSMO, MM and MM/GB geometry optimizations of UNCG TL conformations

We have carried out a series of gradient geometry optimizations in order to compare the MM (i.e., *ff*) description with advanced QM methods. The initial r(acUUCGgu) and r(gcUUCGgc) 8-mers were either taken from eight different MD snapshots (structures **1**-**8**) or excised from the NMR structure of the r(ggcacUUCGgugcc) 14-mer (PDB ID 2KOC, structure **9**). The counter-ions (K^+^) were added using the tLEaP module of AMBER 16 to neutralize the RNA. SPC/E water model sphere with radius ∼40 Å from the geometrical center of the RNA was prepared using tLEaP for all the initial structures. Short water equilibrations (< 10 ps MD) of the added water molecules were performed before geometry optimizations for structures **1, 2, 3** and **4**. In contrast, five different water distributions were used for geometry optimizations of structures **5, 6, 7, 8** and **9** obtained by 20 ps, 40 ps, 60 ps, 80 ps, and 100 ps-long solvent equilibrations. The resulting 25 starting structures are denoted as **5a**-**e, 6a-e, 7a-e, 8a-e** and **9a-e**. Water distribution may influence the relaxed solute structures as shown in ref. 26. For the sake of simplicity, only results for structures after the 100 ps-long solvent equilibration (i.e., structures **5-9e**) are documented in the main text while the full set of data can be found in Supporting Information.

The additive QM/MM scheme with point-charge approximation for electrostatic embedding was used for all QM/MM calculations. The QM/MM module of AMBER 14^48,49^ was coupled with TURBOMOLE V7.3^50,51^ using an in-house modified version of Sander^26,52,53^ from the AMBER 14 program package. For geometry optimization the limited-memory Broyden-Fletcher-Goldfarb-Shanno quasi-Newton algorithm (L-BFGS)^54^ was used with a convergence threshold of 1 e^−4^ kcal·mol^−1^·Å^−1^ for the gradient norm. Turbomole was used to calculate energies and gradients of the QM region using the composite PBEh-3c^55^ hybrid-DFT method with resolution-of-identity (RI) approximation for the Coulomb integrals.^56^ The PBEh-3c method employs the double-zeta valence polarized def2-mSVP basis set, the empirical DFT-D3 dispersion correction^57^ with Becke-Johnson (BJ) damping^58^ and the geometrical counter-poise (gCP) correction for intra- and intermolecular basis set superposition error (BSSE).^59^ The QM region included the whole RNA 8-mer while the MM region consisted of water molecules and counter-ions.

Similarly to the work by Pokorna et al.,^26^ QM/MM optimizations were mirrored by equivalent MM optimizations from identical starting structures. For the sake of completeness we note that the *ff*99bsc0*χ*_OL3_ was used for structures **1-5** and **9** and *χ*_OL3CP_ for structures **6**-**8**; the *ff* difference is, however, marginal for optimizations (see **Figure S1** in the Supporting Information). Geometrical parameters (backbone dihedrals, intramolecular distances and signature H-bonds) of the QM/MM- and MM-optimized TLs were analyzed using the CPPTRAJ^60^ module of AMBER 16. Mutual orientation of nucleobase planes, i.e., angle between planes of the G_L4_ *syn* base and of the closing G_S+1_ base, and sugar-base stacking interaction were analyzed using the Geom_util program (https://github.com/hokru/geom_util).

Besides MM optimizations from the initial structures, we also performed MM re-optimizations starting from the QM/MM-optimized structures to reduce potential uncertainty due to comparison of too distinct local minima, i.e., a situation where the QM/MM and MM optimizations from the MD snapshot diverge to non-equivalent (too different) local minima.^26^ The QM/MM to MM re-optimized geometry should typically stay close to the QM/MM-optimized geometry (see the Supporting Information). Therefore, comparing geometrical parameters between QM/MM-optimized and QM/MM to MM re-optimized geometries should conservatively show differences between QM and MM potential energy surfaces (PES), although the re-optimization may also mask MM imbalances by excessively attenuating the rearrangements, for detailed discussion see ref. 26.

RNA 8-mers were additionally optimized in implicit solvent. The Conductor-like screening model (COSMO)^61^ and the Generalized Born model (GB)^62,63^ were used for QM and MM, respectively, with dielectric constant *ε* of 78.5.

### Relaxed U_L1_(2’-OH) dihedral scans of RNA TLs

In order to analyze quality of the MM description of orientation of the key U_L1_(2’-OH) group, we compared QM and MM relaxed conformational energy scans of the U_L1_(C1’-C2’-O2’-O2’H) dihedral for structures **2, 7, 8** and **9**.

Firstly, the ONIOM scheme^64^ was used for MM optimizations of the UNCG TL 8-mer (the high-layer) and solvent (the low-layer) using Gaussian 09.^65^ As the same MM method (*χ*_OL3CP_ RNA *ff*) was used for both layers the energy/gradient components of the high-layer calculations effectively cancel out. This approach results in MM geometry optimization with constraints using the microiterations approximation where water molecules are relaxed after each dihedral angle change. This is a technical workaround to enable efficient internal coordinate quasi-Newton optimizations necessary for constraints for an otherwise prohibitively large molecule. We mark this approach as MM/MM optimization throughout the paper.

To save computer time, the MM/MM-optimized geometries were used to initiate the corresponding QM/MM optimizations. The AMBER 14 coupled to Turbomole was used for QM/MM optimizations applying tight geometry restraints on the U_L1_(C1’-C2’-O2’-O2’H) dihedral (see the Supporting Information for details). The technical difference between using geometry constraints and restraints has no practical impact on the results.

### QM and MM calculations of small model systems

We have also carried out calculations on various small model systems. We extracted specific structural moieties of the UNCG TL (see Section S2 in the Supporting Information for the list of studied small models) for comparative QM and MM analyses, i.e., local geometry optimizations, interaction energy scans and conformational energy scans. The *Xopt*^66,67^ tool coupled with Turbomole was used for all the QM geometry optimizations performed at the PBEh-3c level using the RI approximation. Turbomole was also used for the rigid-monomer QM energy scans performed at the same level of theory. ORCA V4.1.0^68,69^ was used for the reference (highest-quality) QM calculations performed at the DLPNO-CCSD(T)/TightPNO/CBS level using aug-cc-pV(T-Q)Z and corresponding auxiliary aug-cc-pVQZ/C basis sets, and with TightSCF setting.^70–74^ MM geometry optimizations were performed using *Xopt* coupled with AMBER 14 or using Gaussian 09. QM and MM optimizations and single-point energy calculations of dinucleotide monophosphate models were performed with COSMO and GB implicit solvents (*ε* = 78.5). For rigid-monomers MM interaction energy scans, we used the development version of the *bff* program^75^ using non-bonded energy terms of the AMBER *ff*. For the energy scans of small models, the MM method is denoted as AMBER *ff* and details are specified in the Supporting Information. Gaussian 09 was used for the MM relaxed conformational energy scan of the O3’-methylated-guanosine-monophosphate model. For additional computational details about calculations of the small models see Section S3 in the Supporting Information.

We used programs Molden,^76^ VMD^77^ and PyMOL^78^ for visualizing and analyzing the structures.

## RESULTS AND DISCUSSION

We applied diverse computational techniques, namely standard MD simulations, enhanced sampling REST2 simulations, QM, MM and multiscale QM/MM calculations (see Methods) in order to understand limitations of current state-of-the-art AMBER RNA *ff*s in description of the UNCG TL. Firstly, we analyzed disruption (unfolding) paths from standard as well as enhanced sampling MD simulations. We identified common features that seem to be responsible for instability of the native topology, i.e., specific structural reconformations that precede the collapse of the loop. We then performed various QM/MM and MM calculations of the UNCG TL to analyze differences in backbone dihedrals, signature H-bonds and other structural features, as described by QM and MM, in order to assess potential *ff* problems. Furthermore, we applied QM and MM calculations to small model systems to study interactions that may be triggering structural changes within the loop. Finally, we proposed certain fixes, namely (i) modulation of the key 0BPh interaction, (ii) adjustment of the sugar-base signature H-bonds and (iii) reduction of the vdW radii of non-polar hydrogens, which were subsequently tested during series of enhanced sampling MD simulations. In the text we first describe in detail the loss of the TL’s native structure in MD simulations and identify the individual potentially problematic energy terms. Then, we present the specific investigations visualizing and quantifying the *ff* limitations with aid of the QM methods. Finally, we describe modifications of the *ff* and their tests in simulations.

### Anatomy of disruption of the UNCG TL in simulations: a progressive loss of several interactions

We inspected specific parts of trajectories from standard as well as enhanced sampling MD simulations of the UNCG TL from our previous work,^19^ where signature interactions within the UNCG TL are lost and G_L4_ is repelled from its native conformation (**Figure 2**). Interestingly, disruption paths using multiple versions of the AMBER RNA *ff*^19,36,38,39^ have common features and could be characterized as an interplay of several factors. Those factors involve interactions that are, apparently, wrongly described with current RNA *ff*s.

**Figure 2.**
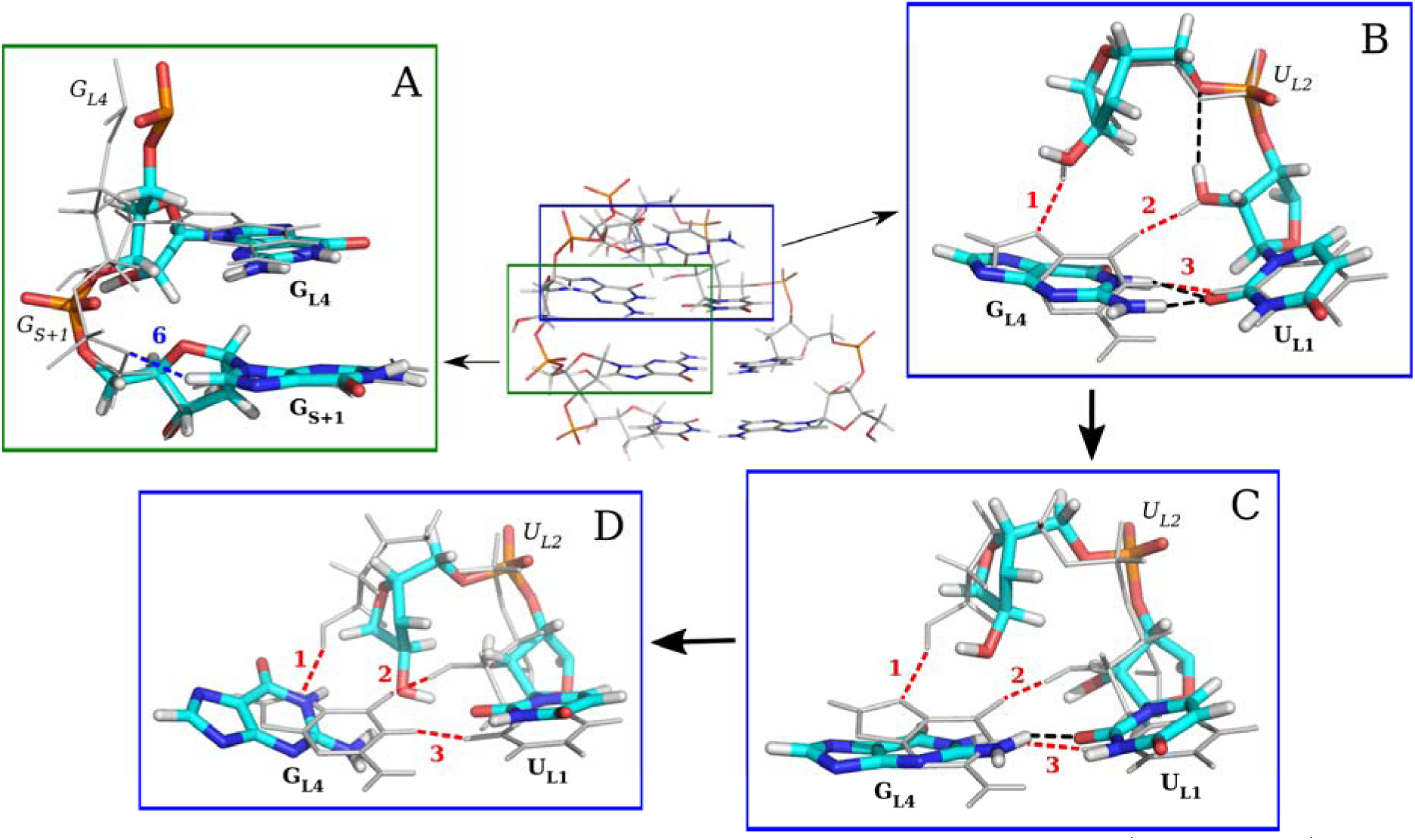
Disruption of the UNCG TL. The native structure of the 8-mer r(acUUCGgu) TL is shown in the middle with two important regions: (i) G_L4_-G_S+1_ dinucleotide moiety (green) and (ii) tetranucleotide loop (blue). Insets **A**-**D** show only the specific fragments, where details about loss of important interactions or key conformational characteristics are highlighted. The overlaid native structure is depicted by grey lines in each of the insets. The important interactions 1-3 and 6 of the native UNCG TL are marked in a similar way as in **Figure 1** for each of the insets. The disruption pathway of the UNCG TL is depicted by insets **B, C** and **D** and its flow is indicated by arrows. All the interactions and nucleotides shown by the insets are labelled similarly to **Figure 1**. (**A)** G_L4_-G_S+1_ dinucleotide with a flipped G_S+1_ phosphate accompanied by loss of the 0BPh interaction (6). (**B**) Part of the loop region, where the U_L1_(2’-OH) group is oriented towards the U_L2_ phosphate forming the spurious U_L1_(2’-OH)…U_L2_(O5’) H-bond. The partial twist of the G_L4_ nucleobase leads to formation of the bifurcated G_L4_(N1H/N2H)…U_L1_(O2) H-bond. Signature H-bonds 1 and 2 (see **Figure 1**) are lost; the U_L1_(2’-OH)…U_L2_(O5’) and G_L4_(N1H/N2H)…U_L1_(O2) H-bonds are depicted by black dashed lines. (**C)** Signature H-bonds (1-3) are lost and G_L4_ is forming only the spurious G_L4_(N2H)…U_L1_(O2) H-bond (black dashed line). (**D**) G_L4_ is further repelled from the loop and is not stabilized by any H-bond (bulge-out state). The common disruption pathway seen in simulation is documented as a movie in the Supporting Information.

First important factor is conformation of the G_S+1_ phosphate that is forming (in its native state) a 0BPh interaction^29^ (**Figure 2A**). The sugar-phosphate backbone around the G_S+1_ phosphate has a specific native conformation (see below) that is characterized by *β* and *γ* dihedrals in *trans* and *gauche*+ *(g*+) orientations, respectively (**Figure 3**). The *ff*s rather prefer G_S+1_ phosphate in an alternative, i.e., flipped state, characterized by *β*_*g*+_ and *γ*_*trans*_ dihedrals. However, counterintuitively, disruptions of the UNCG TL are initiated from the native state of the G_S+1_ phosphate and not from the alternative (likely spurious) flipped conformation. We suggest that occurrence of the alternative backbone state is a consequence of structural strain in the native structure (in the *ff* description) which is reduced either by adopting the flipped backbone conformation, or by onset of the overall TL structure disruption.

**Figure 3.**
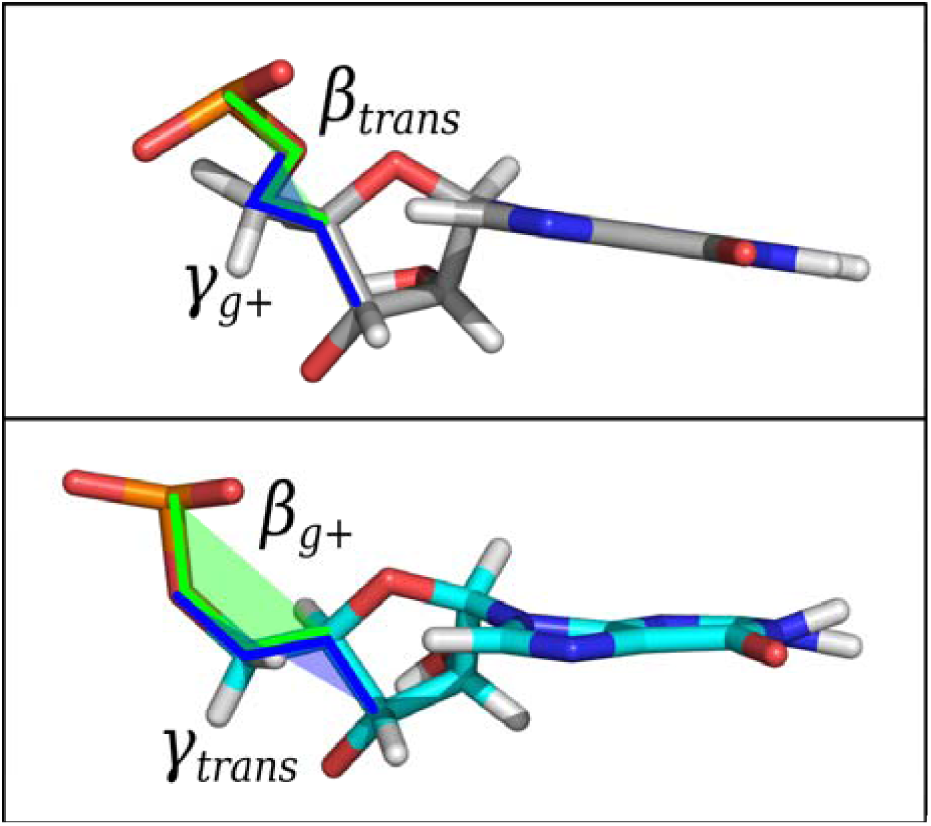
G_S+1_ phosphate in its native topology characterized by *β* and *γ* dihedrals in *trans* and *gauche*+ (*g*+) orientations, respectively (top), and in the flipped state characterized by *β* and *γ* dihedrals in *g*+ and *trans* orientations, respectively (bottom).

The other structural factors are the underestimated stability and subsequent loss of both signature sugar-base H-bonds within the loop, i.e., the U_L1_(2’-OH)…G_L4_(O6) and U_L2_(2’-OH)…G_L4_(N7) H-bonds. The 2’-OH groups in both H-bonds form alternative transient H-bonds during standard MD simulations. The former H-bond is reversibly lost due to the flip of U_L1_(2’-OH) towards the U_L2_ phosphate,^21^ so that a transient U_L1_(2’-OH)…U_L2_(O5’) H-bond is established (**Figure 2B**). The U_L2_(2’-OH)…G_L4_(N7) H-bond is also reversibly lost and the U_L2_(2’-OH) is then either pointing towards the solvent or alternative U_L2_(2’-OH)…G_L4_(O6) H-bond is formed. Importantly, formation of the alternative U_L2_(2’-OH)…G_L4_(O6) H-bond is not in disagreement with the experiment (see below).^30^ We further realized that loss of the U_L1_(2’-OH)…G_L4_(O6) H-bond is accompanied with structural rearrangement of the G_L4_ nucleobase. G_L4_ nucleotide (in its *syn* state) forms the GU wobble base pair with the U_L1_ nucleotide. Once the U_L1_(2’-OH)…G_L4_(O6) H-bond is lost, G_L4_ makes a partial twist within the loop, where the signature base – base G_L4_(N1H)…U_L1_(O2) H-bond is weakened and a bifurcated G_L4_(N1H/N2H)…U_L1_(O2) H-bond is established instead (**Figure 2B**). This transient state is frequently observed in simulations though the signature U_L1_(2’-OH)…G_L4_(O6) H-bond is typically reformed, followed by return of the G_L4_ nucleobase back to its canonical conformation. However, when the second sugar-base H-bond is lost simultaneously, i.e., neither the signature U_L2_(2’-OH)…G_L4_(N7) H-bond nor the alternative U_L2_(2’-OH)…G_L4_(O6) H-bond are formed, G_L4_ has an increased tendency to flip out of the loop (**Figure 2C**). Consequently, another structural feature, i.e., sugar-base stacking between the G_L4_ nucleotide and the O4’ atom of the C_L3_ ribose, is lost. In other words, the simultaneous loss of the signature interactions formed by both U_L1_(2’-OH) and U_L2_(2’-OH) groups opens the path for further destabilization of the structure, i.e., flip out of the G_L4_. As mentioned above, the scenario when G_L4_ flips out of the loop requires the G_S+1_ phosphate in its native state.

Once the G_L4_ is oriented towards the solvent (bulge-out state,^6,79^ **Figure 2D**) it typically remains there, i.e., the disruption pathway appears to be irreversible within typical timescales of standard MD simulations (several μs). Nonetheless, recently published 10 μs-long standard MD simulations with the *χ*_OL3CP_ + gHBfix19 *ff* version revealed occasional return of the G_L4_ nucleotide back from the bulge-out state to its native conformation within the loop.^19^ We identified the same flip-out flip-in pathway also in simulations with the DESRES^24^ RNA *ff*.^23^ Thus, the results indicate existence of a rather narrow conformational corridor (bottleneck) for expulsion and insertion of the G_L4_. It is worth noting that the tendency of expelling G_L4_ out of the loop is higher when another signature interaction within the loop, i.e., the C_L3_(N4H)…U_L1_(*pro*-R_P_) base-phosphate (BPh) interaction type 7 (7BPh)^29^ is weakened.

### Relationship between the G_S+1_ phosphate conformation and the 0BPh interaction

As noted above, in simulations of folded UNCG TL the G_S+1_ phosphate fluctuates frequently between two substates. The G_S+1_ phosphate flipping is mainly characterized by the shift of *β*_*trans*_*/γ*_*g*+_ dihedrals towards alternative *β*_*g*+_*/γ*_*trans*_ conformations (**Figure 3**). The flips can be monitored by the RNA suitename code^80^ as they change backbone conformations (RNA suites) from the canonical **1a** towards unusual **1e, 1c**, or **unidentified** RNA suites. Before the loss of native UNCG TL topology the G_S+1_ phosphate flipping is entirely reversible. In these initial parts of simulations the alternative flipped state is prevalent (∼70% population, **Figure S2** and **Figure S3** in the Supporting Information)^19^ and is accompanied by loss of the G_S+1_(C8H8)…G_S+1_(O5’) 0BPh interaction (**Figure 2A**).

0BPh is a weak C-H…O H-bond established by the C6-H6/C8-H8 groups of the pyrimidine/purine nucleotides which in most RNAs typically involves bridging O5’ phosphate oxygen from the same residue).^29^ In **1a** suite conformation of canonical A-RNA duplexes, O5’ oxygens are often located below planes of the nucleobases. The unique structural context of UNCG TL characterized by the presence of neighboring G_L4_ nucleotide in *syn* conformation is causing a local structural change (deformation with respect to canonical **1a** suite of A-RNA duplexes) of the sugar-phosphate backbone around the G_S+1_ nucleotide (**Figure 4**). According to the sugar-phosphate backbone nomenclature,^81^ the typical values of dihedrals around the G_S+1_ phosphate in its native UNCG conformation are on the edge of canonical **1a** suite definition (A-RNA like) with the *γ* dihedral shifted towards higher values of ∼100°, similarly to **1e** suite. The UNCG-specific conformation of the G_L4_-G_S+1_ sugar-phosphate moiety is under potential strain of two steric clashes of G_S+1_(O5’) oxygen with (i) G_S+1_(C8-H8), i.e., the above mentioned 0BPh interaction, and (ii) G_S+1_(O4’) of ribose. The suspected steric clash of 0BPh interaction can be easily relaxed in structural context of A-RNA duplex by slight elongation of C8H8/C6H6…O5’ distances (**Figure S4** in the Supporting Information). However, in the specific structural context of UNCG TL caused by G_L4_ *syn* orientation, the suspected steric clash of 0BPh interaction of G_S+1_ phosphate can be relaxed only at the expense of intensifying the other steric clash between G_S+1_(O5’) and G_S+1_(O4’) oxygens. As a consequence, the G_S+1_ phosphate is prone to flip towards the alternative (noncanonical) *β*_*g*+_*/γ*_*trans*_ state, where the 0BPh G_S+1_(C8H8)…G_S+1_(O5’) interaction is lost.

**Figure 4.**
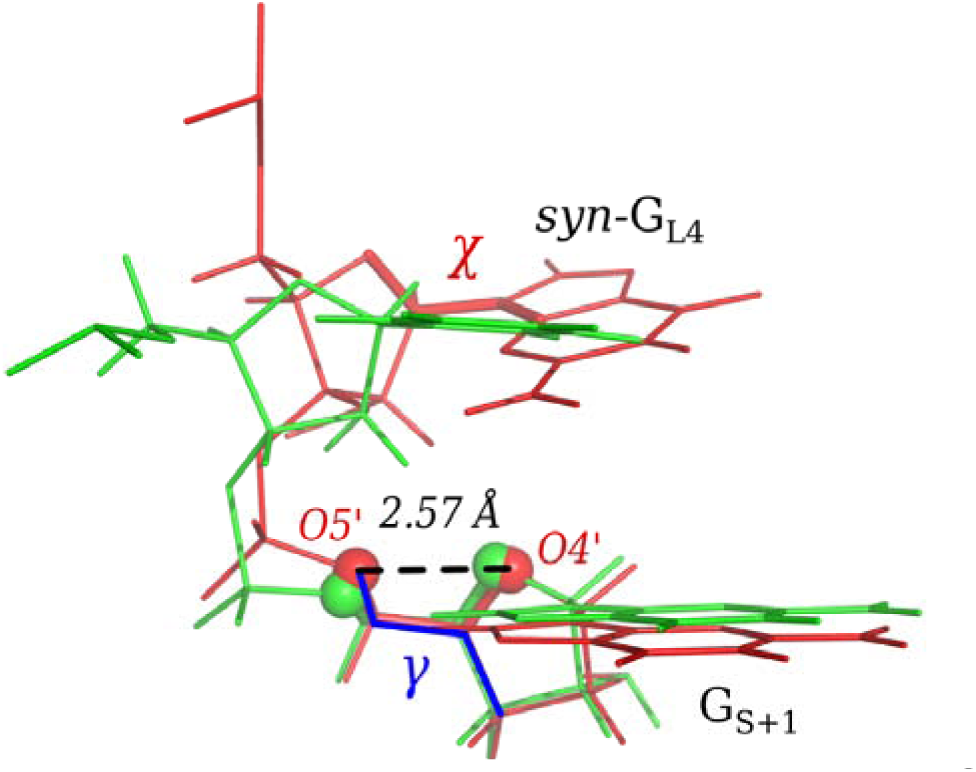
Local deformation of the sugar-phosphate backbone around the G_S+1_ phosphate caused by the *syn* conformation of G_L4_ nucleotide (red) and comparison with the canonical **1a** suite of A-RNA duplex (green). Both superimposed fragments are taken from the starting structure of the r(ggcacUUCGgugcc) 14-mer (PDB ID 2KOC^30^). The UNCG-specific conformation of sugar-phosphate backbone (in red) contains two close contacts of the G_S+1_(O5’) oxygen with (i) G_S+1_(O4’) and (ii) G_S+1_(C8H8) groups, which might cause steric clashes under the *ff* description and then lead to the G_S+1_ phosphate flip. The *χ* dihedral of the *syn* G_L4_ nucleotide, *γ* dihedral of the G_S+1_ nucleotide and the average experimental distance^30^ between O5’ and O4’ atoms are highlighted.

Unambiguous characterization of the G_L4_-G_S+1_ sugar-phosphate moiety is difficult even for experiments as there are only few nuclear Overhauser effect intensities (NOEs) and no ^3^J-scalar couplings available for detailed classification of this structural part.^30^ We realized that some NOEs, namely G_S+1_(H8) – G_S+1_(H4’) and G_S+1_(H8) – G_L4_(H3’) are violated even for the NMR starting structure. Other violations (considering H5’ and H5’’ protons) can be explained by possible neglect of spin diffusion in the NMR experiment. In the MD ensemble those parts of trajectories with native G_S+1_ phosphate conformation have better agreement with the experiment (for ten NOE signals around the G_L4_-G_S+1_ sugar-phosphate moiety) than those with flipped phosphate states except of two signals: (i) G_L4_(H8)…C_L3_(H5’’) and (ii) G_S+1_(H8)…G_S+1_(H5’’). Nevertheless, although the preference of RNA *ff*s for the alternative G_S+1_ phosphate conformation is likely a *ff* artifact, the experimental uncertainties around the G_L4_-G_S+1_ moiety may indicate some real dynamics in this key part of the UNCG TL.

### The 0BPh interaction is incorrectly described by the *ff*

We analyzed the 0BPh interaction using electronic structure calculations. We compared QM/MM and MM behavior of two distances that characterize the 0BPh interaction, namely the G_S+1_(C8H8)…G_S+1_(O5’) and G_S+1_(O4’) – G_S+1_(O5’) interactions. We took the UNCG 8-mer TLs, where the G_S+1_ phosphate maintained its native geometry (starting structures **6a-e, 7a-e** and **9a-e**, see Methods). Fifteen gradient geometry optimizations carried out from these starting structures showed that the RNA *ff* prefers longer distances in comparison with QM by ∼0.13 Å and ∼0.05 Å for the G_S+1_(C8H8)…G_S+1_(O5’) and G_S+1_(O4’) – G_S+1_(O5’) interactions, respectively (**Table 1, Table S1** and **Table S2** in the Supporting Information). Additionally, we used dinucleotide monophosphate model (**Figure S5** in the Supporting Information) possessing the *native* G_S+1_ phosphate conformation and analyzed its QM and MM-optimized geometries. Again, both the G_S+1_(C8H8)…G_S+1_(O5’) and G_S+1_(O4’) – G_S+1_(O5’) distances are longer (by ∼0.26 Å and ∼0.10 Å) in MM compared to the QM description (**Table 1** and **Table S3** in the Supporting Information).

**Table 1.** Comparison of the QM and MM description of the G_S+1_(C8H8)…G_S+1_(O5’) and G_S+1_(O4’) – G_S+1_(O5’) distances (in Å) within UNCG 8-mers and dinucleotide monophosphate models.^a^

| | $G_{S+1}(\text{C8H8})\dots G_{S+1}(\text{O5}')$ | | | $G_{S+1}(\text{O4}') - G_{S+1}(\text{O5}')$ | | |
| --- | --- | --- | --- | --- | --- | --- |
| | initial value | $\Delta d_{\text{QM/MM}}^b$ | $\Delta d_{\text{MM}}^c$ | initial value | $\Delta d_{\text{QM/MM}}^b$ | $\Delta d_{\text{MM}}^c$ |
| UNCG TL |  |  |  |  |  |  |
| Structure **6e** | 2.18 | -0.09 | 0.10 | 2.74 | 0.04 | 0.09 |
| Structure **7e** | 2.39 | 0.00 | 0.09 | 2.70 | 0.05 | 0.13 |
| Structure **9e** | 2.15 | 0.06 | 0.15 | 2.57 | 0.05 | 0.11 |
| Dinucleotide monophosphate |  |  |  |  |  |  |
| | initial value | $\Delta d_{\text{QM/COSMO}}^d$ | $\Delta d_{\text{MM/GB}}^e$ | initial value | $\Delta d_{\text{QM/COSMO}}^d$ | $\Delta d_{\text{MM/GB}}^e$ |
| *planar-native* | 2.37 | -0.14 | 0.13 | 2.76 | -0.05 | 0.11 |
| *tilted-native* | 2.18 | 0.04 | 0.26 | 2.74 | 0.00 | 0.06 |

<sup>a</sup> See Methods for description of structures. UNCG TLs (structures **6**, **7** and **9**) and dinucleotide monophosphate models (*planar-native*, *tilted-native*) possessing the native conformation of $G_{S+1}$ phosphate were chosen for the analysis. UNCG TLs were optimized by QM/MM or MM while the dinucleotide monophosphate models were optimized with geometry restraints by QM/COSMO or MM/GB. For UNCG 8-mer, values are shown only for structures **6e**, **7e**, and **9e**. See Section S3 in the Supporting Information for the explanation and for complete data, i.e., for 8-mer structures with different water distributions (**a-e**) and for dinucleotide monophosphate models optimized without geometry restraints (Tables S1, S2 and S3).
<sup>b</sup> $\Delta d_{QM/MM}$ = QM/MM value – initial value; i.e., change of the distance upon optimization
<sup>c</sup> $\Delta d_{MM}$ = MM value – initial value
<sup>d</sup> $\Delta d_{QM/COSMO}$ = QM/COSMO value – initial value
<sup>e</sup> $\Delta d_{MM/GB}$ = MM/GB value – initial value

We then performed MM interaction- and conformational-energy scans of the G_S+1_(C8H8)…G_S+1_(O5’) distance using two models: (i) a smaller dimethyl-phosphate – methyl-guanine molecular complex lacking the G_S+1_(O4’) – G_S+1_(O5’) contact, and (ii) a larger O3’-methylated-guanosine-monophosphate intramolecular model, which included the G_S+1_ ribose with its O4’ oxygen atom and thus takes into account the possible repulsion between G_S+1_(O4’) and G_S+1_(O5’) atoms (**Figures S6A** and **S6B** in the Supporting Information). The smaller model showed prolonged MM G_S+1_(C8H8)…G_S+1_(O5’) distance (by 0.11 Å) in comparison with QM (**Figure 5**). The difference doubles to 0.22 Å for the larger model (**Figure S7** in the Supporting Information). We suggest that the increased difference (by 0.11 Å) obtained for the larger model is due to the G_S+1_(O4’) – G_S+1_(O5’) repulsion. In addition, the small-model calculations reveal that the PBEh-3c QM method used for the large-scale QM/MM calculations overestimates the G_S+1_(C8H8)…G_S+1_(O5’) H-bond distance by 0.08 Å with respect to the QM reference (**Figure 5**). In other words, the identified overestimation of short-range by the MM is even larger than indicated by the above comparison between PBEh-3c QM/MM and MM data.

**Figure 5.**
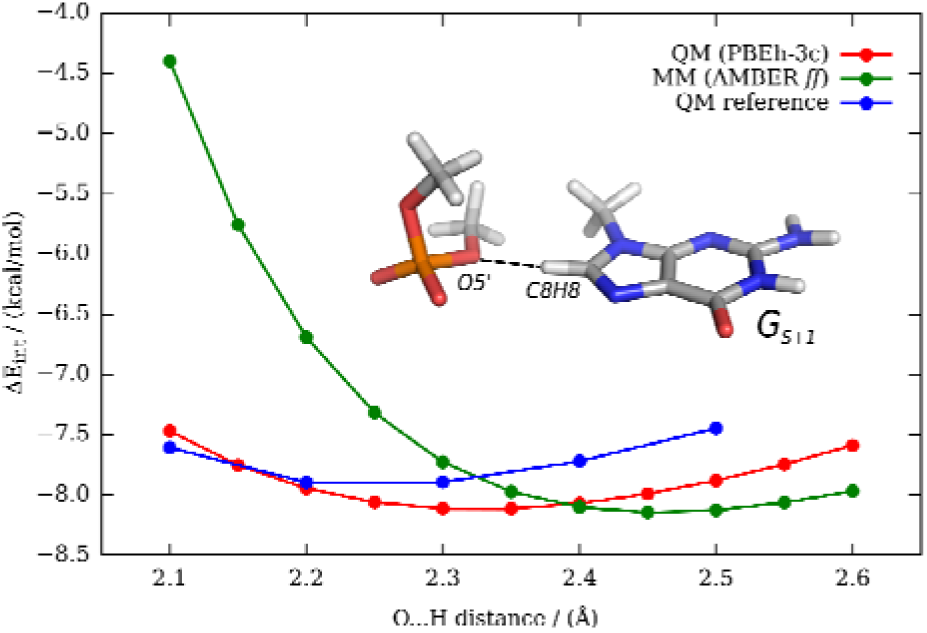
Comparison of the QM (PBEh-3c; red), MM (AMBER *ff* – see Methods; green) and QM reference (DLPNO-CCSD(T)/CBS; blue) interaction energy profiles along the G_S+1_(C8H8)…G_S+1_(O5’) H…O distance of the dimethyl-phosphate – methyl-guanine model. Interaction energy scan was initiated from the G_S+1_ conformation found in the UNCG TL. Optimal distances of 2.32 Å, 2.46 Å and 2.24 Å for QM, MM and QM reference, respectively, show overestimation of the optimal G_S+1_(C8H8)…G_S+1_(O5’) distance by MM. Note that the MM description not only overestimates the optimum H-bond distance, but also sharply exaggerates the steepness of the short-range repulsion, which is a hallmark artifact of the 6-12 Lennard-Jones potentials when describing close interatomic contacts and steric clashes.^82^ Since the computations are done in the gas phase and the system is non-neutral, the magnitude of the overall attraction is overestimated for all three methods compared to the same interaction in its complete environment. However, the relative positions of the curves are expected to be unaffected by this simplification.

In summary, the 0BPh interaction is not properly described by the AMBER RNA *ff*s. Both G_S+1_(C8H8)…G_S+1_(O5’) and G_S+1_(O4’) – G_S+1_(O5’) contacts contribute to the imbalance. The optimal interatomic distance and mainly the short-range repulsion (**Figure 5**) are severely overestimated. We propose that there is enough structural flexibility to overcome this inaccuracy for the canonical A-RNA duplex conformation characterized by the **1a** suite. However, the discrepancy becomes unmasked in the restricted conformational space within the UNCG TL formed by the G_L4_ *syn* nucleotide / G_S+1_ phosphate motif. The overestimated repulsion in the MM approximation forces the G_S+1_ phosphate into the flipped states seen during the MD simulations.

### Relationship between the G_S+1_ phosphate conformation and positioning of the G_L4_ nucleotide

In MD simulations, the angle between planes of the G_L4_ *syn* base and of the closing G_S+1_C_S-1_ base pair fluctuates in the range of ∼10° to ∼30°. We call the arrangements at the edges of the range as *planar* and *tilted* positions of G_L4_. Both positions appear consistent with the NMR data. We have carried out QM analyses of the positioning of the G_L4_ *syn* base, which indicate that the *native* state of the G_S+1_ phosphate could excessively force the G_L4_ nucleotide to the *planar* conformation during MD simulations. This conformation is prone to weakening of the signature U_L2_(2’-OH)…G_L4_(N7) sugar-base H-bond, which may contribute to problems in the MD description. However, we were not able to further quantify this potential MM imbalance due to the uncertainties inherent to the used computations. The full description is available in the Supporting Information.

### The signature U_L2_(2’-OH)…G_L4_(N7) H-bond is imperfectly described by *ff*s

The U_L2_(2’-OH)…G_L4_(N7) sugar-base H-bond is another player in the overall UNCG balance. According to the structural data, it could alternate with the U_L2_(2’-OH)…G_L4_(O6) H-bond.^30^ Melting experiments with various substitutions at the U_L2_(2’-OH) position (preventing the H-bonding) suggested its only negligible effect on the thermodynamic stability of the loop.^83^ Analysis of 10 μs-long standard MD simulations^19^ revealed that the U_L2_(2’-OH)…G_L4_(N7) H-bond is not very stable, since the U_L2_(2’-OH) is pointing either towards the solvent or is too far from both G_L4_(N7) and G_L4_(O6) acceptors in ∼25% of all snapshots with the native TL arrangement. When the U_L2_(2’-OH) is interacting with G_L4_ nucleobase, the native G_L4_(N7) acceptor is preferred over the G_L4_(O6) (∼75% and ∼25%, respectively). In addition, even when formed, the U_L2_(2’-OH)…G_L4_(N7) H-bond shows poor directionality, with the O-H group pointing above the G aromatic moiety rather than directly to the N7 atom. This may be sign of a local imbalance of the *ff* description and understabilization of the U_L2_(2’-OH)…G_L4_(N7) interaction. Importantly, loss of the signature U_L2_(2’-OH)…G_L4_(N7) H-bond allows reconformation of the G_L4_ nucleobase within the loop and formation of the bifurcated G_L4_(N1H/N2H)…U_L1_(O2) H-bond. It is worth mentioning that the bifurcated G_L4_(N1H/N2H)…U_L1_(O2) H-bond is in agreement with some X-ray structures,^21^ however, in simulations its formation is typically followed by flip of the G_L4_ nucleobase out of the loop and subsequent disruption of the native state (**Figure 2B**). The U_L2_(2’-OH)…G_L4_(N7) dynamics is thus directly kinetically related to loss of the native state.

We employed a set of QM calculations to investigate this signature sugar-base H-bond. Initially, we compared donor-acceptor distances and directionality of the U_L2_(2’-OH)…G_L4_(N7) H-bond in QM/MM- and MM-optimized UNCG 8-mers (**Tables S5** and **S6** in the Supporting Information). We observed that although the QM and MM donor-acceptor distances are similar (**Table S5**) the O-H…N angle is smaller (by ∼18° in average) in MM compared to QM (**Table 2**, and **Table S6** and **Figure S9** in the Supporting Information). Thus, we designed a small model (composed of G_L4_ base and U_L2_ ribose; **Figures S6C** and **S6D** in the Supporting Information) and performed interaction energy scans of the U_L2_(2’-OH)…G_L4_(N7) H-bond. We used two different values of the O-H…N angle, i.e. (i) ∼170° (almost ideal angle for the H-bond), and (ii) ∼143° (mimicking the H-bond often seen during MD simulations). MM minimum of the interaction energy profiles is slightly shifted to shorter donor-acceptor distances (by ∼0.15 Å) and is slightly narrower in comparison with the broad QM minimum (**Figure 6**). Thus, any enforced prolongation of the O…N distance by e.g. structural dynamics of other moieties within the loop would result in excessive weakening of the U_L2_(2’-OH)…G_L4_(N7) H-bond under MM description. In addition, the *ff* description obviously neglects the stabilizing effect of X-H bond stretching; this point is not analyzed by our calculations since the QM scan was done with rigid monomers, but it has been discussed elsewhere.^26,53^

**Table 2.** Description of directionality of the signature U_L2_(2’-OH)…G_L4_(N7) H-bond (O-H…N angle in °) in QM/MM and MM optimizations of the whole TL 8-mers.^a^

| structure | initial angle | $\Delta\text{angle}_{\text{QM/MM}}^b$ | $\Delta\text{angle}_{\text{MM}}^c$ |
| --- | --- | --- | --- |
| **1** | 143.4 | 16.1 | 5.2 |
| **2** | 118.1 | 45.2 | 20.8 |
| **4** | 140.8 | 14.4 | -12.6 |
| **5e** | 161.7 | 9.3 | 2.5 |
| **6e** | 157.0 | 5.5 | -7.0 |
| **9e** | 143.5 | 25.6 | -0.9 |

<sup>a</sup> structure **3** was excluded because the $U_{L2}(2'-OH)$ forms an alternative H-bond with $G_{L4}(O6)$ instead of $G_{L4}(N7)$ atom. Structures **7** and **8** were excluded because they contain spurious state of the $U_{L1}(2'-OH)$ group, which may influence position of the $G_{L4}$ nucleobase and subsequently the $U_{L2}(2'-OH)\dots G_{L4}(N7)$ H-bond. Values are
<sup>b</sup> $\Delta\text{angle}_{\text{QM/MM}} = \text{angle}_{\text{QM/MM}} - \text{angle}_{\text{initial}}$ ; i.e., change of the angle upon optimization
<sup>c</sup> $\Delta\text{angle}_{\text{MM}} = \text{angle}_{\text{MM}} - \text{angle}_{\text{initial}}$

**Figure 6.**
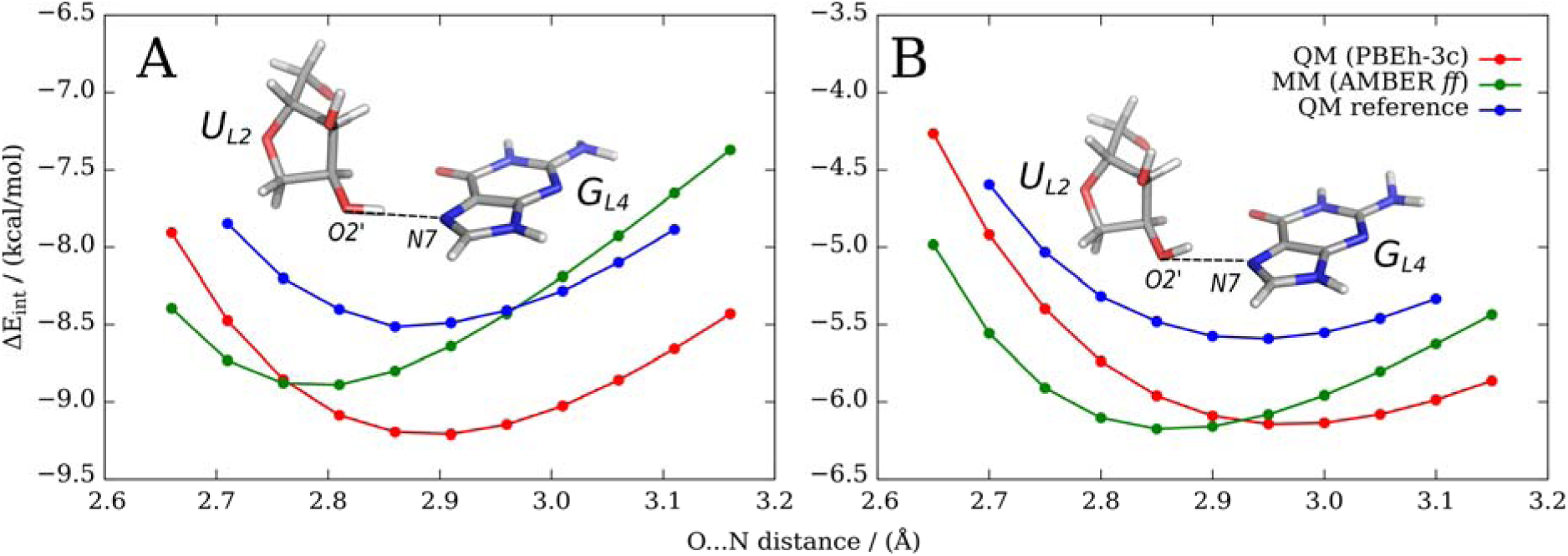
Interaction energy scans of the U_L2_(2’-OH)…G_L4_(N7) H-bond using QM (PBEh-3c; red), MM (AMBER *ff*; green) and QM reference (DLPNO-CCSD(T)/CBS; blue) methods. Ribose is shifted along the O…N vector in a distance range from 2.65 Å to 3.15 Å with the step length of 0.05 Å. All remaining degrees of freedom were frozen (see the Supporting Information for details). **(A)** The O-H…N angle is 170° at start of the scan. **(B)** The O-H…N angle is initially 143°.

More importantly, comparison of QM and MM PES’s revealed that the effect of loss of directionality of this H-bond, i.e. comparing the almost ideal H-bond angle (∼170°) with the lowered one (∼143°) is underestimated (by ∼0.7 kcal/mol) by the MM method (**Figure 6**). Although MM still prefers (in the isolated small system) a H-bond with good directionality, the penalty for making the H-bond less directional is lower at the MM level compared to the QM level. Therefore, if some other energy terms in the UNCG TL profit from non-planarity of this H-bond due to the overall conflict among some other energy terms, the MM description may be excessively submissive to such strains.

Next, we used the U_L2_ ribose model and analyzed behavior of the C2’-O2’-O2’H angle and C1’-C2’-O2’-O2’H dihedral by QM and MM optimizations (**Figure S10** in the Supporting Information). We performed two conformation energy scans using the model: (i) C2’-O2’-O2’H angle bending and (ii) rotation of the C1’-C2’-O2’-O2’H dihedral. Both QM and MM conformational scans were comparable for the bending of the C2’-O2’-O2’H angle (**Figure S11** in the Supporting Information) and free optimizations showed a slightly shifted minimum (by 2.8°) between the QM and MM levels of theory (**Table S7** in the Supporting Information). In contrast, conformational energy scans of the C1’-C2’-O2’-O2’H dihedral revealed differences especially around the region of interest, where the U_L2_(2’-OH)…G_L4_(N7) H-bond could be established (conformations with dihedral values between ∼150° and ∼210°, **Figure 7**). We identified a very flat QM minimum (∼190°) that is completely absent on the MM potential energy surface (PES). Instead, the MM curve is steeply descending in this region until it finds a deeper minimum around 260° (**Figure 7** and **Table S7** in the Supporting Information). This result is in agreement with analysis published by Mladek and coworkers^84^ using larger sugar-to-sugar models (see **Section S4** in the Supporting Information for more details).

**Figure 7.**
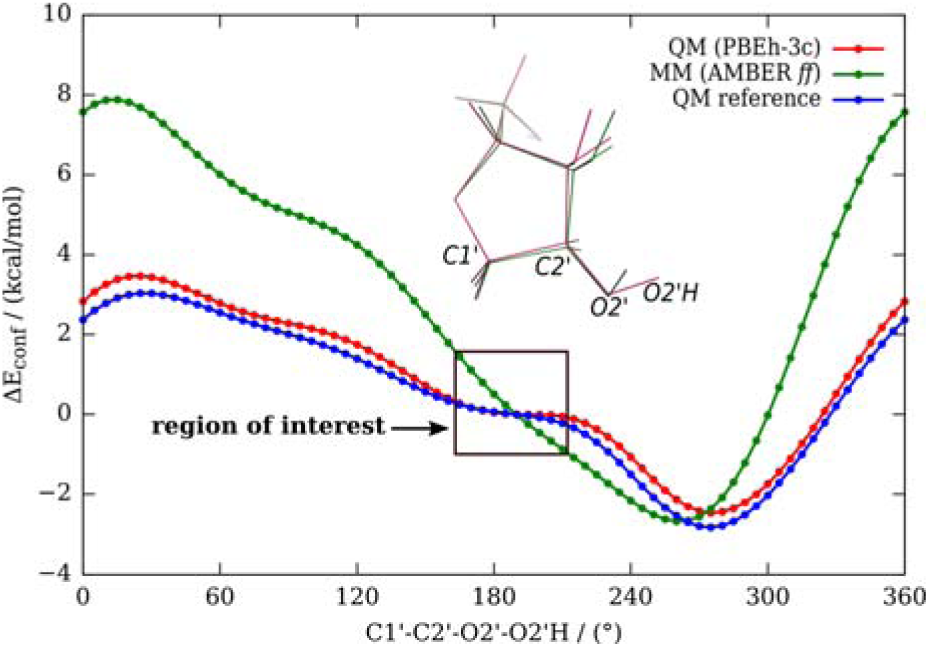
Conformation energy profile of the C2’-*endo* U_L2_ ribose C1’-C2’-O2’-O2’H dihedral using QM (PBEh-3c; red), MM (AMBER *ff*; green) and QM reference (DLPNO- CCSD(T)/CBS; blue) methods. Zero energy is set for a dihedral value of 190° (**Figure S10** in the Supporting Information). Region of interest where the U_L2_(2’-OH)…G_L4_(N7) H-bond could be established is highlighted by the black rectangle. While QM finds flat region ∼190° MM continues steeply to the global minimum at ∼260°, where the intramolecular O2’H…O3’ interaction is formed (**Table S7** in the Supporting Information). See Section S3 in the Supporting Information for computational details and Section S4 and ref. ^84^ for comparison with the earlier scans of the 2’-OH group.

We note that after the U_L2_(2’-OH)…G_L4_(N7) or the alternative U_L2_(2’-OH)…G_L4_(O6) H-bond is lost in MD simulations, the U_L2_(2’-OH) group binds to another acceptor within the loop. Thus, it is meaningless to analyze a shift of the C1’-C2’-O2’-O2’H dihedral once the H-bond to G_L4_ is lost. However, QM/MM and MM optimizations of UNCG TL 8-mers revealed that the C1’-C2’-O2’-O2’H dihedral is larger in MM-optimized geometries (**Table S8** in the Supporting Information), which is possibly associated with non-planarity of the U_L2_(2’-OH)…G_L4_(N7) H-bond in MD simulations. Thus, all calculations indicate that both directionality and strength of the signature U_L2_(2’-OH)…G_L4_(N7) H-bond are not optimally described by the *ff*, which may contribute to the common loss of this H-bond during MD simulations of UNCG TL.

Any large imbalance in description of the 2’-OH…N interactions could have major impact on RNA simulations, as these sugar-base H-bonds are abundant.^85^ Further, correction of such imbalance would be difficult, due to lack of specific *ff* terms to affect directionality of H-bonds. Thus, we performed a comparable analysis for the two 2’-OH…N1(A) H-bonds in an RNA kink-turn. However, in contrast to the UNCG TL case, these H-bonds behave stably (see **Section S5, Table S9** and **Figure S12** in the Supporting Information for details). We thus suggest that the accuracy of the description of the 2’-OH…N interactions is generally acceptable for simulations of majority of RNAs and the moderate imbalance towards non-planarity is unmasked for the UNCG TL due to its uniquely tight conformation.

### The U_L1_(2’-OH)…G_L4_(O6) signature sugar-base H-bond is weakened by flips of the U_L1_(2’-OH) towards the U_L2_ phosphate

The signature U_L1_(2’-OH)…G_L4_(O6) sugar-base H-bond is crucial for stability of the UNCG TL. It should be firmly established according to the structural data^30^ and any substitution at the U_L1_(2’-OH) position leads to a significant drop of the thermodynamic stability.^83^ MD simulations show reversible loss of the U_L1_(2’-OH)…G_L4_(O6) H-bond and transient formation of an alternative U_L1_(2’-OH)…U_L2_(O5’) H-bond, a known *ff* artifact that was firstly classified in ref. 21. We performed several standard MD simulations to understand this flip and its dependency on other factors. Although the signature U_L1_(2’-OH)…G_L4_(O6) H-bond is favored, there is a 10-30% population of the competing U_L1_(2’-OH)…U_L2_(O5’) H-bond. The latter H-bond is typically short-living (∼hundreds of ps) and the flips are entirely reversible. The flips were more frequent in SPC/E solvent in comparison with TIP3P and OPC water models (**Table S10** in the Supporting Information). Moreover, the flips were more common in 1.0 M KCl salt-excess ionic conditions than with 0.15 M KCl ion-strength and K^+^ net-neutral ionic conditions. The modified vdW parameters for phosphate oxygens^40^ did not bring any improvement (**Table S10** in the Supporting Information).

We then investigated competition between the signature U_L1_(2’-OH)…G_L4_(O6) and alternative U_L1_(2’-OH)…U_L2_(O5’) H-bond using QM methods. Firstly, we analyzed the QM/MM- and MM-optimized 8-mer TLs with native U_L1_(2’-OH)…G_L4_(O6) H-bond and observed comparable QM and MM description of this H-bond (structures **1**-**5, 6a-e** and **9a-e**). Then we analyzed structures **7a-e** and **8a-e** where the U_L1_(2’-OH) group is flipped towards the U_L2_ phosphate. For the structure **7** having the *native* state of the G_S+1_ phosphate, we observed renewal of the signature U_L1_(2’-OH)…G_L4_(O6) H-bond in all five (**a**-**e**) QM/MM optimizations, whereas in equivalent MM optimizations the U_L1_(2’-OH) group remained flipped towards the U_L2_(O5’) (see the Supporting Information for starting and final structures). Note that the starting structures **a-e** used for optimizations differ only in solvent equilibration while the RNA configuration is identical (see Methods).

Subsequently, we performed relaxed MM/MM and QM/MM scans (see Methods for details) of the UNCG TL 8-mers along the C1’-C2’-O2’-O2’H dihedral to quantify the potential energy curves between the two conformations (**Figure 8**). We used MD snapshots with different states of the G_S+1_ phosphate and different orientations of the U_L1_(2’-OH) group as starting points, namely: (i) *native* state of the G_S+1_ phosphate and *native* orientation of the U_L1_(2’-OH) group (structure **9**), (ii) *native* G_S+1_ phosphate and *flipped* U_L1_(2’-OH) group (structure **7**), (iii) *flipped* state of the G_S+1_ phosphate and *native* orientation of the U_L1_(2’-OH) group (structure **2**), and (iv) *flipped* G_S+1_ phosphate and *flipped* U_L1_(2’-OH) group (structure **8**).

**Figure 8.**
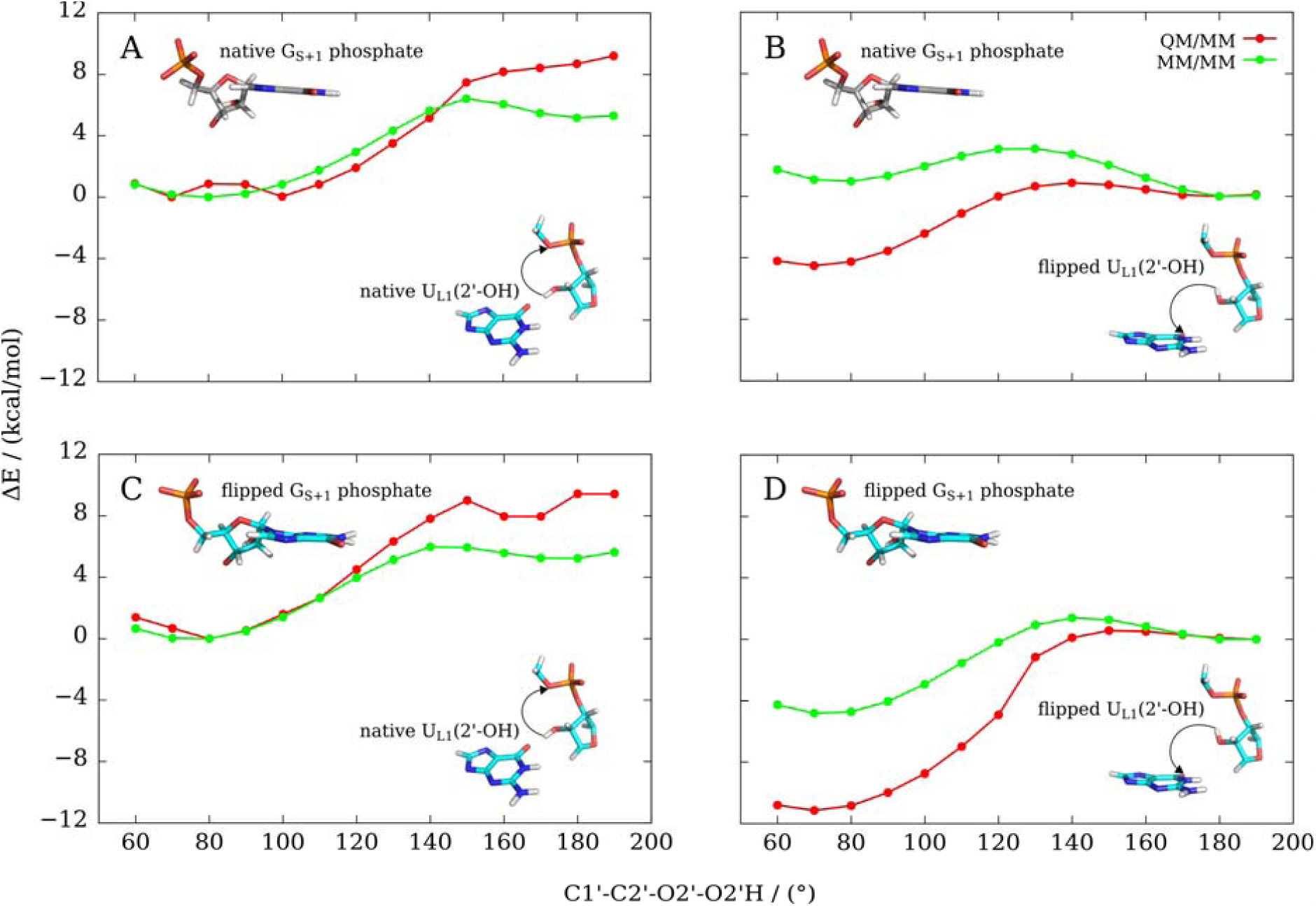
Comparison of the conformational energy profiles of the U_L1_ ribose along the C1’- C2’-O2’-HO2’ dihedral for QM/MM (red) and MM/MM (green) methods. Initial structural orientations of the G_S+1_ phosphate and the U_L1_(2’-OH) group are highlighted as structural insets and directions of scans are shown as black arrows.

For all four scans (**Figure 8**), the MM/MM calculation (the green curves) overstabilizes the structure with alternative U_L1_(2’-OH)…U_L2_(O5’) H-bond (the C1’-C2’-O2’-O2’H dihedral ∼180°) with respect to the native U_L1_(2’-OH)…G_L4_(O6) H-bond (the C1’-C2’-O2’-O2’H dihedral ∼80°), relatively to the QM/MM data (red curves).

For UNCG TLs with native G_S+1_ phosphate conformation we observed that going from the native H-bond (∼80°) towards the alternative H-bond (∼180°), the QM approach did not find any second minimum for ∼180° (**Figure 8A**). In contrast, the alternative orientation of the U_L1_(2’-OH) group towards U_L1_(O5’) was recognized as a local minimum by MM (**Figure 8A**). Starting from the MD snapshot with the alternative H-bond, both QM/MM and MM/MM approaches identified the alternative state as a minimum. However, while for the QM/MM scan the native orientation of the U_L1_(2’-OH) group was the global minimum, the alternative orientation was the global minimum of the MM/MM scan (**Figure 8B**).

Energy scans with the G_S+1_ phosphate in the flipped conformation (shift of *β/γ* dihedrals and absence of the G_S+1_(C8H8)…G_S+1_(O5’) 0BPh interaction) revealed the following picture. Starting from the native orientation of the U_L1_(2’-OH) group both QM/MM and MM/MM approaches indicated that the global minimum is located near ∼80°and the alternative orientation has no local minimum (**Figure 8C**). However, the MM/MM curve clearly underestimates the energy difference. Starting from the alternative state of the U_L1_(2’-OH) group, both QM/MM and MM/MM indicated the native orientation as the global minimum of the scan. However, the MM/MM curve is again substantially shifted in favor of the U_L1_(2’-OH)…U_L2_(O5’) H-bond relatively to the QM/MM curve (**Figure 8D**).

The MM/MM energy scans were performed with the modified vdW parameters for phosphate oxygens (CP). We thus performed the MM/MM energy scan for structure **8** also without the CP modification. The difference was negligible (**Figure S13** in the Supporting Information).

In summary, all PES scans showed that formation of the alternative U_L1_(2’-OH)…U_L2_(O5’) H-bond is overstabilized by the MM potential. In addition, in the simulations, the spurious and reversible flipping of the U_L1_(2’-OH) group (∼180°) is connected with dynamics of the G_L4_-*syn* nucleotide by preferring the planar G_L4_ conformation. It has, subsequently, a higher tendency to depart from the loop and collapsing it (see the paragraph above about the disruption of the UNCG TL).

We note that the energy scans should always be considered within the limits of sampling inherent to mere structure optimizations.^26^ Detailed inspection of **Figure 8** shows that there is an apparent inconsistency of (especially) the MM curves between panels A – B and C – D, i.e., between scans with the same G_S+1_ phosphate conformation. This indicates that other (still unidentified) structural factors are probably involved. We explicitly inspected those mentioned in our earlier study,^21^ i.e., *ε*(U_L1_), *ε*(C_S-1_), *ζ*(C_S-1_), and *ζ*(U_L1_) dihedrals as well as sugar pucker of the U_L1_ nucleotide but we did not find any clear structural changes along the PES curves. Thus, it appears that the flipped state of the U_L1_(2’-OH) group might cause some structural adjustments distributed across many parameters that are difficult to identify (see the Supporting Information for initial and final structures from PES scans for comparison).

### The C_L3_(O4’)…G_L4_ sugar-base stacking is slightly over-repulsive in AMBER *ff*

Sugar-base stacking between C_L3_(O4’) and G_L4_ nucleobase belongs to the key characteristics of the ‘Z-like’ motif.^86^ This contact represents a moderately attractive vdW interaction^87–89^ which, however, appears to be rather imposed by the overall structural context than being formed on its own.^89^ As it includes an exceptionally short distance between the O4’ atom and the nucleobase ring, its MM description may be challenging, which may lead to destabilization of the position of the C_L3_ ribose above the G_L4_ nucleobase in MD simulations. Indeed, loss of the sugar-base stacking during MD simulations is associated with flip of the G_L4_ nucleobase out of the loop. Thus, we analyzed distance of the C_L3_(O4’) atom to the G_L4_ *syn* base plane in our QM/MM- and MM-optimized UNCG 8-mers. MM overestimates the O4’ – guanine distance by ∼0.04 Å compared to QM/MM (**Table S11** in the Supporting Information). Additional calculations using a small ribose-guanine sugar-base stacking model (**Figure S6E** in the Supporting Information) showed that MM prefers a larger optimal distance (by ∼0.13 Å; **Figure S15**) compared to QM (PBEh-3c). In summary, short-range repulsion of the sugar-base stacking is overestimated by MM, which likely also contributes to disruption of the UNCG loop during MD simulations, especially when assuming that the sugar-base stacking is a consequence of the overall complex topology of UNCG TL.

### Large-scale implicit solvent QM calculations are consistent with the QM/MM data

In addition to QM/MM and MM calculations performed in the explicit solvent, we also tested the effect of implicit solvation for QM and MM geometry optimizations of the UNCG TL. We used the COSMO implicit solvent for the QM calculations which are compared with equivalent MM/GB calculations. The results obtained with implicit solvation are consistent with the above QM/MM and MM data in explicit solvent and are discussed in Section S3 of the Supporting Information.

### Attempts to improve the UNCG TL simulations

We made several attempts to improve the UNCG simulations by simple corrections based on the above investigations. We tested REST2 folding simulations of the r(gcUUCGgc) 8-mer and standard simulations of the r(ggcacUUCGgugcc) 14-mer. The former test includes the complete folding landscape, but the simulations could be affected also by stability of the two-base-pair stem, which may be underestimated by the *ff*. For the 14-mer, we monitored transitions between the fully native conformation and G_L4_ bulged-out conformation.^6,23,79^

### NBfix correction of the 0BPh interaction maintains the G_S+1_ native phosphate conformation

We first attempted to eliminate the G_S+1_ phosphate flipping. We modified the pairwise vdW parameters via breakage of the combination (mixing) rules by using the so-called nonbonded fix (NBfix).^90^ We changed the minimum-energy distance of Lennard-Jones potential (i.e., *R*_*i,j*_ parameter) for the –H8…O5’– pair (i.e., between H5 and OR atom types) to 2.8808 Å. The depth of the potential well (*e*_*i,j*_ parameter) was kept at its default value of 0.0505 kcal/mol. In other words, we decreased the sum of vdW radii between H8 atoms of all purine bases and O5’ oxygens of phosphates by 0.25 Å.

The effect of the NBfix correction was significant for both the interaction energy scan of the dimethyl-phosphate – methyl-guanine model (see paragraph The 0BPh interaction is incorrectly described by the *ff*) and MD simulations of the r(ggcacUUCGgugcc) 14-mer. For former case, we changed the G_S+1_ –H8…O5’– pair of a small model and it resulted in significant improvement of MM interaction energies with shift of the optimal distance towards QM level description (**Figure S16**). For the later, the NBfix correction was applied for all six purine bases of the r(ggcacUUCGgugcc) 14-mer. The G_S+1_ phosphate favored its native geometry compared to the flipped one (∼80%/∼20%, **Figure S17** in the Supporting Information). The NBfix correction shifts the G_S+1_(C8H8)…G_S+1_(O5’) distance histograms towards shorter values (**Figure S18** in the Supporting Information). Thus, elimination of the clash stabilizes the 0BPh interaction and the native phosphate moiety position, as intended. However, it does not bring any overall help to the TL simulations. The disruption of the loop in standard simulations is even accelerated once the native G_S+1_ phosphate conformation is stabilized; in one of the three simulations G_L4_ left its binding pocket just after ∼35 ns (**Figure S17** in the Supporting Information). Hence, elimination of the G_S+1_ phosphate flipping, which is partially easing the apparent conformational stress within the UNCG loop, is actually speeding up loss of the TL structure initiated by the G_L4_ *syn* nucleotide departure. As discussed above, the local flipped backbone conformation “sequesters” G_L4_ in the pocket away from the global unfolding pathway. This is a typical example of compensation of errors, in this case affecting lifetime of the folded state. Thus, despite the lack of improvement in simulations, we consider the modification of 0BPh interaction as correct *per se*.

### gHBfix alone is insufficient to fold the gcUUCGgc TL

gHBfix potentials are general *ff* terms added to the basic *ff* for tuning stability of H-bonds. In our previous work we have shown that combining the basic *χ*_OL3CP_ *ff* with the gHBfix19 improves RNA simulations without side-effects, though it was still insufficient to fold the UNCG TL.^19^

We have thus tried several additional gHBfix settings to specifically improve folding of the r(gcUUCGgc) 8-mer. The most complex version strengthened base – base H-bonds and weakened sugar – phosphate interactions comparably to the gHBfix19 potential. In addition it also strengthened the sugar donor – base acceptor H-bonds (as they represent key signature H-bonds in the UNCG TL) and weakened base donor – sugar acceptor and sugar – sugar H-bonds. This version is abbreviated as gHBfix_UNCG19_ (see Supporting Information for details). With gHBfix_UNCG19_ we have been capable to see occasional UNCG folding events but the overall population of the folded TL 8-mer remained negligible. Full methods and results are summarized in the Supporting Information.

### Proposed pathway of the G_L4_ departure and insertion

Although the gHBfix_UNCG19_ variant is not sufficient to correctly fold the r(gcUUCGgc) 8-mer, it gives interesting insights. We analyzed all available REST2 simulations of the r(gcUUCGgc) 8-mer as well as the r(ggcacUUCGgugcc) 14-mer standard simulations showing some capability of G_L4_ to return back to the native position after its initial departure, i.e., from the bulge-out state^6,79^ back to the native state.^19,23^ We have found that the folding (and refolding) pathway requires a conformation, where the G_S+1_ phosphate is in the native state. Thus, the identified *ff* imbalance between native and flipped conformation of the G_S+1_ phosphate is kinetically stabilizing the broader native basin (including the flipped G_S+1_ phosphate) once reached, as discussed in the previous section. However, it also hinders folding of the TL as the folding pathway goes through a bottleneck with the native G_S+1_, which is enthalpically disfavored by the *ff* due to steric clash associated with the 0BPh interaction (**Figure 9**). This explains why attempts to tune solely the stability of H-bonds were insufficient to correct the free-energy imbalance between native and misfolded states of the r(gcUUCGgc) TL.

**Figure 9.**
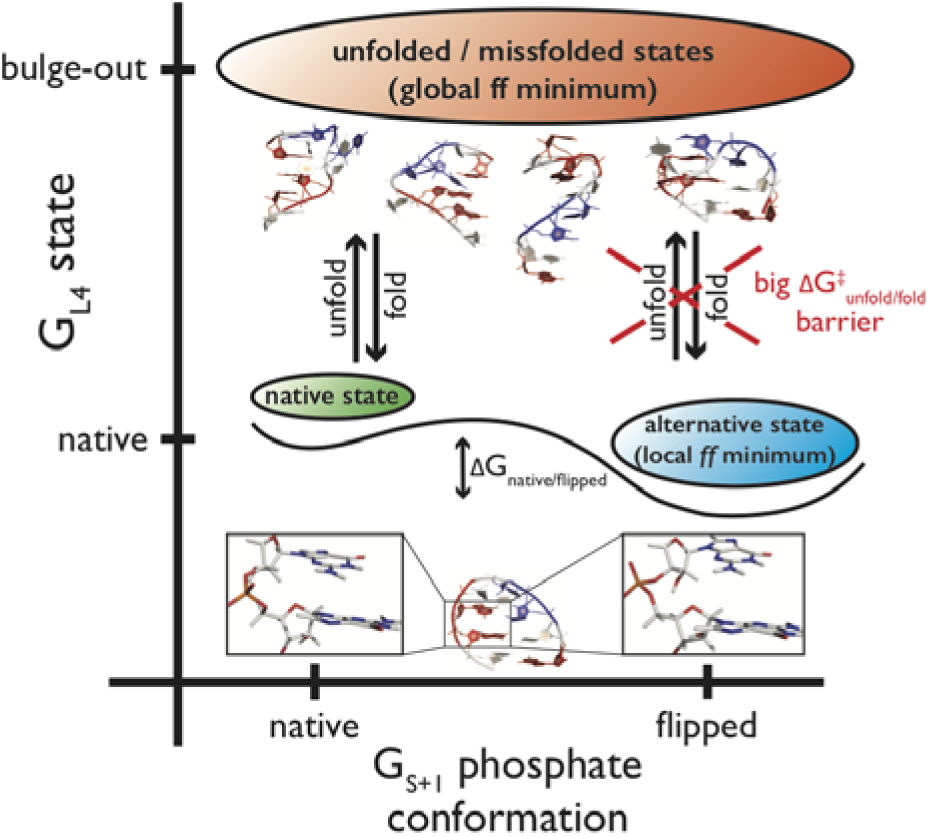
Proposed scheme describing folding and unfolding pathways of the r(gcUUCGgc) TL in MD simulations based on key conformational states of the G_S+1_ phosphate and G_L4_ nucleotide. The main conformational states are highlighted: (i) native state with both G_L4_ and G_S+1_ phosphate in native conformation, (ii) alternative state (local minimum in *ff*) with G_L4_ in native and G_S+1_ phosphate in flipped conformation, and (iii) global *ff* minimum including all misfolded and unfolded states. Both folding and unfolding pathways require the G_S+1_ phosphate in the native conformation. The diagram explains why eliminating just the flipped G_S+1_ phosphate structures does not help to improve the simulations. Representative native structure of the r(gcUUCGgc) TL is shown at bottom and both native and flipped G_S+1_ phosphate conformations are highlighted as insets. Some examples of misfolded structures often found in MD ensembles are also shown (top). G, C, and U nucleotides are in red, white and blue, respectively.

### Final set of REST2 simulations

As an additional effort, we took the following *ff* version: the standard *χ*_OL3CP_ RNA *ff* combined with the gHBfix_UNCG19_ correction for H-bonds and NBfix correction for the –H8…O5’– atom-pair clash. In addition, we reduced the vdW radii of all non-polar H atoms (H1, H4, H5 and HA atoms) universally to 1.2 Å. We have tried this additional vdW modification as it seemed to us that these hydrogens may cause some steric conflicts in the native UUCG conformation, similarly to the 0BPh interaction. The depths of the Lennard-Jones potentials were not modified. All these fixes were aimed to stabilize the 0BPh interaction and further ease some possible – yet unidentified – steric clashes within the UNCG loop. We performed 20 μs-long REST2 folding simulation of the r(gcUUCGgc) 8-mer (all replicas initiated from the completely unfolded single stranded structure). There was an increased probability of folding (**Figure S20** in the Supporting Information) though the overall population of the native state within the reference replica was still low. We obtained ∼33% population of structures with correct stems and ∼7% population of entirely correct native structure. This means that population of the correctly structured loop in the sub-ensemble with correct stem formation was ∼20%. Thus, the introduced corrections increased the propensity of folding but were still not able to fully stabilize the native state within the reference replica. Nevertheless, it is probably the best folding simulation that has been achieved for the 8-mer UNCG TL system by any currently available *ff*.

As the last attempt, we carried out another REST2 simulation starting now from the native state (in all replicas), where we added to the above potential a restraint (see the Supporting Information for details) to the U_L1_(C1’-C2’-O2’-O2’H) dihedral to prevent flipping of the U_L1_(2’-OH) group. It aimed to stabilize the signature U_L1_(2’-OH)…G_L4_(O6) H-bond instead of the alternative U_L1_(2’-OH)…U_L2_(O5’) H-bond. In this simulation, the native state was lost in all replicas after ∼7.5 μs (**Figure S21** in the Supporting Information) with no tendency for refolding, which may indicate that the dihedral restraint caused some undesired side-effects.

## CONCLUDING REMARKS

The compact and well-determined native conformation of the UNCG TL has served as a hallmark reminder of persisting problems of RNA *ff*s for the structural description of RNA molecules.^4,6,8–11,19,21,79,91^ In this work we clarify why this RNA motif is so difficult for MD simulations.

We first inspected in detail the disruption (unfolding) paths of the UUCG motif during standard as well as enhanced sampling MD simulations and identified common rearrangements that precede disruption of the UUCG native topology. Collapse of the loop is characterized by a gradual loss of several key structural features: (i) G_L4_-G_S+1_ phosphate moiety conformation, (ii) G_S+1_(C8H8)…G_S+1_(O5’) 0BPh interaction, (iii) C_L3_-G_L4_ sugar-base stacking interaction, and (iv) both U_L1_(2’-OH)…G_L4_(O6) and U_L2_(2’-OH)…G_L4_(N7) sugar-base H-bonds.

We then performed numerous QM/MM and high-level QM calculations that reveal diverse limitations of the empirical *ff* potential to properly describe the complex network of energy contributions shaping up the UNCG TL. Namely, we identified that both directionality and strength of the signature U_L2_(2’-OH)…G_L4_(N7) H-bond are not well described by the *ff*. The second signature U_L1_(2’-OH)…G_L4_(O6) H-bond is weakened by formation of the alternative U_L1_(2’-OH)…U_L2_(O5’) H-bond, which is overstabilized by the *ff*. Flipping of the U_L1_(2’-OH) group (from base to phosphate) is connected with moderate relocation of the G_L4_-*syn* nucleotide within the binding pocket. Subsequently, G_L4_ has higher tendency to depart from the loop. Further tests indicate that the sugar-base stacking interaction also suffers from a problematic MM description.

We realized that collapse of the UUCG TL is connected with intrinsic dynamics of the G_L4_-G_S+1_ sugar-phosphate moiety. Experimental characterization of this region is ambiguous as there are only few direct experimental observables^30^ and some NOEs are violated even in the NMR-based models. MD simulations show that there are two states of G_S+1_ phosphate, i.e., native and flipped one, characterized by the shift of *β*_*trans*_*/γ*_*g*+_ dihedrals towards alternative *β*_*g*+_*/γ*_*trans*_ conformations. Thus, it appears that there is some intrinsic dynamics within this key part of the UNCG TL, which could be an analogy to recent indication of dynamic behavior of the G_L4_-*syn* nucleotide.^79^ Nevertheless, preference of RNA *ff*s for the alternative G_S+1_ phosphate conformation is clearly a *ff* artifact caused by incorrect description of the G_S+1_(C8H8)…G_S+1_(O5’) 0BPh interaction. Its optimal interatomic distance and short-range repulsion are severely overestimated in *ff*s, forcing the G_S+1_ phosphate into the flipped state seen during the MD simulations.

Although being spurious, population of the flipped G_S+1_ substate kinetically decelerates the unfolding. This is because folding (and refolding) pathways transit through a conformation having the G_S+1_ phosphate in the native state. Hence, also folding pathways reaching the (artificial) local minimum involving the flipped G_S+1_ phosphate have to pass through the native state conformation, which is enthalpically disfavored by the overestimated G_S+1_(C8H8)…G_S+1_(O5’) steric clash in AMBER RNA *ff*s. This is probably the reason why all our previous attempts of tuning solely the H-bond interactions were insufficient to correct the free-energy imbalance between native and misfolded states of the r(gcUUCGgc) TL.^19^

We attempted to fix the 0BPh clash by the NBfix correction, in order to stabilize the native G_S+1_ phosphate conformation. This, however, accelerated disruption of the loop in standard simulations. Nevertheless, coupling this correction with the gHBfix_UNCG19_ potential specifically designed to support H-bonds important for the UNCG TL improved REST2 folding simulations of the r(gcUUCGgc) 8-mer. We achieved ∼7% total population of the fully correct (both loop and stem) native structure while population of the correctly structured loop upon stem formation was ∼20%. It is fair to admit that this result is still not fully satisfactory and was achieved by extended *ff* modifications that might cause side-effects for other RNA systems, which we did not test.

In conclusion, the UNCG TL remains a considerable challenge for atomistic simulations with classical *ff*s. We have identified a number of individual energy imbalances without claiming to achieve a complete coverage of the problems. Our data could serve as an important guidance for tuning of pair-additive as well as polarizable *ffs*.^92,93^ However, when trying to design *ff* corrections based on this knowledge, we still could not get a decisive improvement. There could be several reasons for this. For example, there could be additional imbalances at the stem/loop interface that remain to be identified. Additionally, the extent of how simulations are affected by the uncorrected sugar-base stacking interaction is unknown. Lastly, a more fundamental problem could be the simplicity of the *ff per se*. Even when we correctly identify the imbalances, it could be challenging to suggest their correction with simple MM terms that fail to mimic the true electronic structure effects. As the key result, our work reveals that the poor behavior of UNCG simulations cannot be explained by a singular dominant factor that would be straightforwardly correctable. There is a concerted effect of multiple and mutually coupled *ff* inaccuracies, acting in the context of the uniquely tight conformation of the UNCG TL, where the RNA chain does not provide a sufficient conformational flexibility to absorb or circumvent the individual imbalances. In addition, tuning the simulation performance of the UUCG TL is complicated due to the enormous computer demands of the folding simulations and difficulty to reach their quantitative convergence.

## Supporting information

Supporting Information to the article

## ASSOCIATED CONTENT

### Supporting Information

The following files are available free of charge https://pubs.acs.org. Details about MD protocol, list of small models, additional computational details of small models, additional details about conformation scans of the 2’-OH group, computational results of the kink-turn Kt-7, details about large scale QM calculations in implicit solvent, relationship between G_S+1_ phosphate conformation and positioning of the G_L4_ nucleotide, additional results from folding of the gcUUCGgc TL using the gHBfix potential, description of the U_L1_(C1’-C2’-O2’-O2’H) dihedral restraint, supporting tables and figures (PDF). Starting and final structures from QM/MM, MM optimizations and PES scans (PDF). Video showing the common UNCG disruption pathway during MD simulations (MP4).

## ACKNOWLEDGMENT

This work was supported by Financial support from the Operational Programme Research, Development and Education, European Regional Development Fund project no. CZ.02.1.01/0.0/0.0/16_019/0000754 [to M.O., P.B., P.K.] of the Ministry of Education, Youth and Sports of the Czech Republic, by Praemium Academiae (J.S.), Czech Science Foundation (20-16554S to M.K., V.M. and J.S., 18-25349S to P.B., P.K.), and by the project SYMBIT reg. number: CZ.02.1.01/0.0/0.0/15_003/0000477 financed by the ERDF (H.K., J.S., P.P.).

## ABBREVIATIONS

RNA, UNCG, tetraloop, molecular dynamics, force field, QM/MM

## Notes

### Competing Interest Statement

The authors have declared no competing interest.

