## Supporting Information to the article for "UNCG RNA tetraloop as a formidable force-field challenge for MD simulations"

**S1. Details about MD protocol.** The minimum distance between box walls and solute was 12 Å. The RNA molecule remained constrained during energy minimization of water and ions. Subsequently, all RNA atoms were frozen and the solvent molecules with counter-ions were allowed to move during a 500-ps long MD run under NpT conditions (p = 1 atm., T = 298.16 K) in order to relax the total density. After this, the RNA molecule was relaxed by several minimization runs, with decreasing force constant applied to the sugar-phosphate backbone atoms. Subsequently, the system was heated in two steps: the first step involved heating under NVT conditions for 100 ps, whereas the second step involved density equilibration under NpT conditions for additional 100 ps. The particle mesh Ewald (PME) method for treating electrostatic interactions was used. The standard unbiased MD simulations were performed under periodic boundary conditions in the NpT ensemble at 298.16 K using weak-coupling Berendsen thermostat[1] with coupling time of 1 ps. The SHAKE algorithm, with a tolerance of $10^{-5}$ Å, was used to fix the positions of all hydrogen atoms, and a 10.0 Å cut-off was applied to non-bonding interactions to allow a 2-fs integration step.

**S2. List of small models**

| | |
|---|---|
| **Figure S5** | Dinucleotide monophosphates |
| **Figure S6** | **A** – Dimethyl-phosphate – methyl-guanine |
| | **B** - O3’-methylated-guanosine-monophosphate |
| | **C** – guanine-ribose (170°) |
| | **D** – guanine-ribose (143°) |
| | **E** – sugar-base stacking model |
| **Figure S10** | Optimized $U_{L2}$ C2’-*endo* ribose model |

### S3. Additional computational details of small models

*QM and MM geometry optimizations of dinucleotide monophosphate models*

Dinucleotide monophosphate models were cut from the four representative snapshots of MD simulation, i.e., the snapshots which include the four substates of the $G_{L4}$ nucleotide and $G_{S+1}$ phosphate as discussed in the main text. We performed QM (PBEh-3c) and MM (*ff*99bsc0$\chi_{OL3CP}$; see Methods in the main text) geometry optimizations of these dinucleotide monophosphate models (**Figure S5**). We aimed to compare the characteristic combinations of $G_{L4}$ nucleobase position and $G_{S+1}$ phosphate conformations (*planar-native*, *tilted-native*, *planar-flipped* or *tilted-flipped*), i.e., the energy differences among these states. QM and MM geometry optimizations either in gas phase or with implicit solvent (COSMO[2] and GB[3,4] for QM and MM, respectively) were performed using *Xopt*[5,6] coupled with TURBOMOLE V7.3[7,8] (QM) or AMBER 14[9] (MM). Dihedral restraints include backbone dihedrals α, β, γ, δ, ε, ζ and χ, ribose dihedrals $\nu_0$, $\nu_3$, $\nu_4$, and C4'-C3'-C2'-O2', and base dihedrals C5-N7-C8-N9, C6- C5-N7-C8, N1-C6-C5-N7, C2-N1-C6-C5, N3-C2-N1-C6, N1-C2-N3-C4, C2-N3-C4-N9, C2-N1-C6-O6 and N2-C2-N3-C4 with the restraining constant equal to 0.1 $E_h \cdot rad^{-2}$. Use of tight restraints instead of the common constraints in QM optimizations provides residual flexibility which helps to correctly relax complex biomolecular building blocks.[5,6] Results are shown in **Table S4**.

*Energy scans describing the 0BPh interaction*

Dimethyl-phosphate – methyl-guanine intermolecular complex

Small model of dimethyl-phosphate – methyl-guanine (**Figure S6A**) was extracted from QM/MM-optimized geometry of structure **6e** (native $G_{S+1}$ phosphate). The missing hydrogens were added by hand and the whole geometry was optimized at the PBEh-3c level using angle and dihedral restraints with the restraining constant equal to 1.0 and 0.1 $E_h \cdot rad^{-2}$, respectively (O5'-H8-C8 and C5'-O5'-H8 angles, and C3'-O3'-P-O5', O3'-P-O5'-C5', C3'-O3'-C6-N1, C5'-O5'-C8-N9 and O5'-P-C5-C4 dihedrals). Then, we performed a rigid-monomers QM (PBEh-3c) interaction-energy scan along the C8H8…O5' vector with step length 0.05 Å in range of 2.10 - 2.60 Å. The *bff*[10] program was then used to perform two MM interaction-energy scans with rigid monomers (using non-bonding terms of AMBER *ff* with the vdW modification of phosphate oxygens) using the same geometries. Partial charges were taken from AMBER 16 library files and the partial charges of methyl hydrogens were set to a value of 0.04248 to obtain a molecular charge equal to −1. One of the MM scans includes NBfix correction applied to the –H8…O5'– Lennard-Jones pair. The $R_{i,j}$ parameter for H5 and OR atom types was set to a value of 2.8808 Å, while the $e_{i,j}$ parameter was kept at its default value of 0.0505 kcal/mol. Reference QM calculations at the DLPNO-CCSD(T)/CBS level were performed at C8H8…O5' distances of 2.10, 2.20, 2.30, 2.40 and 2.50 Å.

O3'-methylated-guanosine-monophosphate monomer

The O3'-methylated-guanosine-monophosphate model (**Figure S6B**) was extracted from QM/MM optimized geometry of structure **6e** (native $G_{S+1}$ phosphate) and the missing hydrogens were added by hand. Gaussian 09[11] was used for relaxed MM (*ff*99bsc0$\chi_{OL3CP}$) conformation energy scan along the C8H8…O5' distance in range of 2.05–2.60 Å with step length of 0.05 Å. Geometry constraints were applied on $G_{S+1}$(C8)-$G_{S+1}$(H8)-$G_{S+1}$(O5') angle, and $G_{L4}$(C3')-$G_{L4}$(O3')-$G_{S+1}$(C3')-$G_{S+1}$(C1') and $G_{L4}$(C3')-$G_{S+1}$(C5')-$G_{S+1}$(C2')-$G_{S+1}$(C4) dihedrals. Partial charges were taken from AMBER 16 library files and the partial charges of hydroxyl oxygens were modified, i.e., O3'H from 0.43760 to 0.3525 and O2'H from 0.41860 to 0.3335, in order to obtain a molecular charge equal to −1. Then, we also performed QM (PBEh-3c) single point calculations on the MM-optimized geometries.

*Energy scans describing the $U_{L2}$(2'-OH)…$G_{L4}$(N7) H-bond*

A small guanine-ribose model was extracted from the initial geometry of structure **1**. Two different guanine-ribose models were prepared for interaction energy scans using the following geometry optimizations.

Both models were optimized by QM (PBEh-3c) in gas phase using *Xopt* with geometrical restraints to keep the correct orientation of monomers representing $U_{L2}$ ribose and $G_{L4}$ base. The geometrical restraints were applied on 11 dihedrals (ribose dihedrals $\nu_0$ and $\nu_3$, backbone dihedrals $\gamma$ and $\delta$, C4'-C5'-O5'-O5'H, C5'-O5'-O5'H-C6, O5'-O5'H-N1-C6, C4'-C3'-O3'-O3'H, C8-C3'-O3'-O3'H, N9-C8-O3'-O3'H and C4'-C3'-C5-C4) using restraining constant equal to 0.1 $E_h \cdot rad^{-2}$. O2'-O2'H-N7 angle (representing the $U_{L2}$(O2')-$U_{L2}$(O2'H)-$G_{L4}$(N7) angle) was equal to 170° after the optimization of the first model. For the second guanine-ribose model, an additional geometrical restraint with the restraining constant equal to 0.1 $E_h \cdot rad^{-2}$ was applied on the O2'-O2'H-N7 angle in order to keep it at 143°, i.e., to maintain the lower directionality of the $U_{L2}$(2'-OH)…$G_{L4}$(N7) H-bond often found in MD simulations. Interaction-energy scans along the O2'…N7 vector were then performed with step length of 0.05 Å in range of 2.65 – 3.15 Å for QM (PBEh-3c) and MM (AMBER *ff*), and in range of 2.70 – 3.10 Å for QM reference (DLPNO-CCSD(T)/CBS).

*Geometry optimizations and energy scans of the $U_{L2}$ ribose*

The ribose model was excised from the initial geometry of structure **9**. The missing H atoms were added by hand to the O5'- and O3'-end. Then, the model was optimized by QM (PBEh-3c) or MM (*ff*99bsc0$_{CP}$) in gas phase. For QM optimization, *Xopt* was used with geometrical restraints on two dihedrals (C4'-C5'-O5'-O5'H and C4'-C3'-O3'-O3'H) with restraining constant equal to 0.1 $E_h \cdot rad^{-2}$ to keep the C2'-*endo* pucker of the $U_{L2}$ ribose. For MM optimization, Gaussian 09 with geometrical constraints on the same two dihedrals (C4'-C5'-O5'-O5'H and C4'-C3'-O3'-O3'H) was used while the LJ parameters and partial charges were taken from AMBER 16 library files. Two rigid conformation energy scans were performed on QM-optimized ribose, i.e. bending of the C2'-O2'-O2'H angle and rotation of the C1'-C2'-O2'-O2'H dihedral using MM (*ff*99bsc0$_{CP}$), QM (PBEh-3c) and QM reference (DLPNO-CCSD(T)/CBS) calculations. For MM optimization and single point calculations the partial charges were taken from nucleic10.in file, i.e., parameters for r-uracil with 5'-phosphate group and 3'-O$^-$ group, and the partial charges of the capping 5'- and 3'-end hydrogen atoms were set to value 0.1490 to obtain a neutral charge of the ribose. The H1'' hydrogen atom was assigned by H2 atom type to be equal to the H1’ atom and the H2-CT-H2 angle parameters were set to be equal to H1-CI-H1 angle parameters. The OR-HO bond parameters were set to OH-HO bond parameters, and the CT-OR-HO and CI-OR-HO angle parameters were set to be equal to CT-OR-P and CI-OR-P angle parameters.

*Energy scans describing sugar-base stacking*

A small ribose-guanine model for the stacking interaction (**Figure S6E**) was extracted from QM/MM-optimized geometry of structure **9e**. The model was optimized by QM (PBEh-3c) in gas phase using the *Xopt* program in the following way. Atoms of both monomers (except hydrogens and hydroxyl groups) were frozen during the optimization in order to keep the correct C2'-*endo* pucker of the $C_{L3}$ ribose and the mutual orientation of the $C_{L3}$ ribose and $G_{L4}$ base. Geometrical restraints were applied on three dihedrals involving the hydroxyl groups, i.e. O5'H-O5'-C5'-C4' (pseudo β), C4'-C3'-O3'-O3'H (pseudo ε) and C1'-C2'-O2'-O2'H dihedrals, using restraining constant equal to 0.1 $E_h \cdot rad^{-2}$ to keep the correct orientation of hydroxyl groups. Interaction energy scans were performed along the vector lying on the vertical axis linking the projection point of the O4' atom on the guanine plane and the O4' atom itself. Subsequently, interaction energies were calculated in range from 2.5 – 3.4 Å of the O4' – guanine distance, i.e., distance between the O4' atom and the guanine plane, with the step length of 0.1 Å. The PBEh-3c (QM), AMBER *ff* (MM) and DLPNO-CCSD(T) (QM reference) methods were used for energy calculations. Partial charges for MM calculations were derived as restrained electrostatic potential fit (RESP)[12] procedure by Antechamber.[13] Two types of RESP charges were used, i.e., AMBER-like partial charges derived from HF/6-31G* ESP (RESP-HF) and partial charges derived from B3LYP/def2-QZVP ESP (RESP-B3LYP). ESPs were computed with tight SCF setting using the Merz-Singh-Kollman population analysis by Gaussian 09. LJ parameters were taken from AMBER 16 library file nucleic10.in, i.e., parameters for r-cytosine with 5'-phosphate group and 3'-$O^-$ group for ribose and r-guanosine with 5'-phosphate group and 3'-$O^-$ group for guanine. The capping hydrogen atom on N9 atom of guanine was assigned by H atom type and the capping hydrogens on 3'- and 5'-end were assigned as HO atom types.

**S4. Comment on the comparison of 2’-OH group conformational scans using either $U_{L2}$ ribose or SPS model.** We performed gas phase conformation energy scans of the $U_{L2}$(C1’-C2’-O2’-O2’H) dihedral using C2’-*endo* ribose model and found that results are in agreement with conformation energy scans of the 2’-OH group rotation performed earlier by our group using the sugar-to-sugar (SPS) model (see ref. [14]). In the earlier study, the C3’-C2’-O2’-O2’H dihedral was analyzed using the gas phase and implicit solvent QM and MM calculations for various conformational families[15] of the SPS model.[14] We chose scans performed for 2[ conformational family for a comparison with our scan since the $U_{L2}$-$C_{L3}$ dinucleotide of the UUCG TL possesses the same conformational family.[15] Firstly, we compare the two conformational scans performed in gas phase, i.e. our ribose scan (Figure 7 in the main text) and the earlier SPS scan (see **Figure S1** in the Supporting information of ref. 14). Obviously, use of different models causes some differences between the two scans, i.e., QM energy is steeply descending towards the 260° in the SPS scan instead of finding a very flat QM minimum observed by our ribose scan. However, the steepness is still smaller compared to MM. Furthermore, in the SPS scans with implicit solvent (see Figure 6 of ref. [14]) MM finds a local minimum in the region of our interest, i.e. C1’-C2’-O2’-O2’H dihedral values between ~170° and ~200°, which corresponds to the C3’-C2’-O2’-O2’H dihedral region between ~290° and ~320°. However, the QM energy is increasing in the same region indicating that a shift of the dihedral towards larger values is unlikely. In contrast, shifting of the dihedral towards larger or smaller values on the quite symmetric local minimum of the MM scan indicates no preference for any of the directions. Taking all together, all these conformation energy scans, i.e. the new scans on the C2’-*endo* ribose model present in this study and the earlier scans on the SPS model,[14] show a difference in MM and QM description of the C1’-C2’-O2’-O2’H dihedral in the region of our interest suggesting that MM is prone to shifting the C1’-C2’-O2’-O2’H dihedral towards larger values more than QM. Obviously, as all calculations are done on small model systems, the results should not be over-interpreted.

**S5. MD Simulations and QM/MM and MM optimizations of the kink-turn Kt-7.** The sequence of the kink-turn 7 (Kt-7) from H. marismortui ribosome is close to the consensual sequence shared by all kink-turns.[16] Kt-7 is thus a useful model for studying the behavior of this recurrent RNA motif in calculations. Structurally, the kink-turn structure facilitates a sharp, almost perpendicular bend (kink), between two A-RNA helices. It is composed of canonical and non-canonical stems, and a bulge. The non-canonical stem contains conserved trans-Hoogsteen/sugar-edge A/G base pairs. The bent shape of the kink-turn structure is stabilized by an A-minor RNA tertiary interaction in which the adenine base from one of the A/G base pairs of the non-canonical stem makes contacts with the minor groove of the canonical stem. Furthermore, another adenine from the non-canonical stem forms a characteristic interaction with the first nucleotide of the bulge. The multitude of non-canonical but critically important interactions makes the kink-turn an ideal structure for benchmarking *ff* performance.[17]

Here, we focused our attention on two hydrogen bond interactions in which the 2-OH’ hydroxyl group of the ribose is acting as H-bond donor towards a base nitrogen. To simplify the presentation, we denote these H-bonds as HB1 and HB2, respectively. We show in this work (see the main text) that for the UNCG TL, comparison with the QM/MM calculations reveals excessive non-planarity predicted by the *ff* for a key H-bond involving the 2-OH’ hydroxyl group as a donor and nitrogen of the base as acceptor. Note that the acceptor atom in the UNCG H-bond is N7 while the HB1 and HB2 interactions involve the N1 atom. However, both of these atoms are described by the same vdW parameter in the *ff* and may thus be affected by similar issues.

We first performed five MD simulations of the Kt-7, each five microseconds long. The simulations indicated a generally satisfactory performance with the standard RNA force field *ff*99bsc0$\chi_{OL3}$ in combination with the SPC/E water model. Namely, the kink-turn maintained its bent shape and characteristic non-canonical interactions in all simulations, including the HB1 and HB2 H-bonds. The exception was two simulations in which we observed permanent transition from the A-minor I type of interaction towards the A-minor 0. This transition can be commonly observed in MD simulations of isolated kink-turns as both A-minor interactions are, in principle, possible for the kink-turn structure.[18,19] We subsequently selected three snapshots from the simulation ensemble and performed QM/MM and MM optimizations of these structures. As shown in **Table S9**, the QM/MM did not, in this case, reveal any problem with non-planarity comparable to the one seen for the UNCG TL. The interaction angle indicated by the MM was comparable with the QM/MM values.

*Computational details about QM/MM and MM optimizations of Kt-7*

The three MD snapshots (denoted as structures **1**, **2** and **3**) chosen for the QM/MM and MM optimizations of Kt-7 were solvated by water droplet with radius of ~40 Å from the center of RNA using SPC/E model and by 17 $K^+$ ions to neutralize the RNA using the tLEaP module of AMBER 16 software package. Then, equilibrations of water and ions (> 15 ps) were performed to relax positions of solvent molecules and to obtain a solvent distribution with the ions distributed at least 5 Å far from the QM region (structures **1a**, **2a** and **3a**). For two of the three solvent-equilibrated MD snapshots (structures **2** and **3**), another two snapshots were prepared for QM/MM and MM optimizations, i.e., structures with 60 ps and 100 ps-long water equilibrations (denoted as **b** and **c**). The whole RNA except of the two base pairs at the end of the canonical stem and one base pair at the end of the non-canonical stem was included in the QM region. All the other atoms including water and ions were included in the MM region. At the QM/MM interface the C4'-C5' bonds were cut, and the remaining carbon atoms were capped by hydrogen link atoms. The QM/MM and MM optimizations were performed as described in the main text without usage of the CP van der Waals correction for any of the structures.

**S6. Large-scale implicit solvent QM calculations.** In addition to QM/MM and MM calculations performed in the explicit solvent, we also tested the effect of implicit solvation for QM and MM geometry optimizations of the r(gcUUCGgc) TL. We used the COSMO implicit solvent at the QM level. The QM/COSMO computations are compared with equivalent MM/GB computations. We optimized multiple structural states and observed that the $G_{S+1}$(C8H8)…$G_{S+1}$(O5') 0BPh interaction is described differently by the QM and MM approaches (distance difference of ~0.12 Å, **Table S12**). We also identified that both signature sugar-base H-bonds, i.e., $U_{L2}$(2'-OH)…$G_{L4}$(N7) and $U_{L1}$(2'-OH)…$G_{L4}$(O6), are weakened by MM. The former one is weakened within MM by the decreased O-H…N angle (by ~28°). The latter one revealed prolonged donor-acceptor $U_{L1}$(O2')…$G_{L4}$(O6) and hydrogen-acceptor $U_{L1}$(O2'H)…$G_{L4}$(O6) distances (by ~0.04 Å and 0.03 Å, respectively) in MM-optimized TLs in comparison with QM-optimized TLs. Interestingly, MM-optimized geometries indicate that the $G_{L4}$ nucleotide prefers the *planar* conformation, whereas QM optimizations revealed *tilted* states (**Table S13** and **Figure S5**). Such a behavior could be connected to the already known overstabilization of stacking by current RNA *ff*s.[20–22] The tendency of MM to support *planar* position of the $G_{L4}$ nucleotide is, probably, further weakening the wrongly described $U_{L2}$(2'-OH)…$G_{L4}$(N7) and $U_{L1}$(2'-OH)…$G_{L4}$(O6) H-bonds. The result is consistent with the QM/MM and MM explicit-solvent investigations discussed in the main text. Note that implicit solvent optimizations resulted (in some optimizations) into formation of spurious molecular interactions and such structures were excluded from the analysis. Similar problems in continuum-solvent optimization of larger nucleic acids models were discussed by us elsewhere, where we have tried large-scale QM optimization of nucleic acids building blocks.[23] Generally, we assume that the QM/MM approach is more representative than the QM/COSMO approach for the purpose of this study, though all approaches provide a consistent picture.

**S7. Details about the relationship between $G_{S+1}$ phosphate conformation and positioning of the $G_{L4}$ nucleotide.** The primary NMR data[24] is consistent with two marginally different conformations (states) of the $G_{L4}$ *syn* nucleotide, namely a *planar state* with parallel position to the plane formed by the closing $G_{S+1}C_{S-1}$ base pair or a slightly *tilted state* (**Figure S5**). Based on the visual inspection of MD snapshots, we suspected that the $G_{L4}$ conformation (*planar* or *tilted*) could be related to $G_{S+1}$ phosphate flipping. Thus, we analyzed correlation between the $G_{L4}$ position and conformations of the $G_{S+1}$ phosphate and found that all $G_{L4}$-$G_{S+1}$ substates, i.e. having *native/flipped* $G_{S+1}$ phosphate state combined with *planar/tilted* $G_{L4}$ conformation, were populated during MD simulations (**Figure S8**). However, it is difficult to differentiate between *planar* and *tilted* conformation of the $G_{L4}$ nucleotide during MD simulations due to smooth fluctuations between them (angle between planes in the range ~10° and ~30°, respectively, see Methods in the main text for details about the angle analysis).

In order to get insights into the *ff* performance, we carried out a series of QM investigations. We extracted the dinucleotide-monophosphate model (containing the $G_{L4}$ nucleotide and the $G_{S+1}$ phosphate; **Figure S5**) out of the UNCG TL structure. We compared QM and MM relative potential energies of the four major conformational states of the $G_{L4}$ nucleotide - $G_{S+1}$ phosphate motif, i.e., *planar-native, planar-flipped, tilted-native,* and *tilted-flipped* (see **Section S3** for computational details). The idea to use this simple model was to exclude additional structural constraints coming from neighboring nucleotides and from the sugar-phosphate backbone. Both QM and MM indicate that *native* $G_{S+1}$ phosphate state is preferred independently on the $G_{L4}$ conformation (**Table S4**). However, *native* $G_{S+1}$ phosphate appears to favor *planar* $G_{L4}$ conformation in MM, which is in contrast with the QM description that has no preference (**Table S4**).

In summary, the analysis indicates that the *native* state of $G_{S+1}$ phosphate could drive $G_{L4}$ nucleotide to the *planar* conformation during MD simulations (due to its overstabilization within MM in comparison with the QM method). This conformation is prone to weakening of the signature $U_{L2}$(2'-OH)…$G_{L4}$(N7) sugar-base H-bond (**Figure S9**). In MD simulations, we did not see any correlation between position of the $G_{L4}$ nucleotide and the $G_{S+1}$ phosphate conformation (**Figure S8**).

**S8. Details about folding of the gcUUCGgc TL using the gHBfix potential.** In our previous work we have shown that the $\chi_{OL3CP}$ + gHBfix19 *ff* version generally improves RNA simulations without side-effects, though it was still not sufficient to fold the UNCG TL.[17]

We have then tried several additional gHBfix settings to specifically improve folding of the gcUUCGgc 8-mer. Support of sugar-base interactions appears necessary, as it targets important UNCG signature interactions. gHBfix19 modified by additional strengthening of sugar-base donor-acceptor interactions (i.e., –OH…N– and –OH…O– H-bonds) by 0.5 kcal/mol increased the population of structures with correctly folded A-form stem to ~70%.[17] However, the TL was still not properly structured, as, among other things, some other non-native sugar-base interactions occurred. Thus we tried to further weaken base-sugar donor – acceptor (i.e., –NH…O2'–) and sugar-sugar (i.e., –OH…O4'– and –OH…OH–) interactions (both by 0.5 kcal/mol). This resulted in a complex variant marked as $gHBfix\binom{2-OH...nbO/bO}{-0.5}\binom{NH...N}{+0.5}\binom{NH...O}{+0.5}\binom{2-OH...N/O}{+0.5}\binom{NH...O2}{-0.5}\binom{2-OH...O2/O4}{-0.5}$ when using notation of the original gHBfix paper.[17] For the sake of clarity, **Table S14** shows full list of all interactions affected by this modification which is abbreviated as $gHBfix_{UNCG19}$.

With the gHBfix$_{UNCG19}$ variant, population of the stem dropped to just ~12%, which is probably caused by some undesired side-effects within the unfolded ensemble. However, the overall population of fully correct UUCG TL was ~5 % in the reference replica of the 10 μs-long REST2 simulation. Considering possible convergence uncertainties of REST2 simulations,[17] we now prolonged the REST2 simulation towards 20 μs but we did not observe another complete folding event (both stem and loop correctly folded, **Figure S19**).

**S9. Details about geometry restraints of the $U_{L1}$(2'-OH) dihedral during QM/MM optimizations.** Parameters setting for geometry restraints on the $U_{L1}$(C1'-C2'-O2'-O2'H) dihedral includes R, i.e., the target value of the dihedral restraint, varying in range of 60°-190° with a step length of 10° along the scan, $r_1$ set to R−10°, $r_2$ and $r_3$ equal to R, $r_4$ set to R+10°, and $r_{k2}$ and $r_{k3}$ force constants set to 60 kcal·mol$^{-1}$·rad$^2$. Such a setting results into a two-sided parabolic-well potential restraint with well depth of 60 kcal·mol$^{-1}$·rad$^2$ in the range from R−10° to R+10° with linear extension outside of this range (for more information see the AMBER manual).

**S10. Details about the $U_{L1}$(C1'-C2'-O2'-O2'H) dihedral restraint applied during one set of REST2 folding simulations.** We applied the one-sided flat-well potential restraint, so that the C1'-C2'-O2'-O2'H dihedral was unrestrained below 115°, harmonic restraining potential with a force constant of 10 kcal/mol·rad$^2$ was applied from 115° to 135°, and linear extension of this potential was applied for dihedral values below above 135°.

### S11. Supporting Tables

**Table S1.** Comparison of the $G_{S+1}$(O4')…$G_{S+1}$(O5'), i.e., O…O distances (in Å) between UNCG TLs after QM/MM and MM optimizations, and QM/MM → MM re-optimizations.[a]

| | Initial value | QM/MM | MM | QM/MM →MM | $\Delta d_1$ [b] | $\Delta d_2$ [c] |
|---|---|---|---|---|---|---|
| structure **6** – water equilibration structures **6a-6e** | | | | | | |
| **6a** | 2.74 | 2.81 | 2.83 | 2.86 | 0.02 | 0.05 |
| **6b** | 2.74 | 2.73 | 2.79 | 2.78 | 0.06 | 0.05 |
| **6c** | 2.74 | 2.77 | 2.79 | 2.82 | 0.02 | 0.05 |
| **6d** | 2.74 | 2.77 | 2.84 | 2.80 | 0.07 | 0.03 |
| **6e** | 2.74 | 2.78 | 2.83 | 2.86 | 0.05 | 0.08 |
| average | | | | | 0.04 | 0.05 |
| structure **7** – water equilibration structures **7a-7e** | | | | | | |
| **7a** | 2.70 | 2.66 | 2.72 | -- | 0.06 | -- |
| **7b** | 2.70 | 2.69 | 2.77 | -- | 0.08 | -- |
| **7c** | 2.70 | 2.70 | 2.76 | -- | 0.07 | -- |
| **7d** | 2.70 | 2.75 | 2.79 | -- | 0.04 | -- |
| **7e** | 2.70 | 2.75 | 2.84 | -- | 0.08 | -- |
| average | | | | | 0.07 | |
| structure **9** – water equilibration structures **9a-9e** | | | | | | |
| **9a** | 2.57 | 2.60 | 2.62 | 2.67 | 0.02 | 0.07 |
| **9b** | 2.57 | 2.62 | 2.64 | 2.67 | 0.02 | 0.05 |
| **9c** | 2.57 | 2.55 | 2.62 | 2.60 | 0.07 | 0.05 |
| **9d** | 2.57 | 2.62 | 2.68 | 2.70 | 0.06 | 0.08 |
| **9e** | 2.57 | 2.62 | 2.68 | 2.67 | 0.06 | 0.05 |
| average[d] | | | | | 0.05 | 0.06 |

[a] Only structures with the native $G_{S+1}$ phosphate, where 0BPh interaction is established, are listed. In the main text, only the results for structures **6e**, **7e** and **9e** are shown. Note that the QM/MM→MM re-optimizations were performed only for some structures as the results showed similar trend as the MM optimizations. [b] $\Delta d_1$ = MM − QM/MM, [c] $\Delta d_2$ = (QM/MM→MM) – QM/MM, [d] structure **9b** is excluded from the average calculation as flip of the $G_{S+1}$ phosphate occurred during the QM/MM optimization.

**Table S2.** Comparison of the $G_{S+1}$(C8H8)…$G_{S+1}$(O5'), i.e., H…O distances (in Å) between UNCG TLs after QM/MM and MM optimizations, and QM/MM → MM re-optimizations.[a]

| | Initial value | QM/MM | MM | QM/MM →MM | $\Delta d_1$ [b] | $\Delta d_2$ [c] |
|---|---|---|---|---|---|---|
| structure **6** – water equilibration structures 6a-6e | | | | | | |
| **6a** | 2.18 | 2.21 | 2.36 | 2.29 | 0.15 | 0.08 |
| **6b** | 2.18 | 2.18 | 2.31 | 2.32 | 0.13 | 0.14 |
| **6c** | 2.18 | 2.14 | 2.34 | 2.30 | 0.20 | 0.16 |
| **6d** | 2.18 | 2.14 | 2.38 | 2.31 | 0.24 | 0.17 |
| **6e** | 2.18 | 2.08 | 2.28 | 2.24 | 0.20 | 0.16 |
| average | | | | | 0.18 | 0.14 |
| structure **7** – water equilibration structures 7a-7e | | | | | | |
| **7a** | 2.39 | 2.39 | 2.52 | -- | 0.12 | -- |
| **7b** | 2.39 | 2.33 | 2.42 | -- | 0.08 | -- |
| **7c** | 2.39 | 2.37 | 2.46 | -- | 0.09 | -- |
| **7d** | 2.39 | 2.30 | 2.42 | -- | 0.13 | -- |
| **7e** | 2.39 | 2.39 | 2.48 | -- | 0.09 | -- |
| average | | | | | 0.10 | |
| structure **9** – water equilibration structures 9a-9e | | | | | | |
| **9a** | 2.15 | 2.19 | 2.27 | 2.24 | 0.08 | 0.05 |
| **9b** | 2.15 | 2.92 | 2.24 | 2.92 | -0.68 | 0.00 |
| **9c** | 2.15 | 2.11 | 2.28 | 2.25 | 0.17 | 0.14 |
| **9d** | 2.15 | 2.30 | 2.35 | 2.30 | 0.05 | 0.00 |
| **9e** | 2.15 | 2.21 | 2.30 | 2.30 | 0.09 | 0.09 |
| average[d] | | | | | 0.10 | 0.07 |

[a] Only structures with the native $G_{S+1}$ phosphate, where 0BPh interaction is established, are listed. In the main text, only the results for structures **6e**, **7e** and **9e** are shown. Note that the QM/MM→MM re-optimizations were performed only for some structures as the results showed similar trend as the MM optimizations. [b] $\Delta d_1$ = MM − QM/MM, [c] $\Delta d_2$ = (QM/MM→MM) – QM/MM, [d] structure **9b** is excluded from the average calculation as flip of the $G_{S+1}$ phosphate occurred during the QM/MM optimization.

**Table S3.** Comparison of the $G_{S+1}$(C8H8)…$G_{S+1}$(O5'), i.e., H…O and $G_{S+1}$(O4') – $G_{S+1}$(O5'), i.e., O – O distances (in Å) between the two dinucleotide monophosphate models with the native $G_{S+1}$ phosphate (*planar-native* and *tilted-native*) optimized with or without geometrical restraints (restrained and free optimizations, respectively) by QM/COSMO or MM/GB (see **Section S3** for computational details).

| free optimizations | start | QM | $\Delta d_{QM}$ | MM | $\Delta d_{MM}$ | $\Delta d_{MM-QM}$ |
|---|---|---|---|---|---|---|
| planar-native | | | | | | |
| $G_{S+1}$(H8/O5') | 2.37 | 2.28 | -0.09 | 2.56 | 0.19 | 0.28 |
| $G_{S+1}$(O4'/O5') | 2.76 | 2.80 | 0.04 | 2.90 | 0.14 | 0.10 |
| tilted-native | | | | | | |
| $G_{S+1}$(H8/O5') | 2.18 | 2.29 | 0.11 | 2.56 | 0.38 | 0.27 |
| $G_{S+1}$(O4'/O5') | 2.74 | 2.80 | 0.06 | 2.90 | 0.16 | 0.10 |

| restrained optimizations | start | QM | $\Delta d_{QM}$ | MM | $\Delta d_{MM}$ | $\Delta d_{MM-QM}$ |
|---|---|---|---|---|---|---|
| planar-native | | | | | | |
| $G_{S+1}$(H8/O5') | 2.37 | 2.23 | -0.14 | 2.50 | 0.13 | 0.27 |
| $G_{S+1}$(O4'/O5') | 2.76 | 2.71 | -0.05 | 2.87 | 0.11 | 0.16 |
| tilted-native | | | | | | |
| $G_{S+1}$(H8/O5') | 2.18 | 2.22 | 0.04 | 2.44 | 0.26 | 0.22 |
| $G_{S+1}$(O4'/O5') | 2.74 | 2.74 | 0.00 | 2.80 | 0.06 | 0.06 |

**Table S4.** Relative QM and MM energies (in kcal/mol) of the four dinucleotide monophosphate models (*planar-native*, *tilted-native*, *planar-flipped* and *tilted-flipped*) optimized with geometrical restraints on dihedrals using QM (PBEh-3c) or MM (*ff*99bsc0$\chi_{OL3CP}$) method with COSMO and GB implicit solvent, respectively.[a]

| | QM optimized | | MM optimized | |
|---|---|---|---|---|
| $G_{L4}$-$G_{S+1}$ | $\Delta E_{conf,QM}$ | $\Delta E_{conf,MM}$ | $\Delta E_{conf,QM}$ | $\Delta E_{conf,MM}$ |
| implicit solvent single point calculations | | | | |
| *planar-native* | 0.78 | 0.00 | 1.57 | 0.00 |
| *planar-flipped* | 3.90 | 8.00 | 5.71 | 7.71 |
| *tilted-native* | 0.00 | 3.26 | 0.00 | 3.64 |
| *tilted-flipped* | 2.17 | 3.53 | 0.61 | 3.98 |
| gas phase single point calculations | | | | |
| *planar-native* | 5.40 | 2.53 | 5.88 | 1.84 |
| *planar-flipped* | 5.81 | 9.68 | 9.95 | 10.58 |
| *tilted-native* | 0.00 | 0.00 | 0.00 | 0.00 |
| *tilted-flipped* | 10.90 | 10.04 | 9.33 | 11.73 |

[a] See **Section S0** for computational details and dihedral restraints. The most stable structure in each column has energy 0 kcal/mol. Single point energy calculations are performed either with implicit solvent or in gas phase.

**Table S5.** Comparison of the O…N and H…N distances, i.e., donor-acceptor and hydrogen-acceptor distances (in Å, separated by "/") of the $U_{L2}$(2'-OH)…$G_{L4}$(N7) H-bond in UNCG TLs after QM/MM and MM optimizations, and QM/MM → MM re-optimizations.[a]

| structure | initial value | QM/MM | MM | QM/MM →MM | Δd1 [b] | Δd2 [c] |
|---|---|---|---|---|---|---|
| **1** | 2.89/2.06 | 2.86/1.92 | 2.80/1.92 | 2.82/1.93 | -0.06/-0.01 | -0.04/ 0.01 |
| **2** | 2.73/2.14 | 2.72/1.76 | 2.78/1.97 | 2.73/1.86 | 0.06/ 0.21 | 0.01/ 0.10 |
| **4** | 2.93/2.12 | 2.76/1.84 | 2.82/2.11 | 2.79/2.00 | 0.06/ 0.27 | 0.03/ 0.16 |
| structure **5** – water equilibration structures 5a-5e | | | | | | |
| **5a** | 2.75/1.83 | 2.82/1.87 | 2.89/2.03 | 2.80/1.86 | 0.07/ 0.15 | -0.02/-0.01 |
| **5b** | 2.75/1.83 | 2.79/1.83 | 2.82/1.90 | 2.76/1.83 | 0.03/ 0.07 | -0.03/ 0.00 |
| **5c** | 2.75/1.83 | 2.76/1.78 | 2.88/1.94 | 2.74/1.78 | 0.12/ 0.16 | -0.02/ 0.00 |
| **5d** | 2.75/1.83 | 2.80/1.82 | 2.77/1.82 | 2.75/1.81 | -0.03/-0.01 | -0.05/-0.01 |
| **5e** | 2.75/1.83 | 2.88/1.95 | 2.89/1.96 | 2.83/1.90 | 0.02/ 0.01 | -0.05/-0.05 |
| average | | | | | 0.04/0.08 | -0.03/-0.01 |
| structure **6** – water equilibration structures 5a-5e | | | | | | |
| **6a** | 2.71/1.80 | 2.71/1.73 | 2.72/1.75 | 2.70/1.72 | 0.01/ 0.02 | -0.01/-0.01 |
| **6b** | 2.71/1.80 | 2.80/1.85 | 2.78/1.93 | 2.76/1.89 | -0.02/ 0.08 | -0.04/ 0.04 |
| **6c** | 2.71/1.80 | 2.68/1.72 | 2.84/2.07 | 2.68/1.78 | 0.17/ 0.35 | 0.00/ 0.06 |
| **6d** | 2.71/1.80 | 2.77/1.82 | 2.86/2.11 | 2.74/1.85 | 0.09/ 0.29 | -0.03/ 0.03 |
| **6e** | 2.71/1.80 | 2.98/2.03 | 2.95/2.07 | 2.91/1.98 | -0.03/ 0.04 | -0.07/-0.05 |
| average | | | | | 0.04/0.16 | -0.03/0.01 |
| structure **9** – water equilibration geometries a-e | | | | | | |
| **9a** | 2.72/1.88 | 2.72/1.75 | 2.74/1.86 | 2.69/1.76 | 0.03/ 0.11 | -0.03/ 0.01 |
| **9b** | 2.72/1.88 | 2.70/1.74 | 2.74/1.89 | 2.69/1.79 | 0.04/ 0.15 | -0.01/ 0.05 |
| **9c** | 2.72/1.88 | 2.84/1.86 | 2.74/1.89 | 2.81/1.84 | -0.10/ 0.03 | -0.03/-0.02 |
| **9d** | 2.72/1.88 | 2.83/1.88 | 2.80/1.91 | 2.80/1.91 | -0.03/ 0.04 | -0.03/ 0.03 |
| **9e** | 2.72/1.88 | 2.82/1.86 | 2.80/1.96 | 2.75/1.85 | -0.03/ 0.10 | -0.07/-0.01 |
| average | | | | | -0.02/0.09 | -0.03/0.01 |

[a] Each column contains two values separated by a slash, the first number represents O…N donor-acceptor distance and the second number represents H…N hydrogen-acceptor distance. Note that structure **3** is not included in this analysis because $U_{L2}$(2'-OH) forms an alternative H-bond with $G_{L4}$(O6) instead of $G_{L4}$(N7) atom. Structures **7** and **8** are not included in this analysis because spurious state of the $U_{L1}$(2'-OH) group is present in these structures, which may possibly influence position of the $G_{L4}$ nucleobase and subsequently the $U_{L2}$(2'-OH)…$G_{L4}$(N7) H-bond. [b] $\Delta d_1$ = MM – QM/MM, [c] $\Delta d_2$ = (QM/MM→MM) – QM/MM.

**Table S6.** Comparison of the O-H…N angle of the $U_{L2}$(2'-OH)…$G_{L4}$(N7) H-bond (in °) in UNCG TLs after QM/MM and MM optimizations, and QM/MM → MM re-optimizations.[a]

| Structure | initial value | QM/MM | MM | QM/MM →MM | $\Delta a_1$ [b] | $\Delta a_2$ [c] |
|---|---|---|---|---|---|---|
| **1** | 143.4 | 159.5 | 148.6 | 149.7 | -10.9 | -9.8 |
| **2** | 118.1 | 163.3 | 138.9 | 146.5 | -24.4 | -16.7 |
| **4** | 140.8 | 155.2 | 128.2 | 135.9 | -27.0 | -19.3 |
| structure **5** – water equilibration structures **5a-5e** | | | | | | |
| **5a** | 161.7 | 161.6 | 146.8 | 159.3 | -14.8 | -2.3 |
| **5b** | 161.7 | 166.3 | 157.5 | 157.9 | -8.8 | -8.3 |
| **5c** | 161.7 | 168.2 | 160.2 | 163.9 | -8.0 | -4.3 |
| **5d** | 161.7 | 171.0 | 164.2 | 159.0 | -6.7 | -11.9 |
| **5e** | 161.7 | 158.6 | 159.8 | 158.5 | 1.2 | -0.1 |
| Average | | | | | -7.4 | -5.4 |
| structure **6** – water equilibration structures **6a-6e** | | | | | | |
| **6a** | 157.0 | 172.4 | 171.8 | 172.5 | -0.6 | 0.1 |
| **6b** | 157.0 | 162.2 | 143.6 | 145.3 | -18.6 | -16.9 |
| **6c** | 157.0 | 162.0 | 135.8 | 150.7 | -26.2 | -11.3 |
| **6d** | 157.0 | 160.7 | 132.6 | 148.1 | -28.1 | -12.6 |
| **6e** | 157.0 | 162.5 | 150.0 | 158.1 | -12.5 | -4.4 |
| Average | | | | | -17.2 | -9.0 |
| structure **9** – water equilibration structures **9a-9e** | | | | | | |
| **9a** | 143.5 | 168.2 | 149.3 | 157.1 | -14.8 | -7.0 |
| **9b** | 143.5 | 176.2 | 144.3 | 151.2 | -19.2 | -12.3 |
| **9c** | 143.5 | 163.6 | 144.8 | 170.0 | -30.1 | -4.9 |
| **9d** | 143.5 | 158.1 | 149.7 | 149.6 | -14.3 | -14.4 |
| **9e** | 143.5 | 161.8 | 142.6 | 151.5 | -26.5 | -17.6 |
| Average | | | | | -21.0 | -11.2 |

[a] Note that structure **3** is not included in this analysis because $U_{L2}$(2'-OH) forms an alternative H-bond with $G_{L4}$(O6) instead of $G_{L4}$(N7) atom. Structures **7** and **8** are not included in this analysis because spurious state of the $U_{L1}$(2'-OH) group is present in these structures, which may possibly influence position of the $G_{L4}$ nucleobase and subsequently the $U_{L2}$(2'-OH)…$G_{L4}$(N7) H-bond. [b] $\Delta a_1$ = MM – QM/MM, [c] $\Delta a_2$ = (QM/MM→MM) – QM/MM. Note that both comparisons predict visible underestimation of the angle at the MM level of theory. Use of the QM/MM→MM optimized structures leads to a smaller difference than use of the MM optimized structures, for reasons that are in more detail commented on in ref. [25].

**Table S7.** Optimized values of $U_{L2}$ ribose C2'-O2'-O2'H angle and C1'-C2'-O2'-O2'H dihedral (in °). Starting ribose model for QM (PBEh-3c) and MM (*ff*99bsc0$_{CP}$) geometry optimizations was taken from structure **9**.

| | angle C2'-O2'-O2'H | dihedral C1'-C2'-O2'-O2'H |
|---|---|---|
| start (structure **9**) | 110.4 | 163.2 |
| PBEh-3c | 109.3 | 189.8 |
| *ff*99bsc0$_{CP}$ | 106.5 | 261.3 |

**Table S8.** Comparison of the $U_{L2}$ ribose C1'-C2'-O2'-O2'H dihedral (in °) of the QM/MM- and MM-optimized UNCG TLs.[a]

| structure | Initial value | QM/MM | MM | QM/MM→ MM | $\Delta a_1$[b] | $\Delta a_2$[c] |
|---|---|---|---|---|---|---|
| **1** | 210.9 | 180.7 | 186.6 | 182.7 | 5.9 | 2.0 |
| **2** | 181.0 | 171.4 | 187.9 | 181.5 | 16.5 | 10.1 |
| **4** | 203.3 | 183.2 | 192.9 | 193.2 | 9.7 | 10.0 |
| structure **5** – water equilibration structures **5a**-**5e** | | | | | | |
| **5a** | 158.1 | 166.5 | 167.3 | 171.7 | 0.8 | 5.2 |
| **5b** | 158.1 | 168.7 | 171.1 | 175.3 | 2.4 | 6.6 |
| **5c** | 158.1 | 160.0 | 178.2 | 170.3 | 18.2 | 10.3 |
| **5d** | 158.1 | 166.9 | 174.0 | 175.7 | 7.1 | 8.8 |
| **5e** | 158.1 | 143.9 | 150.7 | 146.9 | 6.8 | 3.0 |
| average | | | | | 7.1 | 6.8 |
| structure **6** – water equilibration structures **6a**-**6e** | | | | | | |
| **6a** | 193.8 | 167.8 | 171.4 | 173.8 | 3.6 | 6.0 |
| **6b** | 193.8 | 193.4 | 210.0 | 202.6 | 16.6 | 9.2 |
| **6c** | 193.8 | 175.8 | 170.8 | 184.1 | -5.0 | 8.3 |
| **6d** | 193.8 | 189.9 | 189.0 | 194.0 | -0.9 | 4.1 |
| **6e** | 193.8 | 171.6 | 183.0 | 179.6 | 11.4 | 8.0 |
| average | | | | | 5.1 | 7.1 |
| structure **9** – water equilibration structures **9a**-**9e** | | | | | | |
| **9a** | 175.8 | 168.2 | 164.2 | 170.0 | -4.0 | 1.8 |
| **9b** | 175.8 | 176.2 | 185.1 | 182.6 | 8.9 | 6.4 |
| **9c** | 175.8 | 163.6 | 206.8 | 173.1 | 43.2 | 9.5 |
| **9d** | 175.8 | 158.1 | 169.8 | 168.3 | 11.7 | 10.2 |
| **9e** | 175.8 | 161.8 | 181.6 | 173.9 | 19.8 | 12.1 |
| average | | | | | 15.9 | 8.0 |

[a] Note that structure **3** is not included in the table because $U_{L2}$(2'-OH) forms an alternative H-bond with $G_{L4}$(O6) instead of $G_{L4}$(N7) atom. Structures **7** and **8** are not included in the table because spurious state of the $U_{L1}$(2'-OH) group is present in these structures, which may possibly influence position of the $G_{L4}$ nucleobase and subsequently the $U_{L2}$(2'-OH)…$G_{L4}$(N7) H-bond. [b] $\Delta a_1$ = MM – QM/MM, [c] $\Delta a_2$ = (QM/MM→MM) – QM/MM

**Table S9.** Directionality of the HB1 and HB2 H-bonds in the RNA kink-turn 7, defined by the G(O2')-G(O2'H)-A(N1) angle (in °).[a]

| structure | MD average | initial value | QM/MM | MM | $\Delta d_{MM-QM/MM}$ |
|---|---|---|---|---|---|
| | **HB1** | | | | |
| **1a** | | 164.5 | 172.4 | 178.1 | 5.7 |
| **2a** | | 175.8 | 169.7 | 171.5 | 1.8 |
| **2b** | | 175.8 | 169.0 | 171.8 | 2.8 |
| **2c** | 161.2 | 175.8 | 166.8 | 171.2 | 4.4 |
| **3a** | | 161.7 | 167.5 | 166.2 | -1.3 |
| **3b** | | 161.7 | 168.7 | 168.8 | 0.1 |
| **3c** | | 161.7 | 168.8 | 164.2 | -4.5 |
| | **HB2** | | | | |
| **1a** | | 153.1 | 167.6 | 159.5 | -8.2 |
| **2a** | | 172.7 | 172.5 | 171.2 | -1.2 |
| **2b** | | 172.7 | 171.0 | 177.9 | 6.9 |
| **2c** | 163.8 | 172.7 | 172.3 | 174.9 | 2.6 |
| **3a** | | 162.6 | 172.3 | 168.5 | -3.7 |
| **3b** | | 162.6 | 170.9 | 172.3 | 1.5 |
| **3c** | | 162.6 | 173.9 | 176.6 | 2.6 |

[a] Average values in the MD simulations, the values in structures used as the initial geometries for the QM/MM and MM optimizations, and the resulting values of respective optimizations are presented. Structures **1**, **2**, and **3** represent three different RNA geometries taken from MD simulations and notations **a**, **b** and **c** represent three different water distributions introduced around the RNA for each structure. $\Delta d_{MM-QM/MM}$ column shows the difference between MM-optimized and QM/MM-optimized geometry and is calculated as MM − QM/MM.

**Table S10.** Population (in %) of the alternative $U_{L1}$(2'-OH)…$U_{L1}$(O5') H-bond, i.e. $U_{L1}$(2'-OH) base-phosphate flipping, in MD simulations using different water models (OPC, SPC/E and TIP3P) and salt concentration (0.15 M, 1.0 M and net-neutral).

| $G_{S+1}$ phosphate | 0.15M KCl | 0.15M KCl/CP[a] | 1.0M KCl/CP |
|---|---|---|---|
| Native | 89.4 | 85.2 | 79.6 |
| flipped | 10.6 | 14.8 | 20.4 |
| $G_{S+1}$ phosphate | 0.15M KCl | 0.15M KCl/CP | 1.0M KCl |
| Native | 78.6 | 75.5 | 72.2 |
| flipped | 21.4 | 24.5 | 27.8 |
| $G_{S+1}$ phosphate | 0.15M KCl | Net-neutral | 1.0M KCl |
| Native | 87.2 | 90.9 | 80.7 |
| flipped | 12.8 | 9.1 | 19.3 |

[a] CP = Case phosphates paramaters

**Table S11**. Sugar-base stacking between the $C_{L3}$ ribose and the $G_{L4}$ nucleobase represented by a distance between the $C_{L3}$(O4') oxygen atom and the $G_{L4}$ nucleobase plane (in Å) in the QM/MM- and MM-optimized UNCG TLs and a comparison between these methods.[a]

| structure | initial value | QM/MM | MM | $\Delta d_{QM/MM}$[b] | $\Delta d_{MM}$[c] | $\Delta d_1$[d] |
|---|---|---|---|---|---|---|
| **1** | 3.19 | 2.80 | 2.93 | -0.39 | -0.26 | 0.13 |
| **2** | 2.92 | 2.82 | 2.83 | -0.10 | -0.09 | 0.02 |
| **3** | 2.76 | 2.82 | 2.84 | 0.06 | 0.08 | 0.02 |
| **4** | 2.89 | 2.82 | 2.78 | -0.08 | -0.12 | -0.04 |
| structure **5** – water equilibration structures **5a**-**5e** | | | | | | |
| **5a** | 2.81 | 2.70 | 2.81 | -0.11 | 0.00 | 0.11 |
| **5b** | 2.81 | 2.86 | 2.79 | 0.05 | -0.02 | -0.07 |
| **5c** | 2.81 | 2.66 | 2.75 | -0.15 | -0.06 | 0.09 |
| **5d** | 2.81 | 2.76 | 2.80 | -0.05 | -0.01 | 0.05 |
| **5e** | 2.81 | 2.71 | 2.78 | -0.10 | -0.03 | 0.07 |
| average | | | | | | 0.05 |
| structure **6** – water equilibration structures **6a**-**6e** | | | | | | |
| **6a** | 2.95 | 2.84 | 2.86 | -0.12 | -0.10 | 0.02 |
| **6b** | 2.95 | 2.92 | 2.88 | -0.03 | -0.07 | -0.03 |
| **6c** | 2.95 | 2.81 | 2.86 | -0.14 | -0.09 | 0.05 |
| **6d** | 2.95 | 2.80 | 2.87 | -0.15 | -0.08 | 0.07 |
| **6e** | 2.95 | 2.90 | 2.91 | -0.05 | -0.04 | 0.01 |
| average | | | | | | 0.02 |
| structure **8** – water equilibration structures **8a**-**8e** | | | | | | |
| **8a** | 3.03 | 2.86 | 2.92 | -0.18 | -0.12 | 0.06 |
| **8b** | 3.03 | 2.84 | 2.85 | -0.19 | -0.18 | 0.01 |
| **8c** | 3.03 | 2.84 | 2.87 | -0.19 | -0.17 | 0.03 |
| **8d** | 3.03 | 2.99 | 3.01 | -0.05 | -0.02 | 0.03 |
| **8e** | 3.03 | 2.98 | 2.93 | -0.06 | -0.10 | -0.05 |
| average | | | | | | 0.02 |
| structure **9** – water equilibration structures **9a**-**9e** | | | | | | |
| **9a** | 2.87 | 2.82 | 2.92 | -0.05 | 0.05 | 0.09 |
| **9b** | 2.87 | 2.89 | 2.95 | 0.02 | 0.08 | 0.06 |
| **9c** | 2.87 | 2.92 | 2.89 | 0.05 | 0.02 | -0.03 |
| **9d** | 2.87 | 2.97 | 2.95 | 0.10 | 0.08 | -0.02 |
| **9e** | 2.87 | 2.84 | 2.84 | -0.04 | -0.03 | 0.01 |
| average | | | | | | 0.02 |

[a] Structure **7** had to be excluded from the analysis because of a different $U_{L1}$(2'-OH) group orientation between the QM/MM- and MM-optimized structures. [b] $\Delta d_{QM/MM}$ = QM/MM – initial value, [c] $\Delta d_{MM}$ = MM – initial value, [d] $\Delta d_1$ = MM – QM/MM.

**Table S12.** Comparison of QM- and MM-optimized UNCG TLs using implicit solvent calculations. The analyzed features are the $G_{S+1}$(C8H8)…$G_{S+1}$(O5') H…O distance describing 0BPh interaction, the $U_{L2}$(O2')-$U_{L2}$(O2'H)-$G_{L4}$(N7) O-H…N angle describing $U_{L2}$(2'-OH)…$G_{L4}$(N7) signature H-bond, and the $U_{L1}$(O2')-$G_{L4}$(O6) donor-acceptor and $U_{L1}$(O2'H)-$G_{L4}$(O6) hydrogen-acceptor distances describing $U_{L1}$(2'-OH)…$G_{L4}$(O6) signature H-bond. [a,b]

| | Start | QM | MM | MM − QM |
|---|---|---|---|---|
| $G_{S+1}$(C8H8)…$G_{S+1}$(O5') H…O distance (in Å)[c] | | | | |
| structure **6** | 2.18 | 2.29 | 2.45 | 0.16 |
| structure **7** | 2.39 | 2.31 | 2.39 | 0.08 |
| structure **9** | 2.15 | 2.27 | 2.39 | 0.12 |
| $U_{L2}$(O2')-$U_{L2}$(O2'H)-$G_{L4}$(N7) angle (in °)[d] | | | | |
| structure **1** | 143.4 | 167.7 | 139.3 | -28.4 |
| structure **2** | 118.1 | 167.8 | 137.5 | -30.3 |
| structure **4** | 140.8 | 167.1 | 139.7 | -27.4 |
| structure **5** | 161.6 | 167.6 | 139.6 | -28.0 |
| $U_{L1}$(2'-OH)…$G_{L4}$(O6) H-bond O…O/H…O distances (in Å)[e] | | | | |
| structure **1** | 2.61/1.70 | 2.64/1.68 | 2.69/1.71 | 0.05/0.03 |
| structure **2** | 2.64/1.79 | 2.64/1.68 | 2.68/1.70 | 0.04/0.03 |
| structure **4** | 2.74/1.80 | 2.65/1.69 | 2.68/1.70 | 0.02/0.01 |
| structure **5** | 2.70/1.76 | 2.64/1.68 | 2.69/1.71 | 0.04/0.03 |

[a] The PBEh-3c (QM) and ff99bsc0$\chi_{OL3}$ (MM) methods using implicit solvent (COSMO and GB for QM and MM, respectively) were used for geometry optimizations. [b] Structures **3**, **6** and **9** are not included in the analyses because of formation of the spurious $U_{L2}$(O2'H)…$C_{L3}$(O4') H-bond after MM/GB optimization. [c] Only structures with the native $G_{S+1}$ phosphate, i.e., possessing the 0BPh interaction, were used for this analysis. [d] Structures **7** and **8** are not included in this analysis because spurious state of the $U_{L1}$(2'-OH) group is present in these structures, which may possibly influence position of the $G_{L4}$ nucleobase and subsequently the $U_{L2}$(2'-OH)…$G_{L4}$(N7) H-bond. [e] Structures **7** and **8** are not included in this analysis because spurious state of the $U_{L1}$(2'-OH) group is present in these structures, i.e., the $U_{L1}$(2'-OH)…$G_{L4}$(O6) H-bond is not formed.

**Table S13.** The $G_{L4}$ nucleobase position relatively to $G_{S+1}$ nucleobase after QM (PBEh-3c) and MM (*ff*99bsc0$\chi_{OL3}$) geometry optimization in implicit solvent (COSMO and GB for QM and MM, respectively) for structures **1**, **2**, **4** and **5**. All these analyzed structures possess flipped $G_{S+1}$ phosphate. [a]

| structure | **1** | **2** | **4** | **5** |
|---|---|---|---|---|
| QM | tilted | tilted | tilted | tilted |
| MM | planar | planar | planar | planar |

[a] Structures **3**, **6** and **9** are not included in the analysis because of formation of a spurious $U_{L2}$(O2'H)…$C_{L3}$(O4') H-bond after MM/GB optimization. Structures **7** and **8** are not included in this analysis because spurious state of the $U_{L1}$(2'-OH) group is present in these structures, which may possibly influence orientation of the $G_{L4}$ nucleobase.

**Table S14.** List of interactions that were modified by the gHBfix$_{UNCG19}$ potential for the r(gcUUCGgc) TL.

| Interacting groups [a] | Bias (kcal/mol) | | H-bonds affected |
|---|---|---|---|
| | Support | Weakening | |
| NH…N | 0.5 | - | N1H/N2H/N3H/N4H…N3/N7 |
| NH…O | 0.5 | - | N1H/N2H/N3H/N4H…O2/O4/O6 |
| 2-OH…N | 0.5 | - | 2’-OH/3’-OH/5’-OH…N3/N7 |
| 2-OH…O | 0.5 | - | 2’-OH/3’-OH/5’-OH…O2/O4/O6 |
| 2-OH…bO | - | 0.5 | 2’-OH/3’-OH/5’-OH…O3’/O5’ |
| 2-OH…nbO | - | 0.5 | 2’-OH/3’-OH/5’-OH…*pro*-$R_P$/*pro*-$S_P$ |
| NH…O2 | - | 0.5 | N1H/N2H/N3H/N4H…2’-OH/3’-OH/5’-OH |
| 2-OH…O4 | - | 0.5 | 2’-OH/3’-OH/5’-OH…O4’ |
| 2-OH…O2 | - | 0.5 | 2’-OH/3’-OH/5’-OH…2’-OH/3’-OH/5’-OH |

[a] see Table 2 in ref. [17] for the definition of interacting groups

### S12. Supporting Figures

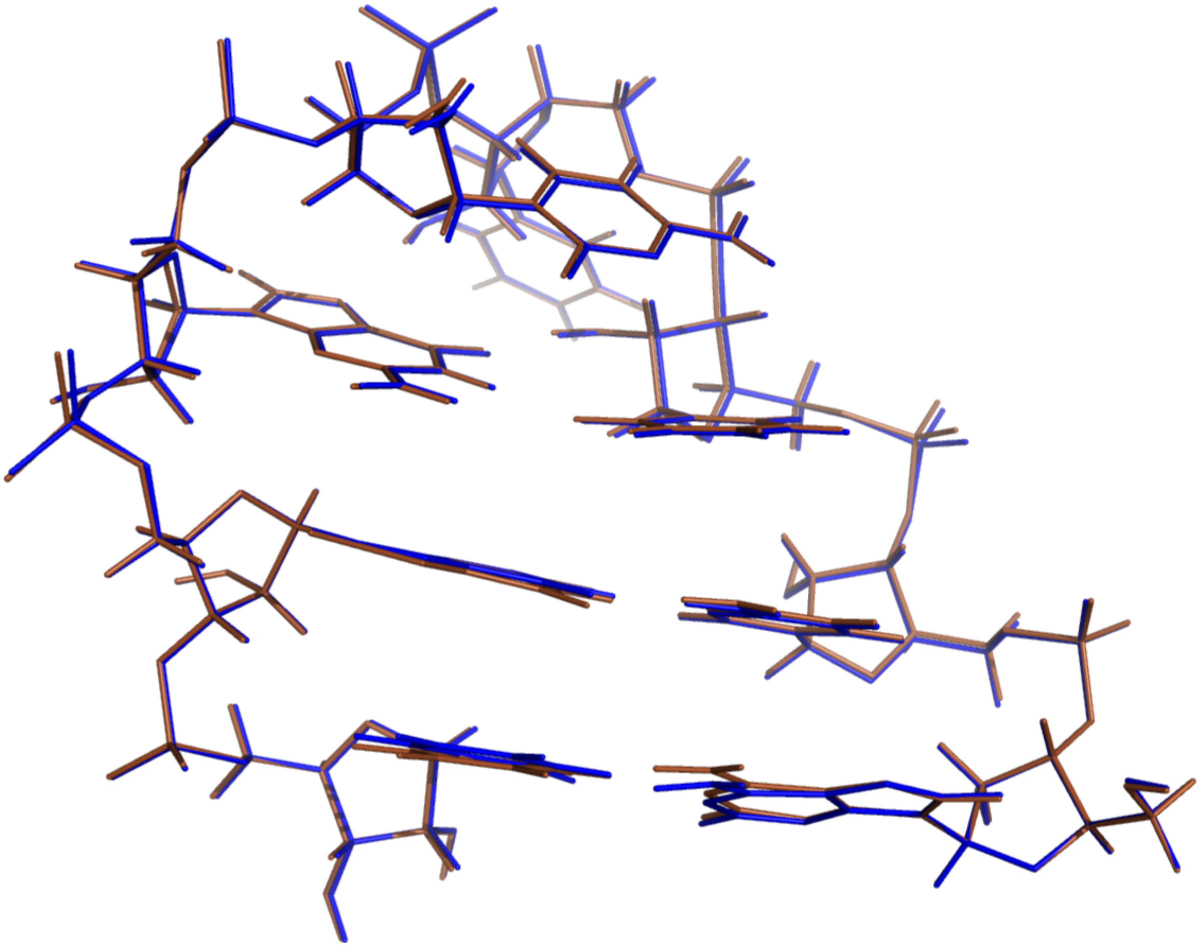

**Figure S1.** Overlay of the MM-optimized UNCG TL of structure **2** with or without vdW modification of phosphate oxygens (CP),[26] i.e. with CP (blue) and without CP (brown).

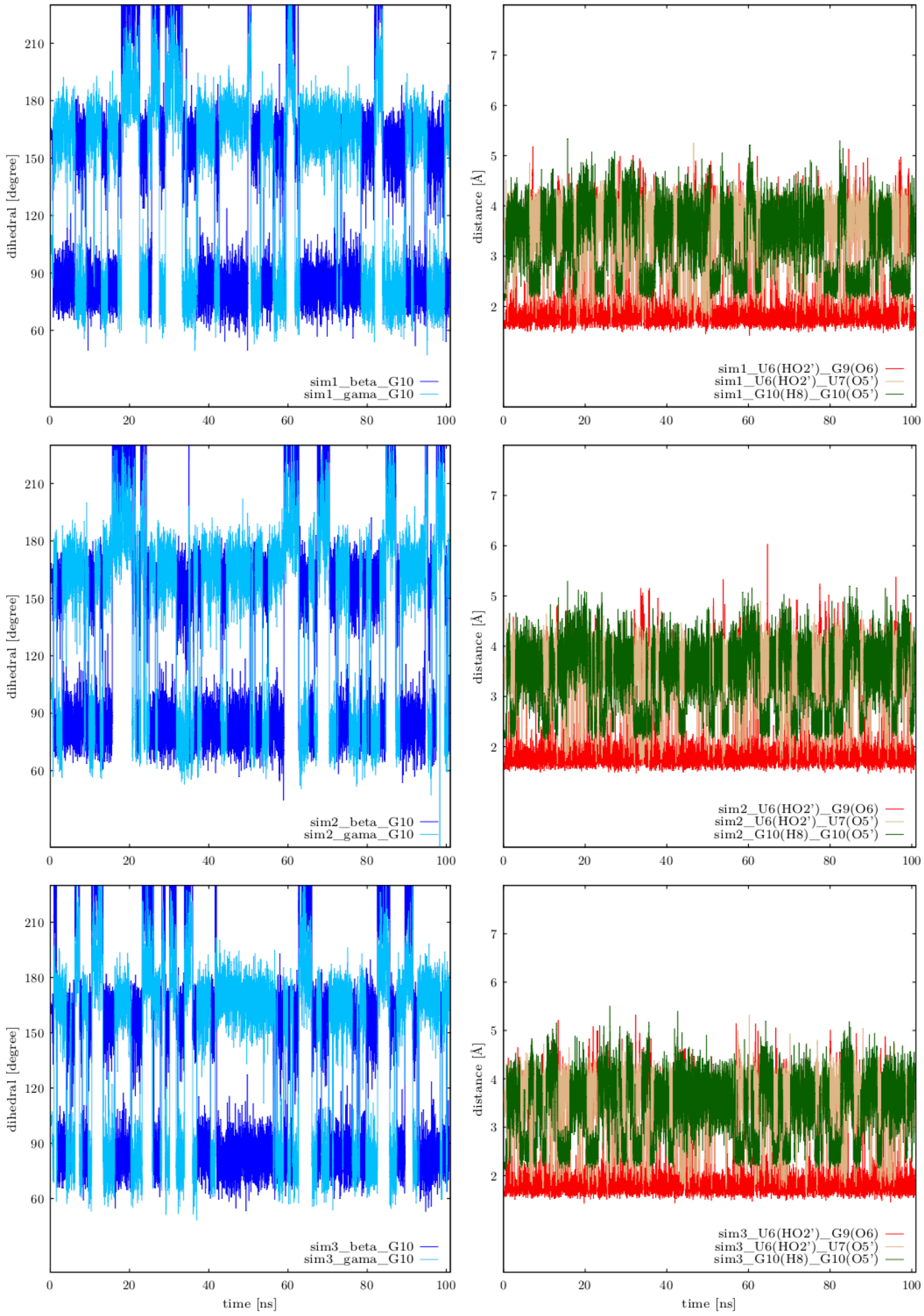

**Figure S2.** Occurrence of the alternative (flipped) $G_{S+1}$ phosphate state in three MD simulations (marked sim1, sim2, sim3) during the initial 100 ns and its possible correlation with the $U_{L2}$(2'-OH) flipping. Native state of the $G_{S+1}$ phosphate (labelled as G10 in panels) is described by $\beta_{trans}/\gamma_{g+}$ dihedrals (left panels) or by presence of the 0BPh interaction, i.e. the $G_{S+1}$(C8H8)…$G_{S+1}$(O5') H…O distance below 2.5 Å (green line, panels on the right). The $\chi_{OL3CP}$ + gHBfix19 *ff* version (see Methods in the main text) was used. PDB (2KOC) numbering of nucleotides from 1 to 14 is used in the panels.

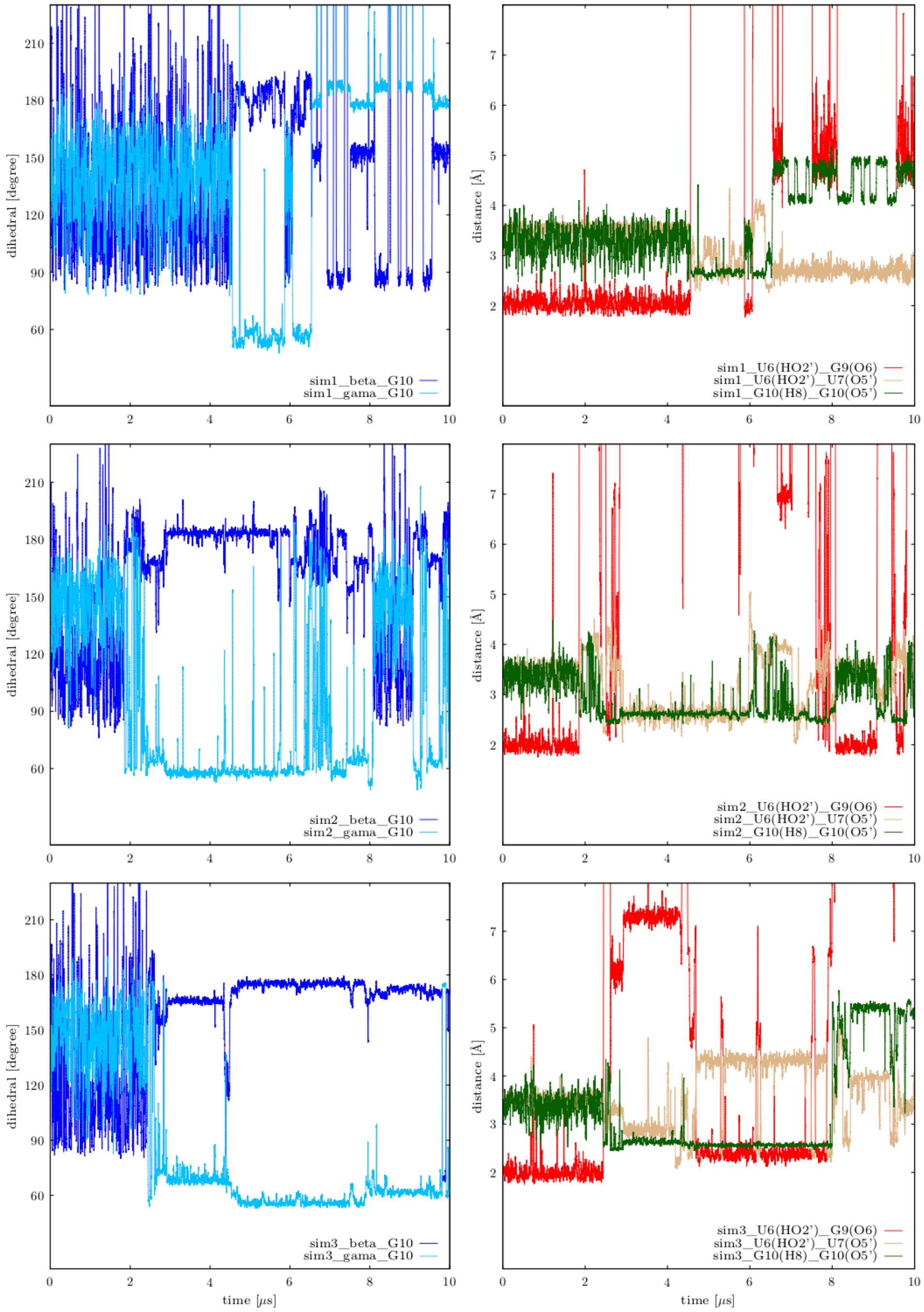

**Figure S3.** Occurrence of the alternative (flipped) G$_{S+1}$ phosphate state in three 10 μs long MD simulations of the r(ggcacUUCGgugcc) 14-mer and possible correlation with the U$_{L2}$(2'-OH) flipping. Native state of the G$_{S+1}$ phosphate is described by $\beta_{trans}/\gamma_{g+}$ dihedrals (left panels) or by presence of the 0BPh interaction, i.e. the G$_{S+1}$(C8H8)…G$_{S+1}$(O5') H…O distance below 2.5 Å (panels on the right). Note that the figure is similar to **Figure S2**, but MD simulations are longer (μs compared to ns time-scale). Thus, disruption of the loop is observed after 4.6 μs for simulation 1 (sim1), ~1.8 μs for simulation 2 (sim2) and ~2.4 μs for simulation 3 (sim3). The $\chi_{OL3CP}$ + gHBfix19 *ff* version was used. PDB (2KOC) numbering of nucleotides from 1 to 14 is used in the panels.

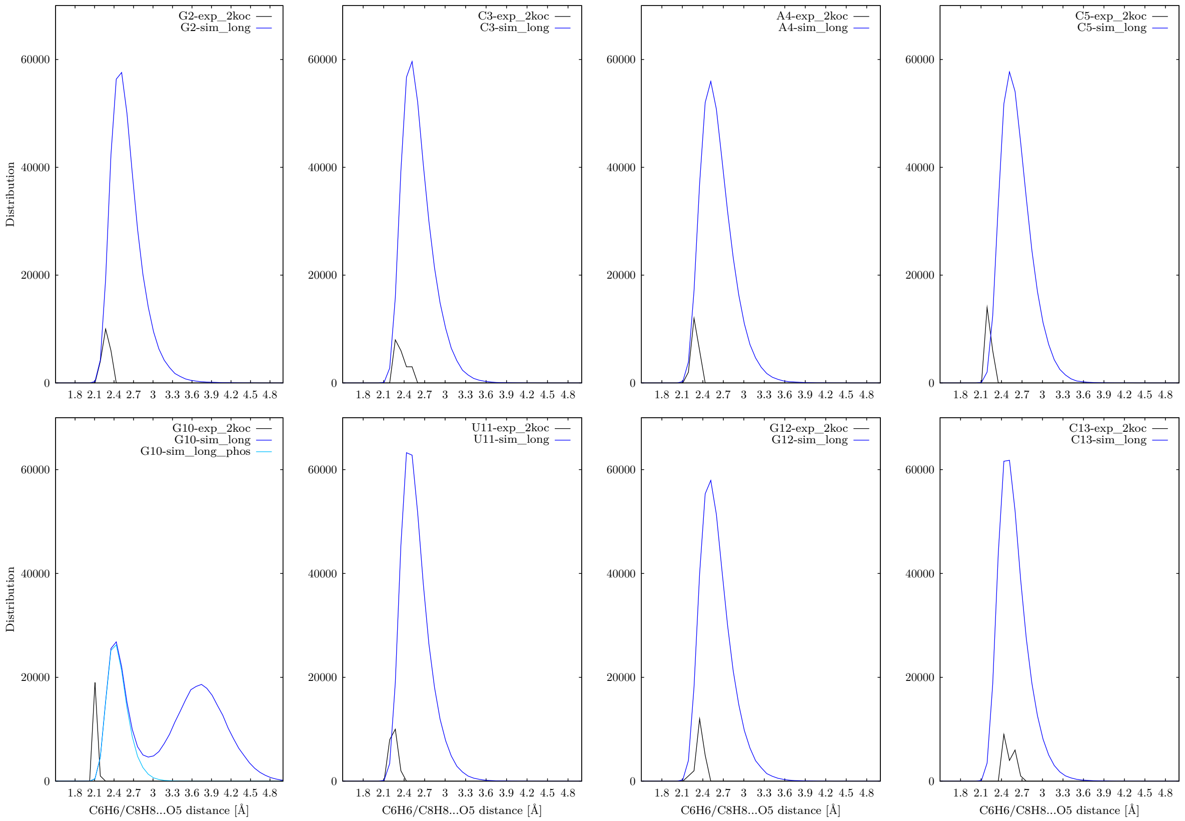

**Figure S4.** Distribution of C8H8/C6H6…O5' distances (0BPh interaction) in MD simulation of the r(ggcacUUCGgugcc) 14-mer using the $\chi_{OL3CP}$ + gHBfix19 *ff* (blue lines). We analyzed only 4.6 μs-long part of MD trajectory before the disruption of the loop occurred, see sim1 in **Figure S3**). There are two states observed for $G_{S+1}$ (first column, second row; $G_{S+1}$ is labelled as G10 in the bottom left panel) corresponding to native and flipped $G_{S+1}$ phosphate, i.e., presence or absence of the 0BPh interaction, respectively. Black lines show distributions in the experimental structure (20 NMR models, PDB ID 2KOC). PDB (2KOC) numbering of nucleotides from 1 to 14 is used in the panels.

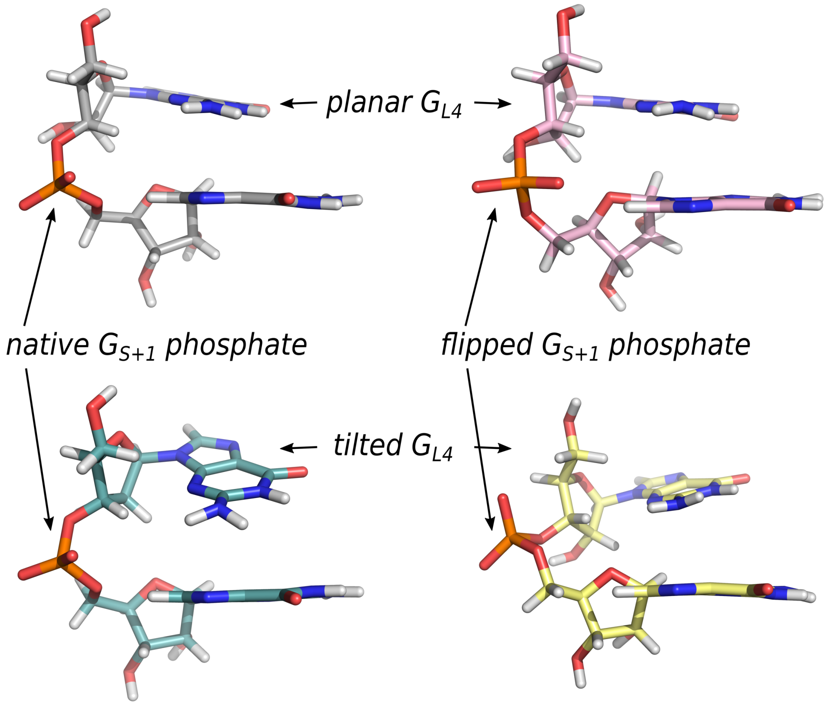

**Figure S5.** The four dinucleotide monophosphate models representing four combinations of the $G_{L4}$ nucleobase (*planar* or *tilted*) and $G_{S+1}$ phosphate (*native* or *flipped*) states.

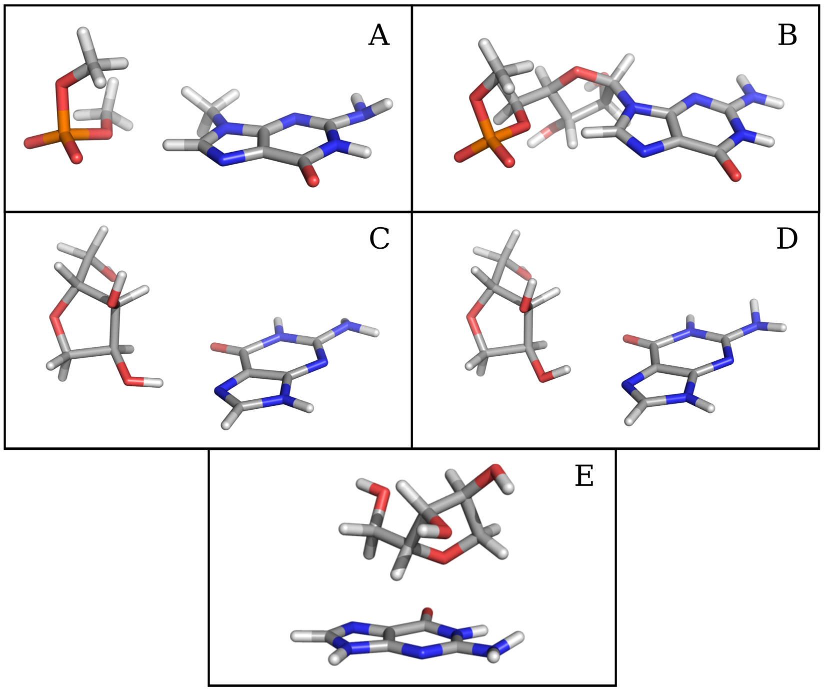

**Figure S6.** Models used for scans describing the 0BPh interaction (first row) dimethyl-phosphate – methylguanine molecular complex (**A**) and O3'-methylated-guanosine-monophosphate unit

(**B**), models used for scans describing $U_{L2}$(2'-OH)…$G_{L4}$(N7) H-bond (second row) with O-H…N angle equal to 170° (**C**) and 143° (**D**), and model used for the scan describing sugar-base stacking (**E**).

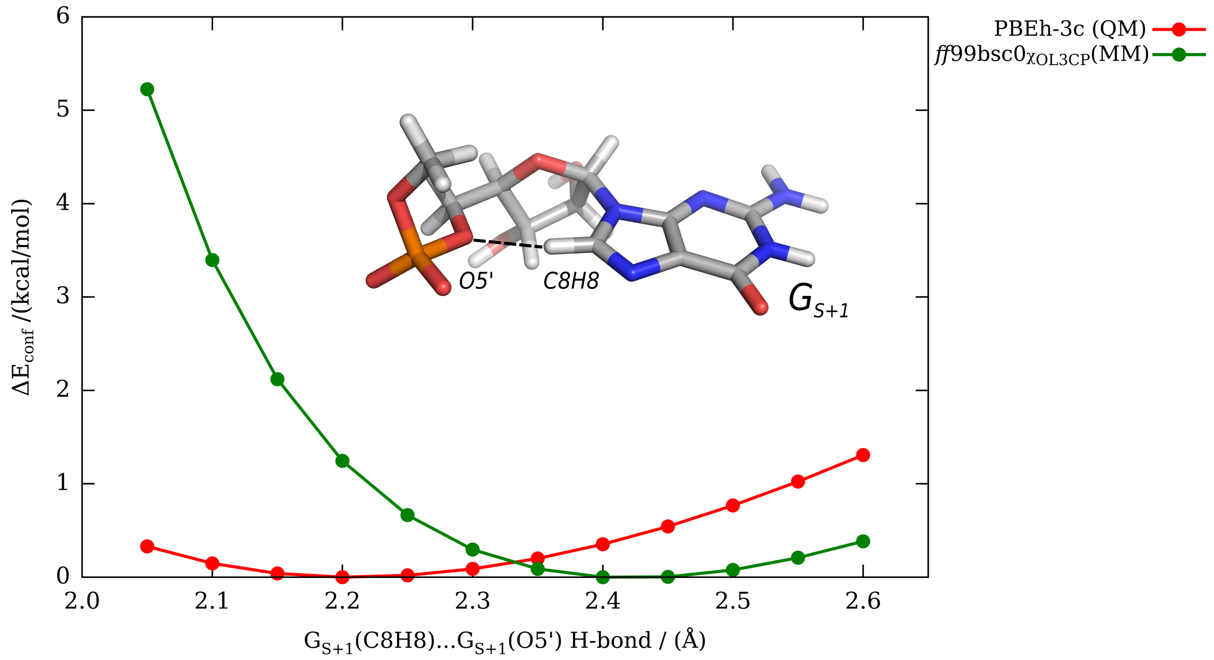

**Figure S7.** Conformational energy scan along the $G_{S+1}$(C8H8)…$G_{S+1}$(O5') distance describing the 0BPh interaction using QM (PBEh-3c; red) and MM (*ff*99bsc0$\chi_{OL3CP}$; green) methods performed on the O3'-methylated-guanosine-monophosphate model.

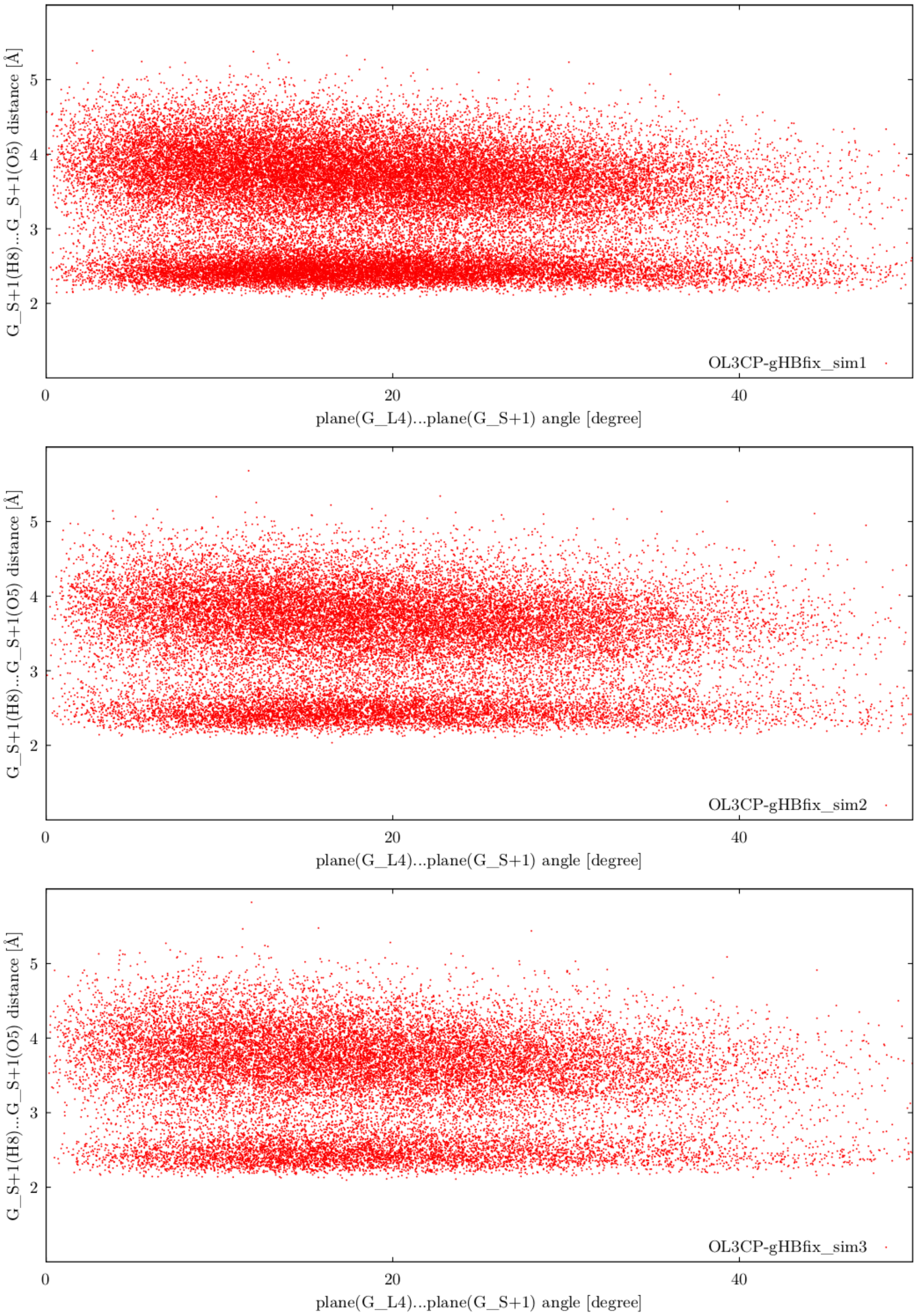

**Figure S8.** Relationship between the 0BPh interaction represented by the $G_{S+1}$(C8H8)…$G_{S+1}$(O5’) H…O distance below 2.5 Å (y-axis) and the $G_{L4}$-$G_{S+1}$ state (*planar* or *tilted*) represented by the angle between planes defined by $G_{L4}$ and $G_{S+1}$ nucleobases (x-axis). Only ~4.6 µs-long (sim1), ~1.8 µs-long (sim2) and ~2.4 µs-long (sim3) parts of three MD trajectories were analyzed (see **Figure S3**).

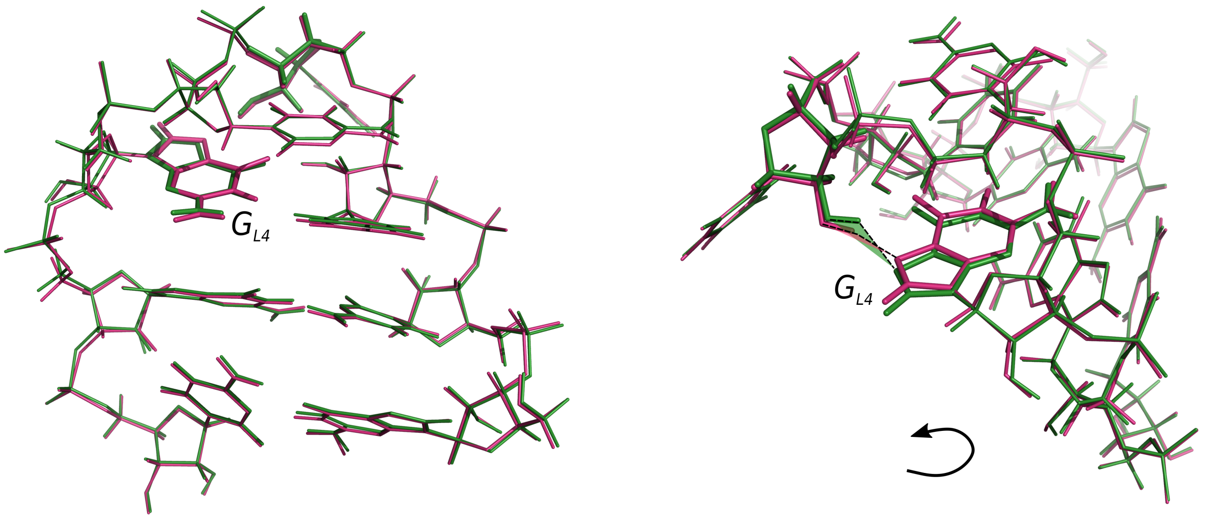

**Figure S9.** UNCG TL structure **6** optimized at QM/MM (PBEh-3c; pink) and MM (*ff*99bsc0$\chi_{OL3}$; green) levels of theory. The Figure shows insufficient description of $U_{L2}$(2'-OH)…$G_{L4}$(N7) signature H-bond by the MM, which is even visually visible on the right part.

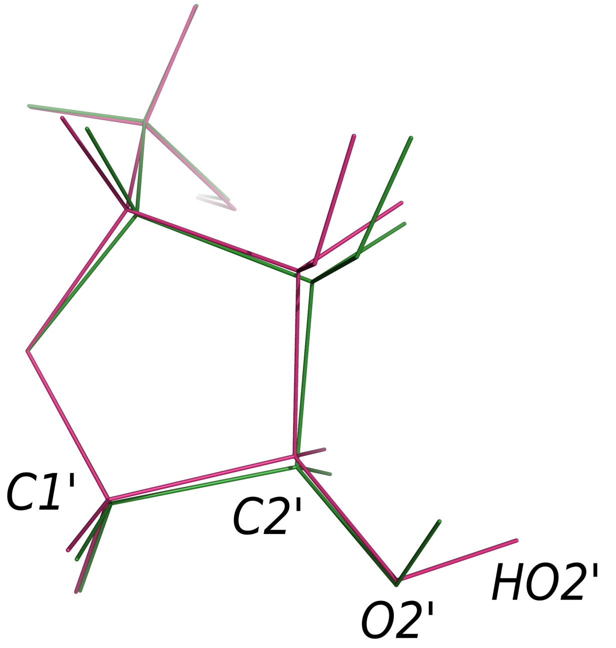

**Figure S10.** Model of the QM (PBEh-3c; pink) and MM (*ff*99bsc0$_{CP}$; green) optimized ribose.

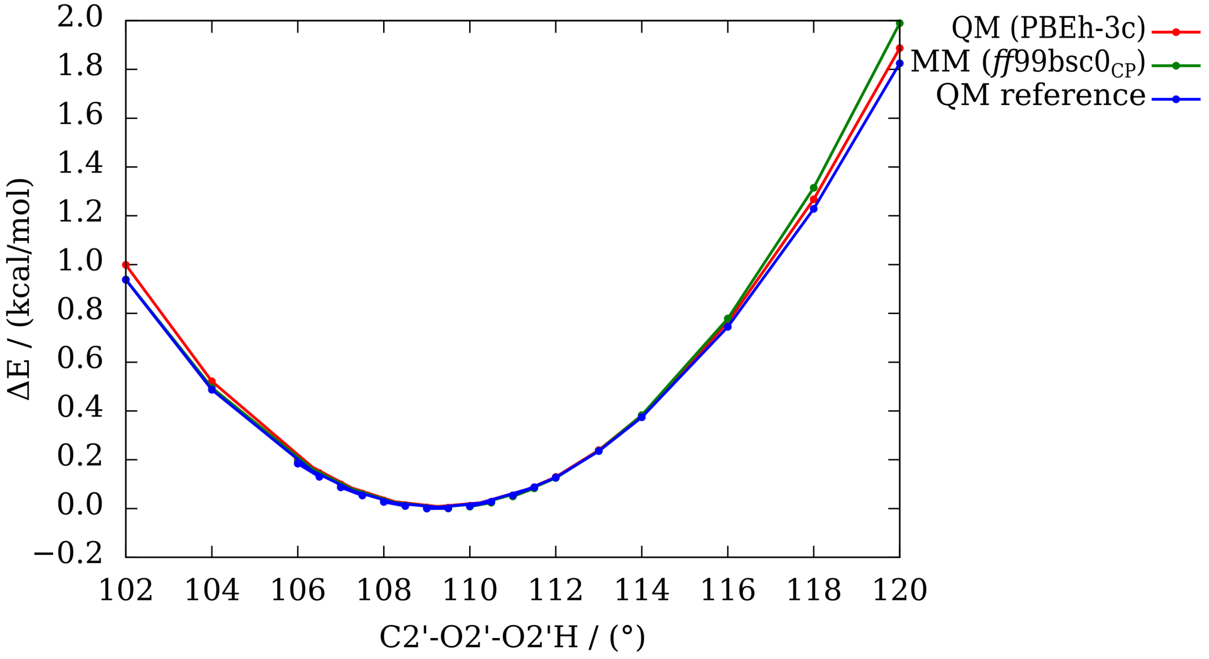

**Figure S11.** Conformational energy profile of the $U_{L2}$ ribose along the C2'-O2'-O2'H angle using QM (PBEh-3c; red), MM (*ff*99bsc0; green) and QM reference (DLPNO-CCSD(T)/CBS; blue) methods. The reference point (zero energy) refers to the C2'-O2'-O2'H angle of the QM-optimized ribose (109.3°).

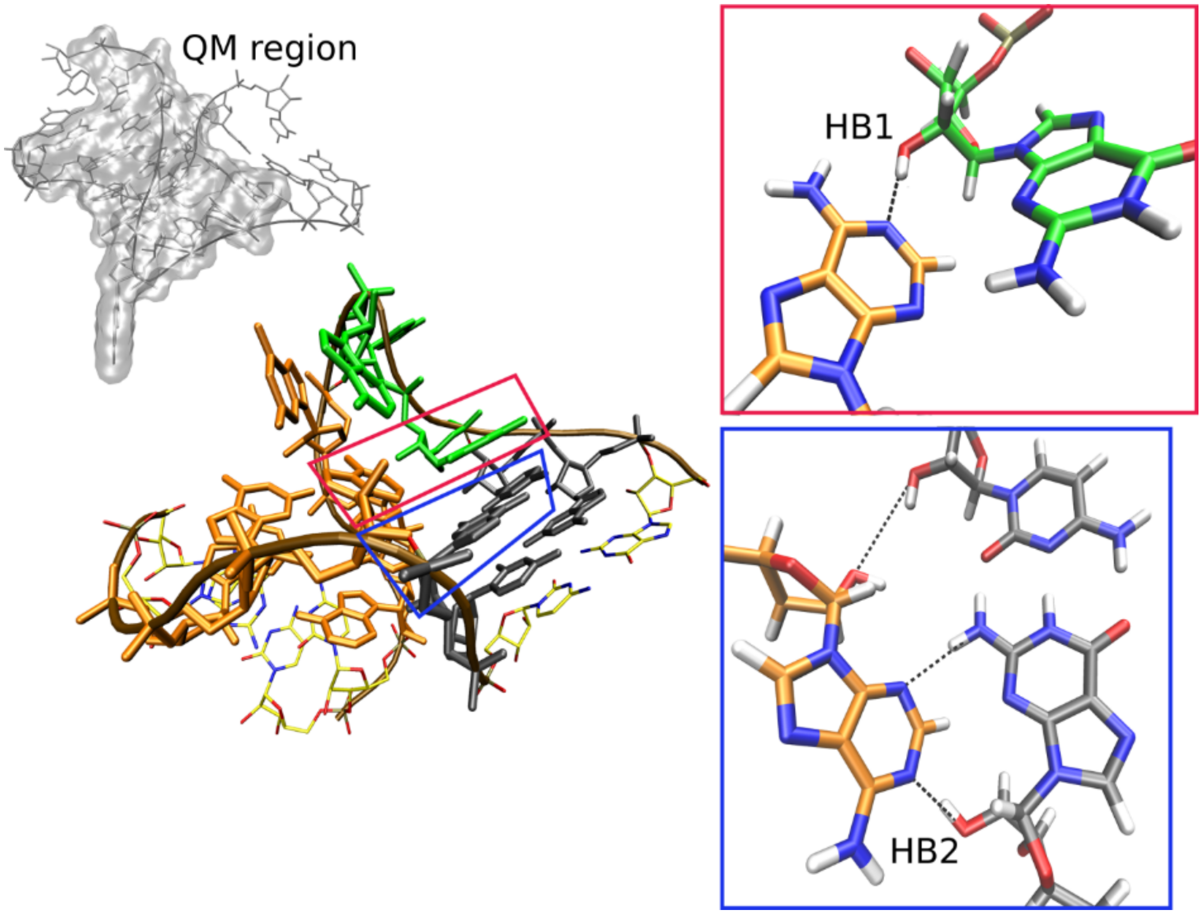

**Figure S12.** 3D representation of the Kt-7 structure, with the canonical and non-canonical stems, and the bulge, in grey, orange, and green, respectively. The RNA backbone is traced in brown. The A-minor interaction and the kink-turn signature interaction are highlighted in blue and red, respectively and their details displayed in insets on the right. The grey Figure upper left depicts segment of the molecule included in QM region of the QM/MM calculations. The dashed black lines (right) indicate H-bonds.

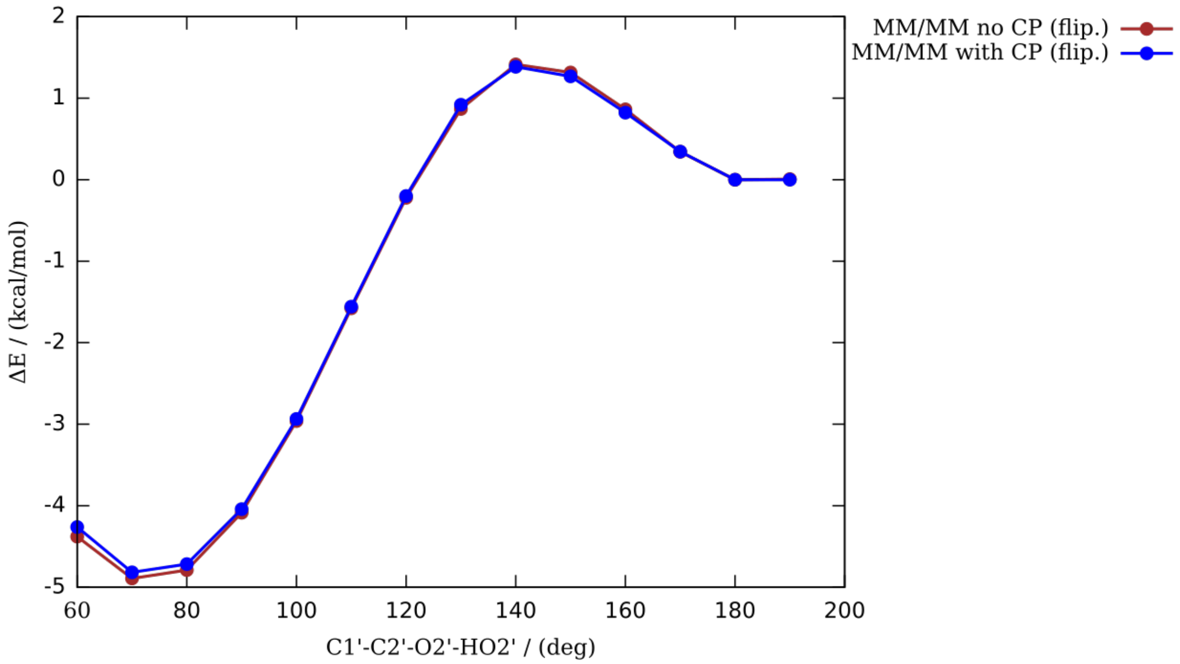

**Figure S13.** MM/MM conformation energy scan for structure **8** with and without the modified vdW parameters for phosphate oxygens (CP).[26]

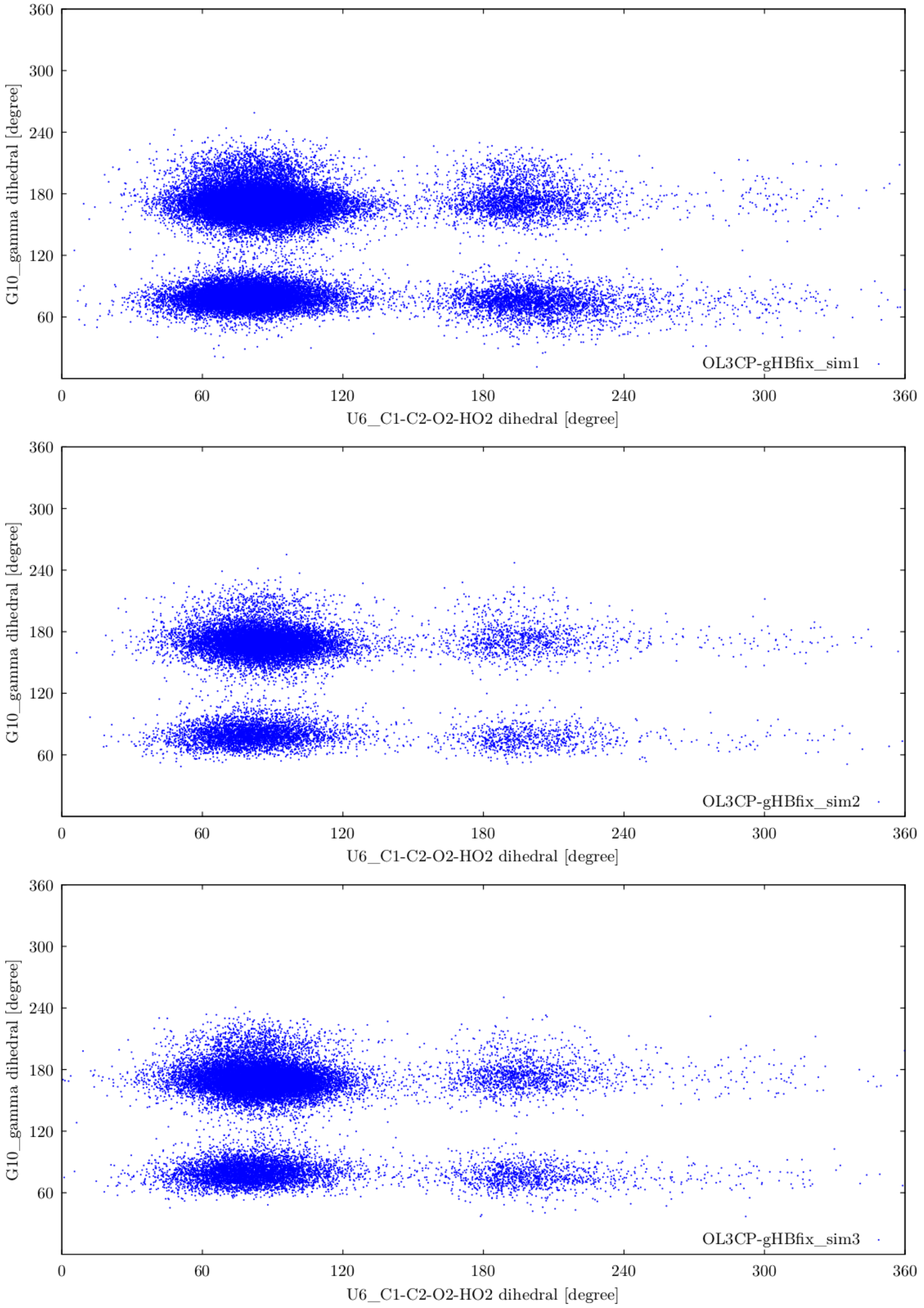

**Figure S14.** Correlation plot of $G_{S+1}$ phosphate conformation, i.e. native state (γ ~60°) or flipped state (γ ~180°), and $U_{L1}$(2'-OH) flipping, i.e. native orientation (C1'-C2'-O2'-O2'H ~80°) or flipped orientation (C1'-C2'-O2'-O2'H ~180°). Note that all four substates, i.e. native/flipped $G_{S+1}$ phosphate combined with native/flipped $U_{L1}$(2'-OH) were populated during MD simulations. $G_{S+1}$ nucleotide is labelled as G10 in panels. Only ~4.6 µs-long (sim1), ~1.8 µs-long (sim2) and ~2.4 µs-long (sim3) parts of three MD trajectories were analyzed (see **Figure S3**).

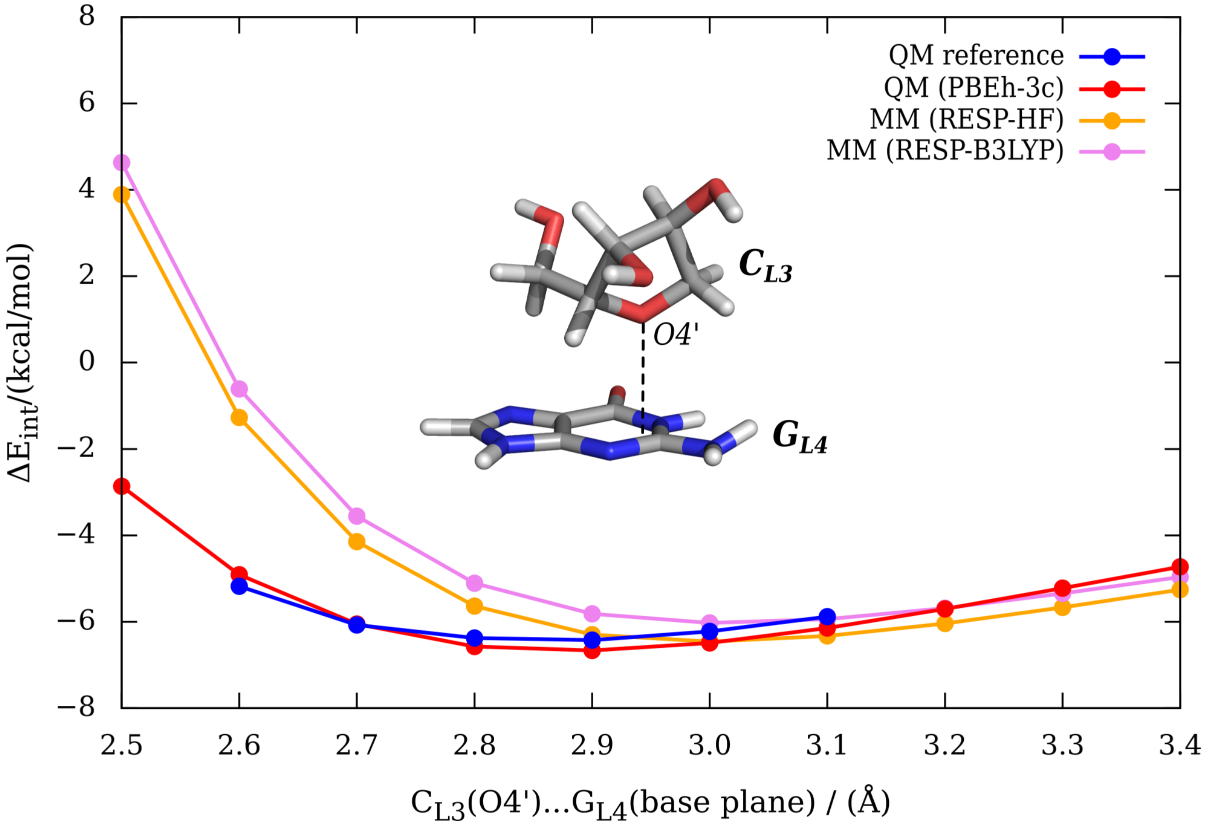

**Figure S15.** Interaction energy scans of the sugar - base stacking between the O4' atom of the $C_{L3}$ ribose and the $G_{L4}$ nucleobase plane. The MM method using two types of partial charges (RESP-HF, RESP-B3LYP; for more information see **Section S3**) is evaluated against the QM methods, i.e. PBEh-3c used in QM/MM optimizations of the UNCG TL and the DLPNO-CCSD(T)/CBS method used as the reference. Optimal distances are 3.00 Å, 3.01 Å, 2.88 Å and 2.87 Å for MM (RESP-HF), MM (RESP-B3LYP), QM (PBEh-3c) and QM reference, respectively, showing that MM overestimates the optimal distance by ~0.13 Å. Interaction energies at the optimal distances are comparable between QM and MM methods, i.e., −6.46 kcal/mol, −6.03 kcal/mol, −6.67 kcal/mol and −6.43 kcal/mol for MM (RESP-HF), MM (RESP-B3LYP), QM (PBEh-3c) and QM reference, respectively. Overestimation of the short-range repulsion by the Lennard-Jones MM is clearly seen.

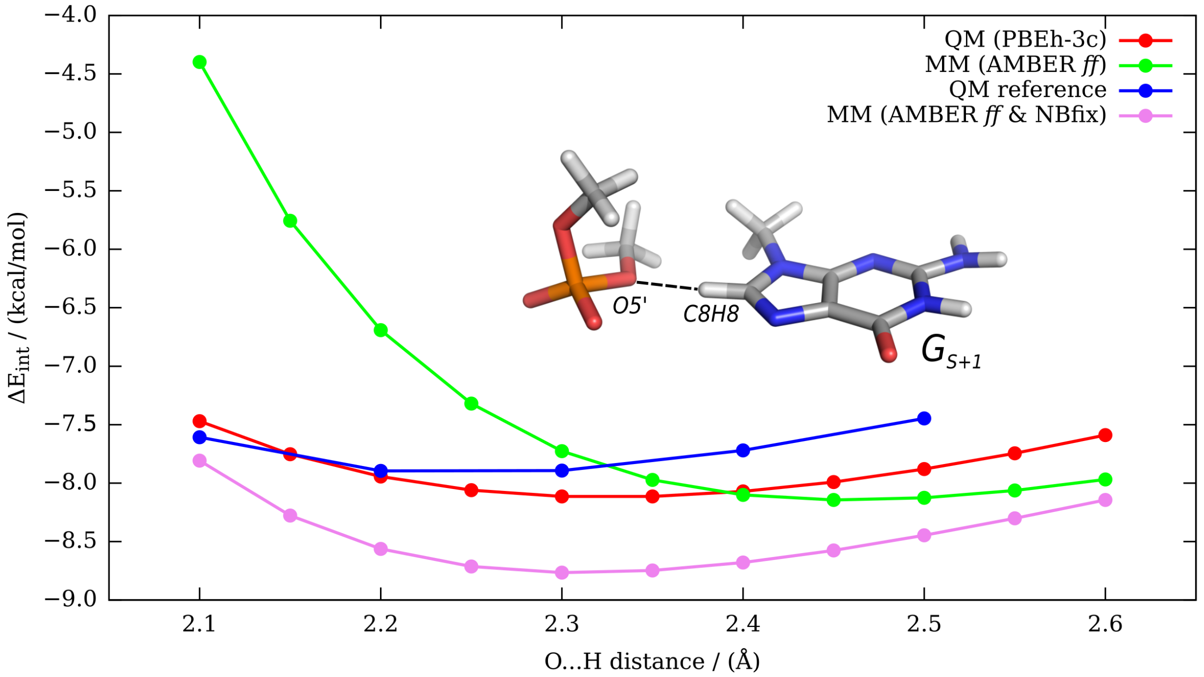

**Figure S16.** Comparison of the QM (PBEh-3c; red), MM (AMBER *ff*; green), QM reference (DLPNO-CCSD(T)/CBS; blue) and MM (AMBER *ff* with NBfix correction; see **Section S3**; purple) interaction energy profiles along the $G_{S+1}$(C8H8)…$G_{S+1}$(O5') H…O distance of the dimethyl-phosphate – methyl-guanine model. Optimal distances are 2.32 Å, 2.46 Å, 2.24 Å and 2.32 Å for QM, MM, QM reference and MM corrected by NBfix, respectively. The overestimation of the optimal $G_{S+1}$(C8H8)…$G_{S+1}$(O5') distance by the original MM potential (green) is corrected by the designed NBfix correction. Graph is extended version of the **Figure 5** in the main text.

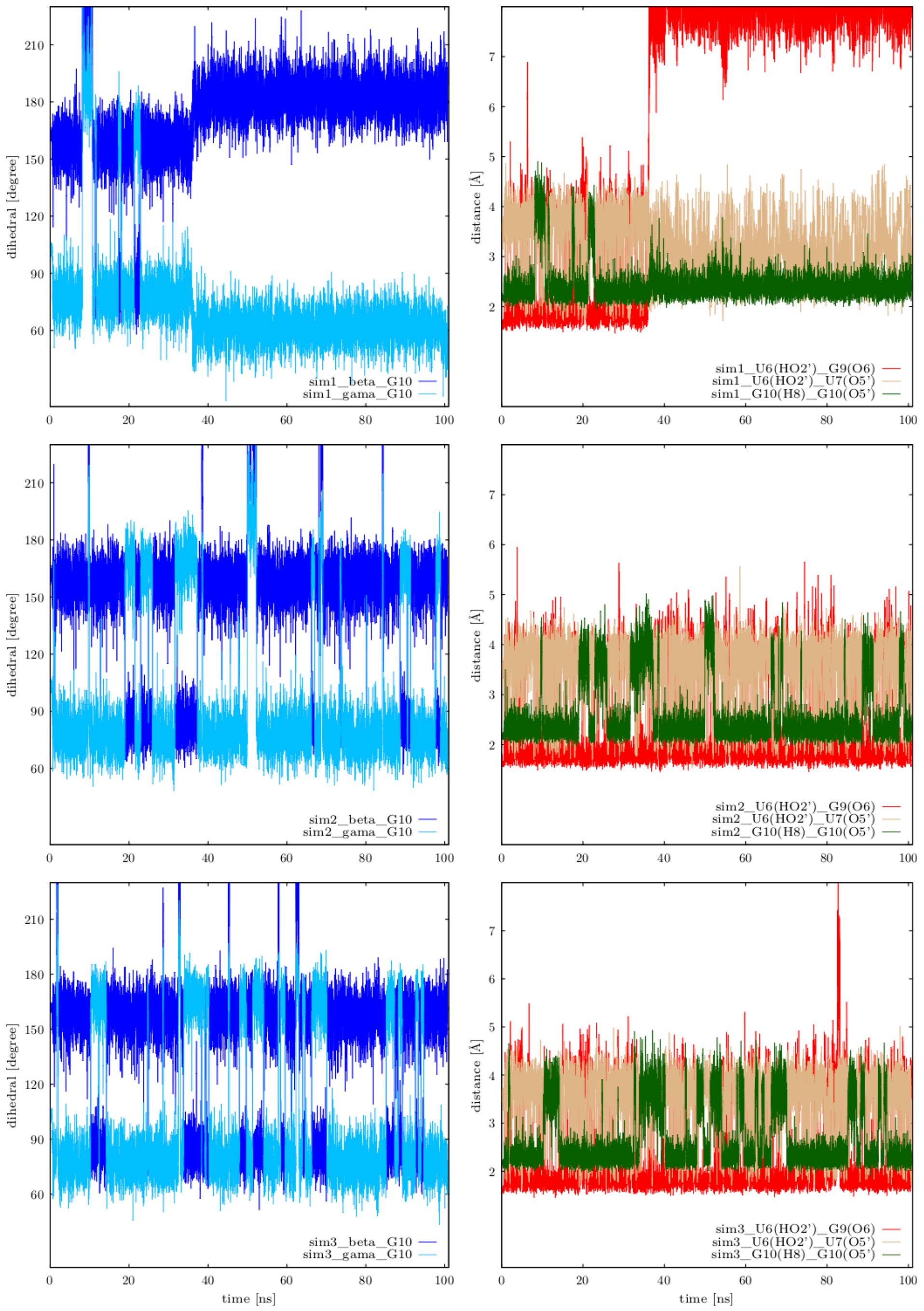

**Figure S17.** Occurrence of the alternative (flipped) G$_{S+1}$ phosphate state in three 100 ns-long MD simulations of the r(ggcacUUCGgugcc) 14-mer and its possible correlation with the U$_{L2}$(2'-OH) flipping. The sum of vdW radii between H8 atoms of all purine bases and O5' oxygens of phosphates was decreased by 0.25 Å in these simulations using NBFix (see the main text). Native state of the G$_{S+1}$ phosphate is described by $\beta_{trans}/\gamma_{g+}$ dihedrals (left panels) and the 0BPh interaction is monitored by the distance between G$_{S+1}$(C8H8) and G$_{S+1}$(O5') atoms, i.e., the 0BPh interaction is present if the G$_{S+1}$(C8H8)…G$_{S+1}$(O5') distance is below 2.5 Å (panels on the right, green). Note that G$_{L4}$ and G$_{S+1}$ are labeled as G9 and G10 in the panels. G$_{L4}$ left its binding pocket just after ~36 ns in simulation 1 (sim1).

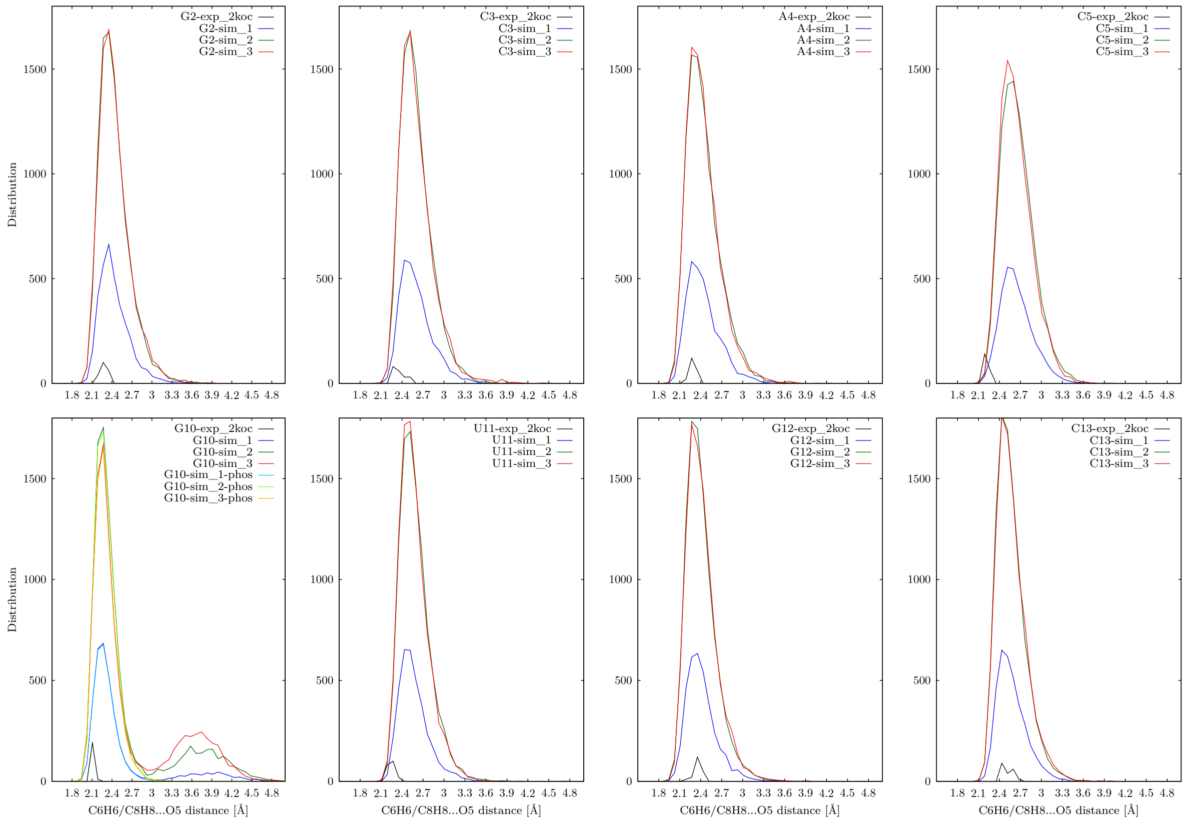

**Figure S18.** Distribution of C8H8/C6H6…O5' distances (0BPh interaction) in three 100 ns-long MD simulations with NBfix correction applied to the –C8H8…O5'– pair of the 0BPh interaction as described in the main text. Note that the NBfix correction is shifting the histogram for $G_{S+1}$ (labeled as G10 in bottom left panel) towards lower values of the $G_{S+1}$(C8H8)…$G_{S+1}$(O5') distance when compared to **Figure S4**. There are two states observed for $G_{S+1}$ (first column, second row; $G_{S+1}$ is labelled as G10 in the bottom left panel) corresponding to native and flipped $G_{S+1}$ phosphate, i.e., presence or absence of the 0BPh interaction, respectively. Distributions considering only snapshots with the native $G_{S+1}$ phosphate conformation are colored as light blue, light green and orange for sim1, sim2 and sim3, respectively (bottom left panel, curves marked as "…-phos"). Black lines show distributions in the experimental structure (20 NMR models, PDB ID 2KOC). PDB (2KOC) numbering of nucleotides from 1 to 14 is used in the panels.

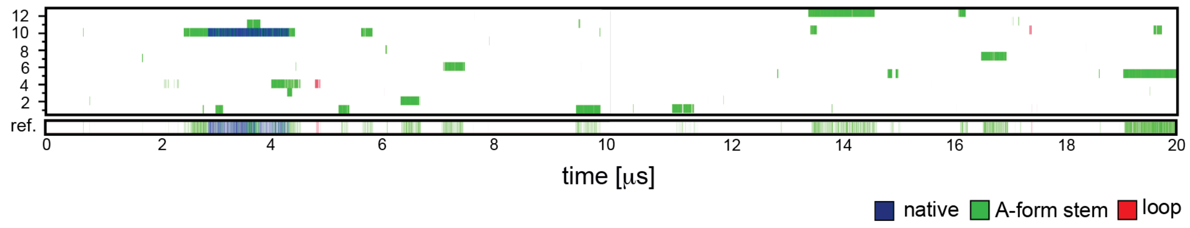

**Figure S19.** Conformational sampling and convergence of REST2 folding simulation of the r(gcUUCGgc) TL with the gHBfix$_{UNCG19}$ correction. Panel shows time evolution of major conformers, i.e., (i) correctly folded A-form stem and loop (native states with all signature interactions formed, blue), (ii) folded A-form stem (loop not in native conformation, green), and (iii) correctly folded loop (stem not in A-form, red), within all twelve continuous (demultiplexed) trajectories and the reference replica.

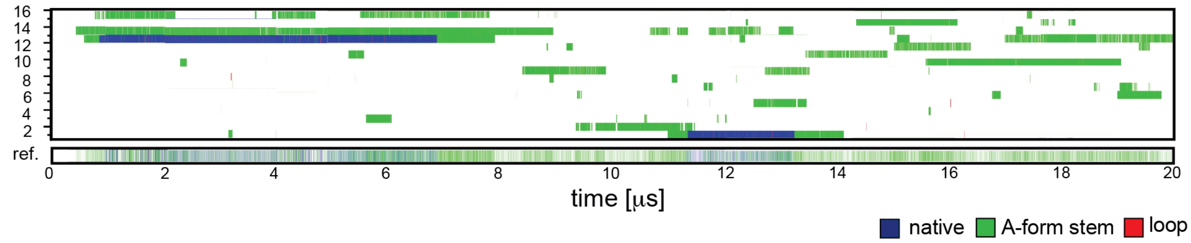

**Figure S20.** Conformational sampling and convergence of REST2 folding simulation of the r(gcUUCGgc) TL with the gHBfix$_{UNCG19}$ correction in combination with NBfix correction applied on the –H8…O5'– pair of the 0BPh interaction. In addition, we reduced vdW radii of all non-polar H atoms (H1, H4, H5 and HA atoms) to 1.2 Å (see the main text). Panel shows time evolution of major conformers, i.e., (i) correctly folded A-form stem and loop (native states with all signature interactions formed, blue), (ii) folded A-form stem (loop not in native conformation, green), and (iii) correctly folded loop (stem not in A-form, red), within all sixteen continuous (demultiplexed) trajectories and the reference replica. Total population of the native state in the reference replica is ~7%, which is still not sufficient in comparison with what is expected from experiments (~25%).[27–29]

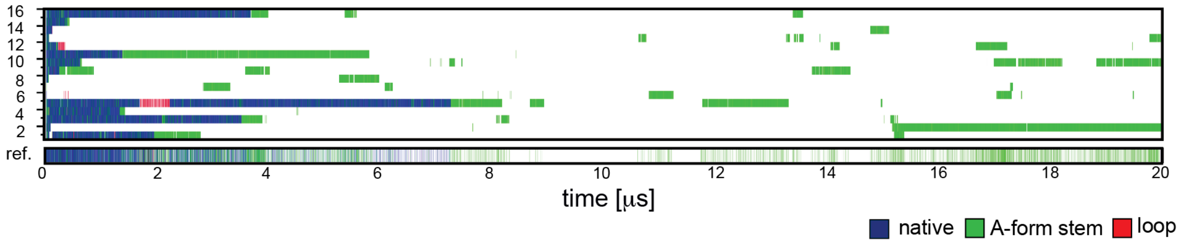

**Figure S21.** Conformational sampling and convergence from REST2 unfolding simulation of the r(gcUUCGgc) TL. All replicas were started from the native state. We applied the $gHBfix_{UNCG19}$ correction in combination with NBfix correction to the –H8…O5'– pair of the 0BPh interaction. We also reduced vdW radii of all non-polar H atoms (H1, H4, H5 and HA atoms) to 1.2 Å and support the native $U_{L1}$(2'-OH)…$G_{L4}$(O6) H-bond by restraining the $U_{L1}$(C1'-C2'-O2'-HO2') dihedrals (see main text). Panel shows time evolution of major conformers, i.e., (i) correctly folded A-form stem and loop (native states with all signature interactions formed, blue), (ii) folded A-form stem (loop not in native conformation, green), and (iii) correctly folded loop (stem not in A-form, red), within all sixteen continuous (demultiplexed) trajectories and the reference replica. The native state was lost in all replicas after ~7.5 μs despite all the restraints and corrections, indicating that the dihedral restraint (which was applied to all nucleotides) rather deteriorated the simulation, cf. with **Figure S20**.
